# CRISPR/Cas9 genome editing to generate single variant *Plasmodium falciparum* lines and enable reverse genetic studies of PfEMP1 function in live parasites

**DOI:** 10.64898/2026.06.07.729090

**Authors:** Stanley E. Otoboh, Hussein M. Abkallo, Jessie Jungels, Nouhoum Diallo, Brian R. Omondi, J. Alexandra Rowe

**Affiliations:** Institute of Immunology and Infection Research, School of Biological Sciences, University of Edinburgh, Edinburgh, UK

## Abstract

Adhesion interactions between Plasmodium falciparum infected erythrocytes (IEs) and human cells bring about microvascular sequestration and contribute to severe malaria pathology. Parasite adhesion molecules on the IE surface are members of the P. falciparum erythrocyte membrane protein 1 (PfEMP1) family, encoded by var genes, which interact with receptors on human cells. Progress in understanding PfEMP1-host receptor interaction is hindered by the lack of genetic tools for PfEMP1 functional studies in live parasites and the spontaneous switching of var gene transcription in culture leading to change in adhesion phenotype. We developed a CRISPR/Cas9 genome editing strategy that takes advantage of var gene mutually exclusive expression to generate single variant P. falciparum lines and enable reverse genetic studies of PfEMP1 function. A drug resistance gene and 2A peptide enabling bi-cistronic transcription were inserted between the promoter and exon I of the it4var60 gene encoding a PfEMP1 variant that mediates the virulence-associated rosetting phenotype. After genome editing and drug selection, only it4var60-transcribing parasites survived, and >90% of IEs expressed IT4VAR60-PfEMP1 on their surface and formed rosettes. When drug pressure was removed, switching to other variants occurred. The approach was adapted to generate epitope tagged-PfEMP1 allowing immunofluorescent detection with commercial antibodies, and modifications of the homology directed repair template enabled investigation of PfEMP1 function including point mutations and a gene knockout that abolished adhesion. These methods can be applied to any var gene in any P. falciparum genotype and are potentially transformative for functional studies of multi-gene family members in live parasites.

## Introduction

Severe malaria caused by Plasmodium falciparum results from extensive sequestration of infected erythrocytes (IEs) in the microvasculature, causing obstruction to blood flow, hypoxia, acidosis and organ dysfunction (Kaul et al., 1991; Miller et al., 2002). Sequestration is mediated by parasite-derived adhesion molecules that are members of variant surface antigen (VSA) families that are expressed on the surface of IEs and bind to human cells (Kyes et al., 2001; Rowe et al., 2009). The best characterised VSA family is P. falciparum erythrocyte membrane protein 1 (PfEMP1), a group of hypervariable proteins of 200 – 450kDa (Leech et al., 1984), encoded by var genes (Baruch et al., 1995; Smith et al., 1995; Su et al., 1995). PfEMP1 proteins are modular and composed of Duffy binding-like (DBL) and cysteine-rich interdomain region (CIDR) domains, which are classified into sequence subtypes such as DBLα, DBLβ, DBLγ, DBLε and CIDRα. These domains are further divided into distinct classes and arranged into conserved domain cassettes (DCs) associated with specific receptor-binding phenotypes and disease outcomes (Rask et al., 2010; Smith et al., 2000).

Each P. falciparum isolate genome contains ∼60 var genes, each encoding an antigenically distinct PfEMP1 variant that binds to one or more host cell receptors. var gene expression occurs in a mutually exclusive manner, with only one transcribed per cell, while the others are silenced (Frank & Deitsch, 2006; Scherf et al., 1998; Voss et al., 2006). After asexual division, daughter cells can switch to transcribe a different var gene in a process known as antigenic variation (Biggs et al., 1991; Dzikowski et al., 2006; Roberts et al., 1992). This allows the parasite to evade clearance by host antibodies and establish long-term infections to enable transmission.

The var gene repertoire of different parasite isolates is largely non-overlapping, resulting in a family of PfEMP1 variants with extreme diversity in the global parasite population (Barry et al., 2007). Despite this amino acid sequence diversity, subsets of PfEMP1 variants from diverse P. falciparum isolates have been characterised that mediate adhesion to various host cells and receptors, of which the PfEMP1 variants that bind to endothelial receptors CD36, ICAM-1 and EPCR are the most well-studied (Hsieh et al., 2016; Lau et al., 2015; Lennartz et al., 2017; Lennartz et al., 2019). Current knowledge of PfEMP1 adhesion functions is primarily derived from evidence using recombinant proteins expressed in heterologous systems (Angeletti et al., 2015; Avril et al., 2016; Avril et al., 2012; Ghumra et al., 2012; Higgins, 2008; Juillerat et al., 2011; Rowe et al., 1997; Vigan-Womas et al., 2012). Such studies have provided ground-breaking insights into PfEMP1 structure and function (Hsieh et al., 2016; Lau et al., 2015; Lennartz et al., 2017; Lennartz et al., 2019) but do have limitations. For example, post-translational modifications of recombinant proteins may not match the native protein (Crosnier et al., 2013), single PfEMP1 domain recombinant proteins have often been used, so features depending on tertiary or quaternary structure of the protein (Rajan Raghavan et al., 2023) could be missed, and the properties of recombinant proteins do not always match those of the native protein (Azasi et al., 2018).

Studies of PfEMP1 function in live parasites are needed to compensate for the above limitations, but this has proved challenging. Culture-adapted parasite lines can be enriched for specific adhesion phenotypes by panning or density gradient techniques (Claessens et al., 2012; Handunnetti et al., 1992) and parasites expressing specific PfEMP1 variants can be selected using variant-specific antibodies (Dalgaard et al., 2025; Guillotte et al., 2016; Joergensen et al., 2010; Moll et al., 2007). These methods are laborious and inefficient, and selected parasite lines rapidly lose their phenotype due to spontaneous var gene switching in culture. This spontaneous switching away from the selected PfEMP1 to other variants during every 48-hour asexual cycle in culture, coupled with the low efficiency of genetic manipulation in P. falciparum (Hasenkamp et al., 2012; O’Donnell et al., 2002) has meant that reverse genetic studies of PfEMP1 function have rarely been attempted. Establishing a clear link between the genotype of a modified var gene and an observable PfEMP1 phenotype is hindered when the transgenic parasites switch during phenotyping. All prior reports of var gene knock-out (Bryant et al., 2017; Duffy et al., 2006; Viebig et al., 2007) or targeted mutation (Dorin-Semblat et al., 2019), mostly focussing on the rare strain-transcending var2csa gene, described significant difficulties achieving the desired genetic modifications.

A better method for genetic studies of PfEMP1 function in live parasites would have great potential to provide novel insights into malaria host-parasite interactions. We developed a strategy stimulated by previous work showing that replacement of exon I of a specific var gene with a drug resistance gene and application of drug pressure resulted in the only active var gene promoter being the one driving the drug resistance gene (Dzikowski et al., 2006; Voss et al., 2006). The rest of the var gene family was effectively silenced due to the mutually exclusive expression system (Dzikowski et al., 2006; Dzikowski et al., 2007; Voss et al., 2006). Building on this observation, we hypothesised that using Clustered Regularly Interspaced Short Palindromic Repeats/CRISPR-associated protein 9 (CRISPR/Cas9) genome editing (Ghorbal et al., 2014; Wagner et al., 2014; Zhang et al., 2014) to insert a drug resistance gene and 2A ribosomal “skip” peptide (de Felipe et al., 2006) upstream of a var gene of interest, would allow both genes to be transcribed from the var gene promoter (Fig. 1a). Under drug selection, only genetically modified parasites transcribing the var gene of interest and the drug resistance gene would survive, resulting in a homogenous single PfEMP1-expressing IE population. Wild-type parasites, or those that switched to transcribe other var genes would be killed by the drug. For functional studies, the skip peptide would ensure that the drug resistance protein is not attached to the targeted PfEMP1 variant but is co-expressed as a separate protein. This strategy should allow generation of single variant P. falciparum lines that maintain a stable adhesion phenotype in long-term culture under drug pressure and simultaneously enable reverse genetic studies of PfEMP1 if targeted mutations are included in the CRISPR/Cas9 homology directed repair (HDR) template.

**Figure 1:**
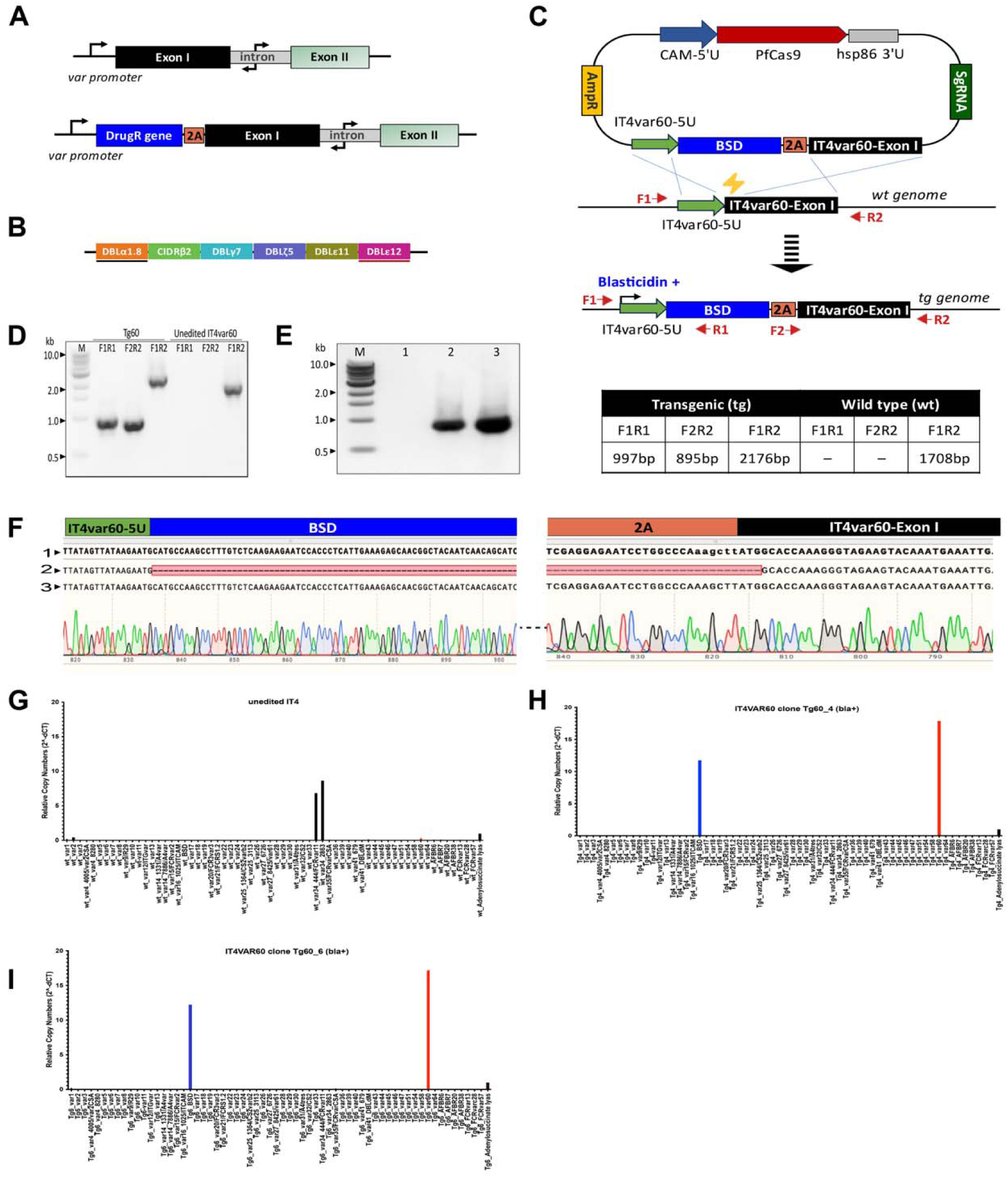
CRISPR-Cas9 genome editing strategy to generate a single variant P. falciparum line. (A) Top: Organisation of a typical var gene, comprising exon I, intron and exon II. Bottom: Strategy to select for expression of a defined var locus by inserting a drug-resistance marker linked via a 2A skip peptide between the endogenous var promoter and exon I, thereby coupling drug selection to transcription of the targeted var gene. (B) Domain architecture of IT4VAR60 PfEMP1. The erythrocyte-binding (black bar) and IgM-Fc binding (red bar) regions are indicated. (C) CRISPR-Cas9-mediated homology directed repair of the it4var60 locus. A Cas9-induced double-strand break is repaired using a donor template containing left (IT4var60-5U) and right (IT4var60-Exon I) homology arms flanking the BSD-2A cassette. Blasticidin pressure selects for parasites with the correct integration. The integration PCR primer positions (F1, R1, F2, R2) in the unedited and edited it4var60 loci are indicated by red arrows. Edited/transgenic parasites yield amplicons of 997 bp, corresponding to 5’ recombination (F1R1); 895 bp, 3’ recombination (F2R2); and 2176 bp (F1R2) respectively. Unedited wildtype parasites yield the native 1708 bp product (F1R2) only. (D) Integration PCR of genomic DNA from Tg60 bulk culture confirming expected amplicon sizes. Unedited IT4var60 parasites served as a control. M, 1kb ladder. (E) Integration PCR of Tg60 clones D11 (Lane 2) and E7 (Lane 3) using F2R2 primers, yielding the expected 895 bp product; no amplification was observed in unedited IT4var60 control (Lane 1). (F) Representative Sanger sequencing chromatogram confirming correct insertion of the BSD-2A cassette at the it4var60 locus (1= plasmid reference; 2=unedited/wildtype IT4; 3=edited Tg60 clone). The BSD-2A sequence is truncated (---) for display. (G) qRT-PCR profiling of var transcription in unedited IT4 parasites showing low level it4var60 (red) transcription and predominant expression of it4var34 compared to the adenylosuccinate lyase (adsl) house-keeping gene. (H-I) qRT-PCR profiling of Tg60 parasites showing it4var60 (red) as the only highly transcribed var gene; bsd (blue) is also highly expressed, normalised to adsl.

A conceptually similar drug selection-linked integration (SLI) approach using traditional homologous recombination techniques was described previously (Birnbaum et al., 2017), and was recently applied to var genes by inserting a drug resistance gene at the 3’ end of a var gene to select parasites expressing a defined PfEMP1 variant (Cronshagen et al., 2025). While this system enabled generation of enriched single-variant parasite lines under drug pressure and facilitated studies of PfEMP1 trafficking and localisation, a key limitation of this innovative approach is that additional rounds of genetic modification would be required after the original SLI to enable reverse genetic experiments of PfEMP1 function.

Here, we present a CRISPR/Cas9 var gene editing strategy that integrates selection and locus-specific modification in a single step by inserting a drug resistance gene and 2A peptide at the var gene 5’ end, enabling simultaneous generation of single-variant parasite lines and reverse genetic modification, including epitope tagging for PfEMP1 detection, gene disruption and targeted point mutagenesis to examine the role of PfEMP1 in adhesion to human cells.

## Results

### Development of a CRISPR/Cas9 genome-editing strategy to generate P. falciparum lines expressing a single var gene under drug pressure

To implement the single variant strategy, we targeted a var gene encoding the IT4VAR60 PfEMP1 (PfIT_120060300) (Fig. 1b), whose N-terminal DBLα domain binds uninfected erythrocytes to form rosettes (Angeletti et al., 2012; Ghumra et al., 2012; McLean et al., 2025) and C-terminal DBLε12 domain binds IgM-Fc for immune evasion (Akhouri et al., 2016). Rosetting is a phenotypically variable IE adhesion property that occurs commonly in parasite isolates from severe malaria patients (Doumbo et al., 2009) and contributes to microvascular congestion and pathology (Kaul et al., 1991). Rosetting is difficult to study in vitro due to rapid switching away from the group A subset of var genes (Peters et al., 2007) that encode rosette-mediating PfEMP1 variants (Ghumra et al., 2012; McLean et al., 2025). Hence, rosetting parasite lines require frequent and inefficient density-gradient selection of IEs to maintain the phenotype (Handunnetti et al., 1992). IT4VAR60 was chosen because the NTS-DBLα domain that binds to uninfected erythrocytes is well-characterised (Angeletti et al., 2015), variant-specific antibodies were available to examine IE surface expression (Ghumra et al., 2012), and successful editing could be indicated microscopically by the presence of rosettes in transfected cultures. A single CRISPR/Cas9 genome editing plasmid was designed with an it4var60-specific gRNA sequence and HDR template to incorporate the drug resistance gene (blasticidin S deaminase, bsd) and 2A peptide at the 5’ end of the it4var60 coding sequence, under the control of the var gene promoter (Fig. 1c). The it4var60 promoter sequence in the plasmid was truncated, so that bsd would only be transcribed when incorporated into the genome, and not directly from the plasmid itself (Dzikowski et al., 2006). After transfection and drug pressure, transgenic parasites (hereafter called Tg60) appeared after 10-14 days. Genomic DNA was extracted and integration PCR with specific primers within and outside the transgene (Fig. 1c) confirmed the correct edit (Fig. 1d). Transgenic parasite clones were obtained by limiting dilution cloning (Kirkman et al., 1996), and the bsd-2A insertion was verified by PCR (Fig. 1e) and Sanger sequencing (Fig. 1f).

Quantitative reverse transcription PCR (qRT-PCR) profiling of var gene expression showed that unedited IT4 parasites predominantly expressed the group C gene it4var34 (Fig. 1g), whereas the it4var60 gene was the only major var gene transcribed by Tg60 parasites, and transcription of bsd was also detected (Fig. 1h,i). These data demonstrate that blasticidin S effectively eliminates parasites transcribing alternative var genes, restricting survival to those transcribing it4var60 and yielding a homogenous population of IEs transcribing a single var gene.

### Single var gene-expressing parasites display PfEMP1 on the infected erythrocyte surface

To confirm that the IT4VAR60 PfEMP1 variant was expressed on the surface of live IEs in the Tg60 parasite clones, we used immunofluorescence staining and flow cytometry with polyclonal rabbit IgG against the IT4VAR60 NTS-DBLa domain (Ghumra et al., 2012). 94% of Tg60 mature IEs were recognised by the IT4VAR60 NTS-DBLa antibody, whereas unedited IT4var60 parasites selected for rosetting by traditional density gradient methods showed positive staining in 67% of mature IEs (Fig. 2a). The expression of the whole extracellular region of the IT4VAR60 PfEMP1 variant in Tg60 parasites was confirmed by staining with antibodies specific for other IT4VAR60 domains, including the C-terminal DBLε12 domain (Fig. 2b). Furthermore, non-immune IgM from human serum bound to the surface of live Tg60 IEs as expected (Fig. 2c) (Akhouri et al., 2016). These results confirm expression of the IT4VAR60 PfEMP1 on the IE surface of edited parasites. A small population of PfEMP1-negative IEs (∼5-15%) were seen in all experiments (Fig. 2a,b). Fluorescence-activated cell sorting was used in an attempt to select this population for further study, but they repeatedly failed to grow, raising the possibility that this population represents IEs that have switched to transcribe a different var gene and are dying due to blasticidin S exposure. The rosette frequency (RF, percentage of mature IEs binding 2 or more uninfected erythrocytes) of the Tg60 parasite clones was assessed weekly over the course of one month under drug pressure and ranged between 80% and 92% (Fig. 2d,e). This was significantly higher than the 52-63% RF seen in the unedited IT4var60 parasites selected for rosetting by traditional methods (Fig. 2d,e; clone D11 vs WT, p<0.0001; clone E7 vs WT, p<0.001, one way ANOVA with Tukey’s multiple comparisons test). The rosetting values in Tg60 and unedited IT4var60 control parasite lines mirrored the percentage of IEs staining positively for IT4VAR60 PfEMP1 but tended to be slightly lower. This is expected due to the presence of some IT4VAR60-expressing IEs that only bind a single erythrocyte, which do not fulfil the accepted definition of rosetting (two or more bound erythrocytes) and therefore are not counted as rosettes (Udomsangpetch et al., 1989).

**Figure 2.**
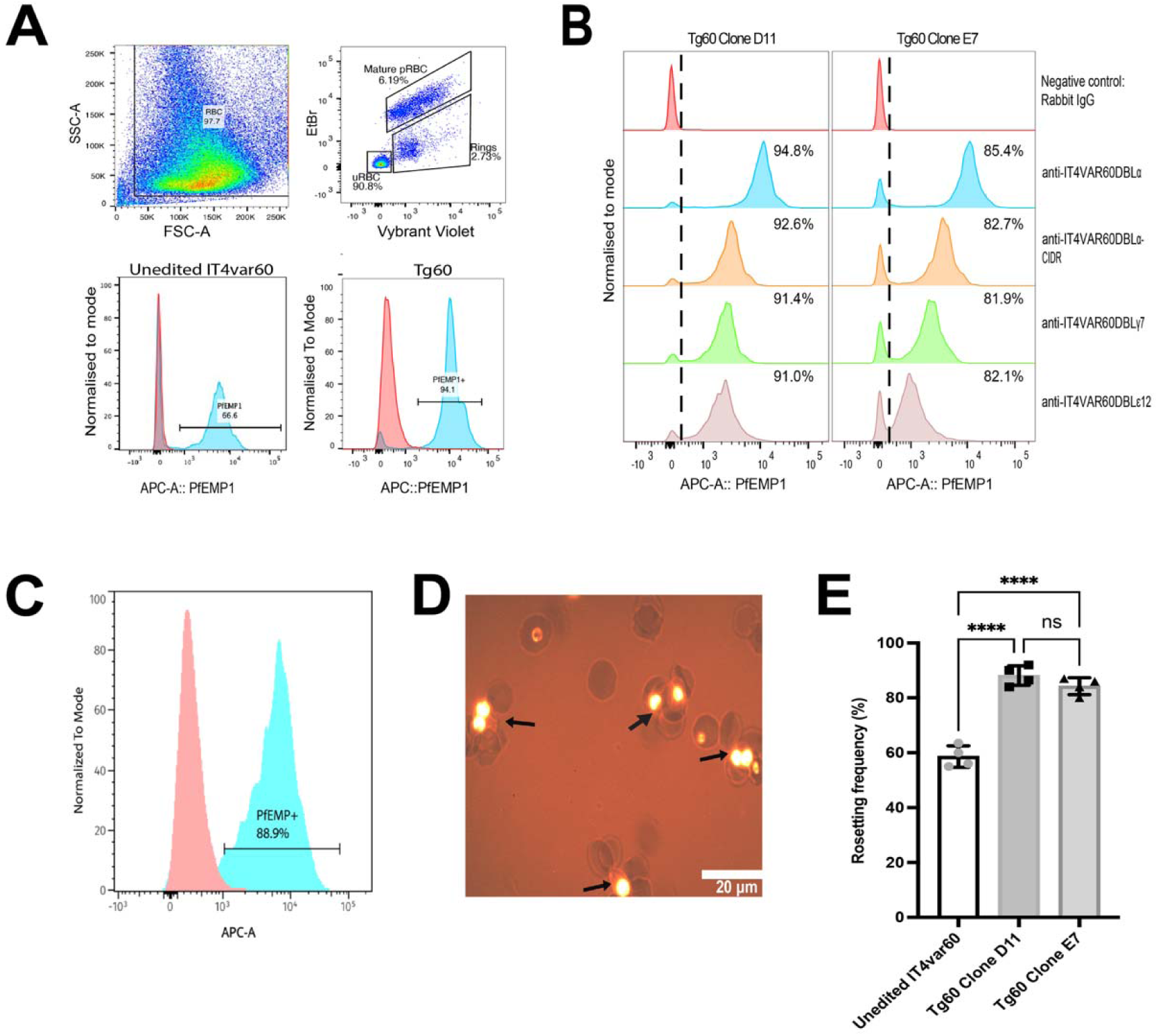
Infected erythrocyte (IE) surface expression of PfEMP1 and rosetting in Tg60 parasites. (A) Flow cytometry gating (upper panels) and surface staining of live pigmented trophozoite and schizont (mature IEs) population with polyclonal anti-IT4VAR60 DBLα antibodies (lower panels). Nucleic acid staining was performed with Vybrant DyeCyle Violet (DNA stain) and ethidium bromide (both DNA and RNA). Mature IEs are distinguished from ring stages by increased RNA content. Histograms show mature IEs stained with non-immunised rabbit IgG (red) or anti-IT4VAR60DBLα IgG (blue). (B) Surface staining of two Tg60 clones (D11 and E7) with polyclonal antibodies to various IT4VAR60 domains (DBLα, DBLα-CIDR, DBLγ7 and DBLε12). Domain recognition ranged from 91.0 – 94.8% in D11 and 82.1 – 85.4% in E7. Flow cytometry images are representative of at least two independent experiments. (C) Surface staining of Tg60 mature live IEs with an antibody to human IgM (blue) or isotype control (red). (D) Representative image of rosettes (black arrows) in a Tg60 clone, stained with ethidium bromide and visualised by fluorescence/bright-field microscopy (400x) on a Leica DM2000 microscope with a Yenway CMOS 5mpx camera. (E) Rosette frequency (percentage of mature IEs binding 2 or more uninfected erythrocytes) measured weekly over 1 month under drug selection. Mean and standard deviation are shown. (**** p<0.0001, one way ANOVA with Tukey’s multiple comparisons test).

To show that the single variant selection approach can potentially be applied to any var gene in any P. falciparum genome, we made two additional parasite lines expressing PfEMP1 types associated with severe malaria: the variants IT4VAR19 (PfIT_010005000, a group B PfEMP1 of the DC8 type) and HB3VAR03 (PfHB3_130080100, a group A PfEMP1 of the DC13 type) (Avril et al., 2012; Claessens et al., 2012; Lau et al., 2015; Lennartz et al., 2017; Turner et al., 2013). In each case, the incorporation of the bsd-2A cassette into the targeted var gene was confirmed by integration PCR (Supplementary Fig. 1,b) and sequencing, and expression of the IT4VAR19 and HB3VAR03 PfEMP1 variants on the surface of live IEs was shown by flow cytometry with variant-specific antibodies (Supplementary Fig. 1c,d). The IT4VAR19-expressing line was panned once (P1) on a human brain endothelial cell (HBEC) line and this increased the level of surface-positive IEs (Supplementary Fig. 1c), whereas >85% of IEs showed surface expression of HB3VAR03 in the bulk culture after transfection without additional selection (Supplementary Fig. 1d). The TgIT4var19 parasite line showed cytoadhesion to HBEC via EPCR as expected (Supplementary Fig. 1e) (Claessens et al., 2012)(Azasi et al., 2018).

### The adhesion phenotype in edited parasites is reversible in the absence/presence of drug

To examine the reversibility of selection in edited parasites, six Tg60 parasite clones were grown for one month without drug pressure, by which time the rosette frequency in the clones varied between 23 and 92% (clone 1, 87%; clone 2, 89%; clone 3, 68%; clone 4, 23%; clone 5, 45%; clone 6, 92%). The two clones with the highest and lowest rosetting were grown for a further 6 weeks, each in the presence or absence of drug. The clone with low rosetting continued to show ∼20% of IE in rosettes in the absence of blasticidin S but reverted to a high rosette frequency (∼85%) within 5 asexual cycles under drug pressure (Fig. 3). Whereas the high rosetting clone maintained its ∼85-90% rosette frequency in the presence of drug, but rosetting decreased gradually overtime in the absence of blasticidin S (Fig. 3). These data show the phenotypic reversibility of the single variant approach, in allowing switching and loss of the adhesion phenotype of interest in the absence of drug, while quickly regaining the adhesion phenotype by re-application of drug pressure.

**Figure 3.**
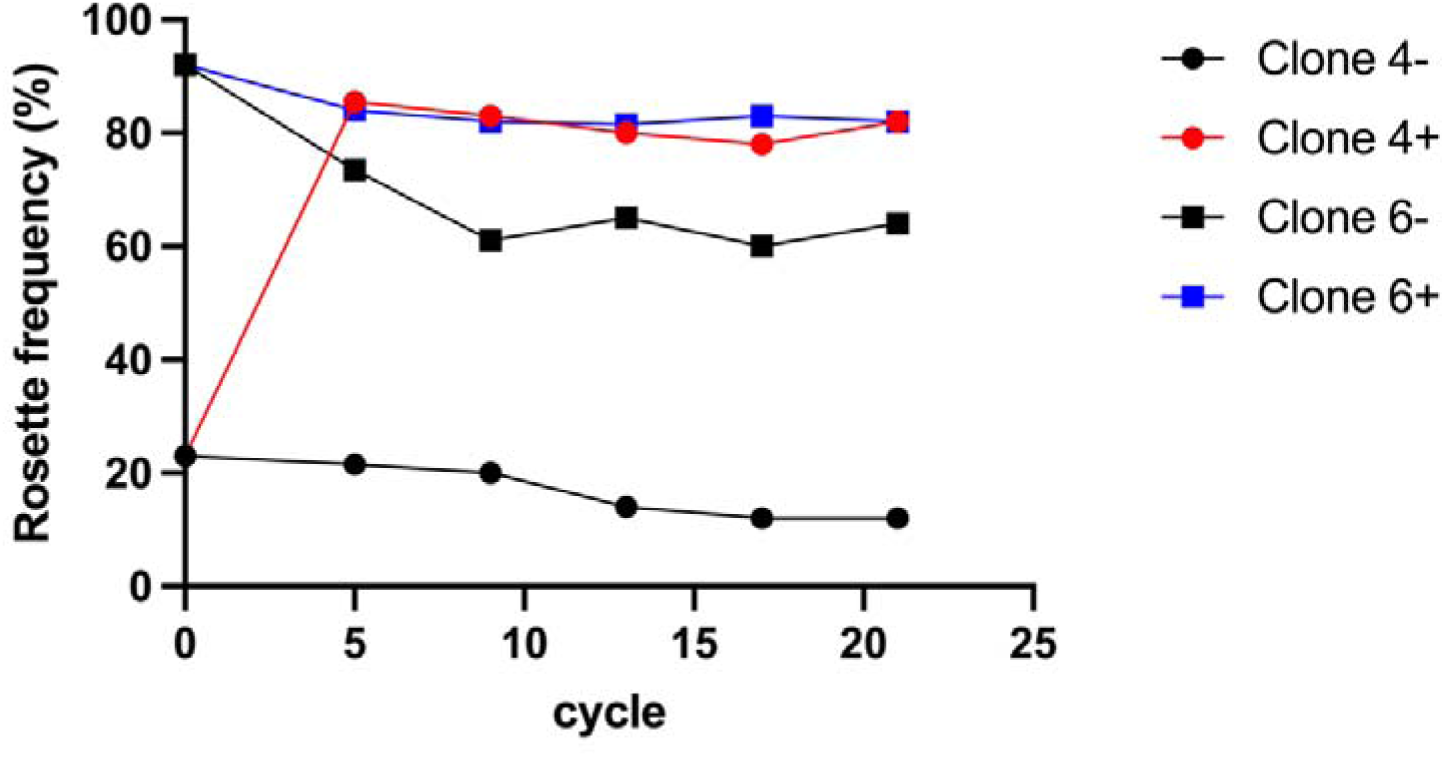
Reversibility of single variant selection. Two Tg60 clones with high (clone 6) or low (clone 4) rosette frequency after a month without drug were grown for a further 21 asexual cycles with (+) or without (-) blasticidin S. The rosette frequency was assessed at cycle 5 and every 4 subsequent cycles. In clone 4, rosetting remained low in the absence of drug, but was restored to a high rosette frequency after only 5 cycles in the presence of drug, showing the reversibility of selection.

### Use of the single variant editing strategy to add epitope tags to PfEMP1 allowing detection with commercial antibodies

To illustrate a useful application of the single variant editing strategy, we introduced epitope tags (myc, 3xFLAG or HA) into the N-terminus of IT4VAR60 PfEMP1. The PfCas9-BSD-2A-IT4var60 plasmid was modified to incorporate the tag sequence upstream of the NTS-DBLα domain and transfected into IT4 parasites under blasticidin selection. Correct integration was confirmed in bulk cultures by integration PCR (Extended Data Fig. 1a-c), and clonal lines were subsequently derived by limiting dilution cloning and validated by integration PCR and Sanger sequencing, which confirmed accurate introduction of each tag at the intended locus (Extended Data Fig. 1d-f).

Expression of the epitope-tagged IT4VAR60 PfEMP1 on the IE surface was assessed by flow cytometry using custom polyclonal antibodies to IT4VAR60-DBLα and commercial monoclonal antibodies (mAbs) against the relevant epitope tag. In each parasite line, ∼85-90% of live IEs stained positively with both the PfEMP1 polyclonal antibodies and the anti-tag mAbs (Fig. 4a-c), showing that the tag did not prevent PfEMP1 trafficking to the IE surface, and that commercial mAbs can replace custom variant-specific antibodies for PfEMP1 detection in tagged parasite lines. As expected, the MFI for the polyclonal antibodies (which bind multiple epitopes) was higher than for the anti-tag mAbs, indicating brighter staining (Fig. 4a-c). The 3XFLAG mAb showed brighter staining than the anti-myc and anti-HA mAbs, also as expected due to the presence of three antibody epitopes per PfEMP1 molecule in the former compared to only one in the latter.

**Figure 4:**
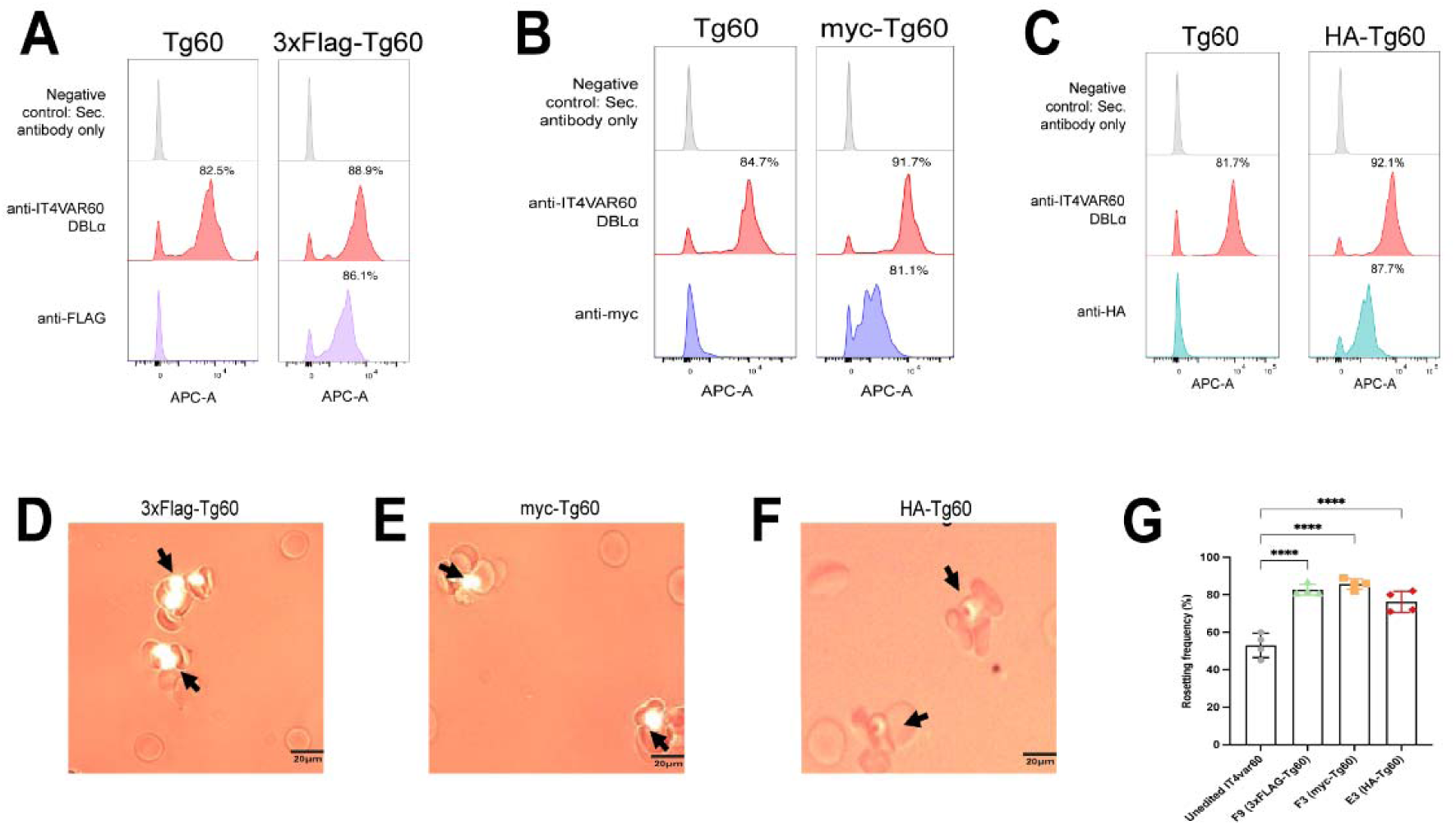
Infected erythrocyte (IE) surface expression and rosetting of epitope-tagged IT4VAR60. (A-C) Flow cytometry histograms showing IE surface expression of epitope-tagged IT4VAR60 in 3×FLAG (F9 clone), myc (F3 clone), and HA (E3 clone) lines. IEs were stained with anti-IT4VAR60DBLα antibodies and the corresponding anti-tag antibodies (anti-FLAG, anti-myc, or anti-HA). No surface staining was detected in untagged Tg60 parasites with anti-tag antibodies. Flow cytometry data are representative of at least two independent experiments. (D-F) Fluorescence/bright-field images of images of rosettes formed by 3xFlag-, myc- and HA-tagged IT4VAR60 clones. Rosettes are indicated by black arrowheads. Images were acquired at 400x magnification on a Leica DM 2000 fluorescent microscope. (G) Rosette frequency of 3xFlag-, myc-, and HA-tagged IT4VAR60 clones across four timepoints compared with unedited IT4var60. Mean and standard deviation are shown (****p < 0.0001, one-way ANOVA with Tukey’s multiple comparison test).

To determine if the N-terminal epitope tags affected parasite adhesion, the rosette frequency of the tagged parasite lines was assessed over one month in continuous culture under drug pressure. The epitope-tagged lines consistently showed ∼80% of IEs in rosettes, which was significantly higher than unedited parasites (Fig. 4d-g), and similar to that demonstrated by edited untagged IT4VAR60 parasites (Fig. 1c-d). These data demonstrate that epitope tags can be added to the N-terminus of PfEMP1 without disrupting IE surface expression or parasite adhesion, allowing detection of PfEMP1 with commercial mAbs.

### Use of the single variant editing strategy to generate PfEMP1 knock-out parasites

To demonstrate the utility of the editing strategy for reverse genetic experiments, we tested whether IT4VAR60 PfEMP1 is required for rosetting in it4var60-expressing P. falciparum IEs. Previous work suggested that rosetting may be partly RIFIN-mediated in this parasite line (Goel et al., 2015). The CRISPR/Cas9 var gene editing plasmid described above was used with a modified homology repair template to generate a functional knock-out (KO), in which a frame-shift mutation was introduced into it4var60 exon I, resulting in a premature stop codon after the third amino acid of the IT4VAR60 PfEMP1 (Extended Data Fig. 2a). After transfection into IT4 parasites and drug selection, Tg60-KO parasites became visible on Giemsa smear 12 days later and were confirmed by genomic DNA extraction and integration PCR from the bulk culture (Extended Data Fig. 2b). Subsequently, Tg60-KO clonal lines were obtained by limiting dilution, and two clones were validated by integration PCR and Sanger sequencing, confirming insertion of a single nucleotide (“T”) after the 9^th^ nucleotide of it4var60 exon I, resulting in a frameshift and premature stop codon (Extended Data Fig. 2c). The var gene transcripts in the Tg60-KO clones were profiled by qRT-PCR, showing it4var60 as the predominant transcript in the clones, with the bsd gene also transcribed, as expected (Extended Data Fig. 2d).

To determine the phenotypic effects of the Tg60-KO genome edit, flow cytometry was performed to evaluate surface and intracellular expression of IT4VAR60 PfEMP1. Live IE surface staining was performed using rabbit IgG antibodies specific to various IT4VAR60 domains (Ghumra et al., 2012). IT4VAR60 PfEMP1 was not detected on the surface of the Tg60-KO IEs, whereas all antibodies showed strong positive staining of unedited IT4var60-expressing IEs (Fig. 5a). Staining was also conducted with anti-IT4VAR60-DBLa IgG on fixed and permeabilised IEs to determine if any intracellular IT4VAR60 remained in the Tg60-KO parasites. Unedited IT4var60 IEs showed positive staining but no intracellular IT4VAR60 PfEMP1 was detected in the Tg60-KO clones (Fig. 5b), providing further validation of the successful functional knock-out of IT4VAR60. The Tg60-KO clones were grown under drug pressure for one month, and the rosette frequency was recorded weekly. Over this period, no rosettes were seen (Fig. 5c-d), hence rosetting was completely abolished in the Tg60-KO clones. RIFINs were previously implicated in rosetting in this parasite line, especially in blood group A erythrocytes (Goel et al., 2015), therefore Tg60-KO parasites were cultured for 2 weeks in blood group A erythrocytes and monitored for rosetting. No rosettes were observed during this period, confirming that PfEMP1 is required for rosetting in it4var60-transcribing IEs, even in blood group A erythrocytes (Fig. 5e).

**Figure 5:**
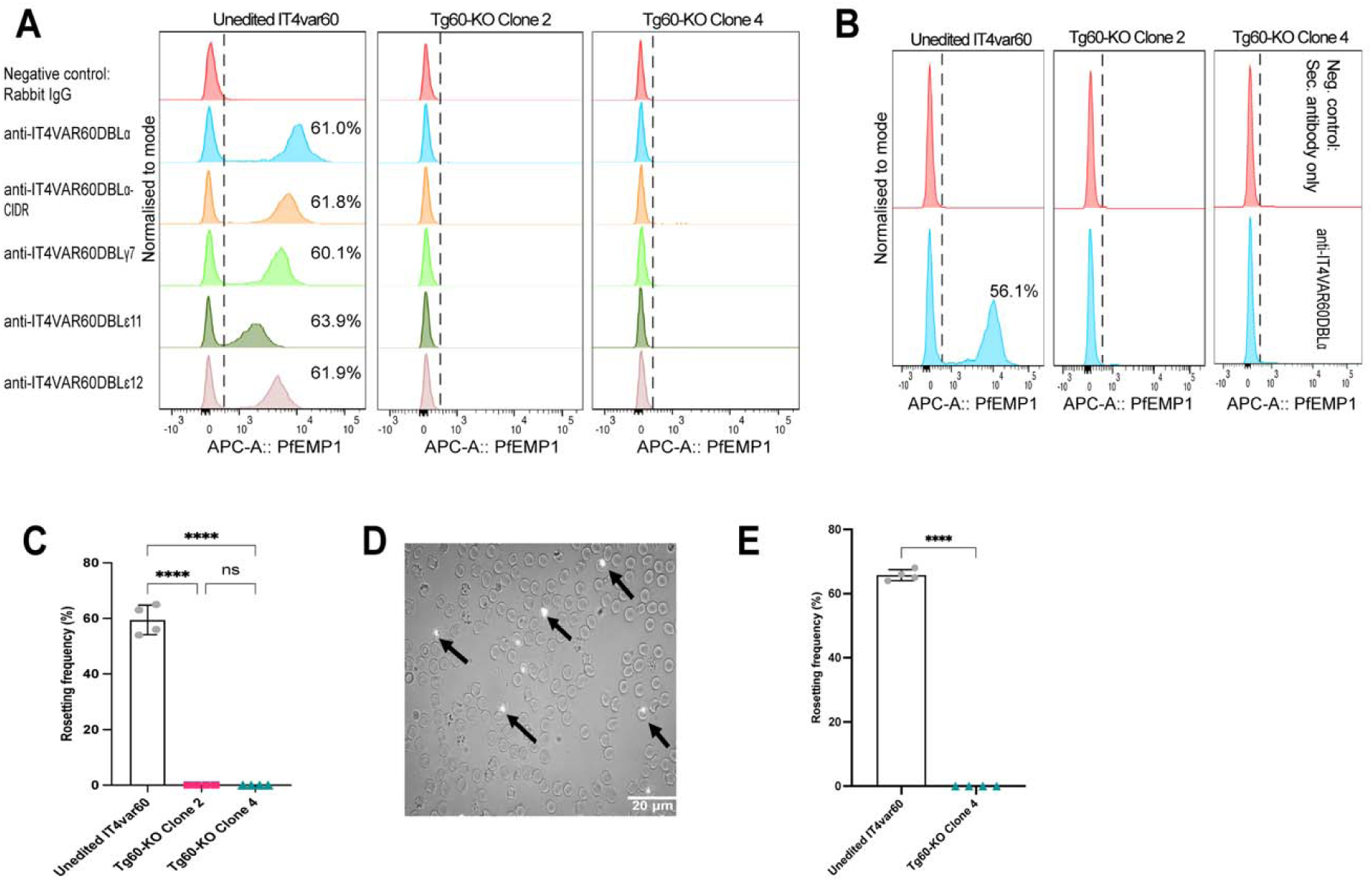
Functional knockout (KO) of IT4VAR60 abolishes rosetting. (A) Flow cytometry analysis of IT4VAR60 surface expression in two Tg60-KO clones (2 and 4) compared with unedited IT4 parasites selected for rosetting by traditional means. Live IEs were stained with polyclonal antibodies to various IT4VAR60 domains (DBLα, DBLα-CIDR, DBLγ7, DBLε11 and DBLε12). (B) Intracellular staining with anti-IT4VAR60 NTS-DBL_a_ IgG in Tg60-KO clones. Showing absence of detectable IT4VAR60 expression in the clones. (C) Rosette frequency analysis over one month, demonstrating complete loss of rosetting in Tg60-KO. Differences between Tg60-KO clones and unedited IT4var60 WT parasites were compared using One-way ANOVA with Tukey’s multiple comparison test (**** p < 0.0001). Error bars represent standard deviation (SD). (D) Fluorescence/bright-field image of Tg60-KO parasite clone stained with ethidium bromide showing absence of rosette formation. IEs are in indicated by black arrow. Images were taken with a Yenway CMOS 5mpx camera at x400 magnification on a Leica DM2000 fluorescent microscope. (E) Rosette frequency of Tg60-KO parasites cultured in blood group A erythrocytes. Time points represent 2-day intervals. Statistical comparison was performed using an unpaired t test (**** p < 0.0001, unpaired t test). Error bars represent SD.

### Use of the single variant editing strategy to generate targeted amino acid substitutions in PfEMP1

Finally, we used the single variant editing strategy to generate parasite lines with mutations in PfEMP1. Previous work by Angeletti et al (Angeletti et al., 2015) using E. coli-derived recombinant proteins, identified several key amino acids within the IT4VAR60 N-terminal DBLα domain that reduced erythrocyte binding when mutated (Supplementary Fig. 2a and Table 1). The effect of these mutations on rosetting in live parasites has not previously been tested. We adapted the single variant strategy to introduce the targeted amino acid substitutions into the it4var60 HDR template to investigate their effect on PfEMP1 function. Three mutant parasite lines were generated, Mut A (Y73A K263E), Mut G (K263E), and Mut F (K97A). Structural modelling based on the DBLα domain suggests that these residues are surface-exposed and unlikely to affect overall domain conformation, suggesting that any functional effects are likely due to disruption of receptor-binding interactions rather than structural perturbation (Supplementary Fig. 2a,b). For Mut A and Mut G, genome editing efficiency was enhanced by constructing a dual-guide version of the plasmid that included a second gRNA targeting a site proximal to residue K263. Successful introduction of the mutations was confirmed by integration PCR (Extended Data Fig. 3a-c). Independent clones were generated for each mutant by limiting dilution cloning and validated by integration PCR and Sanger sequencing (Extended Data Fig. 3d-e).

**Table 1:** Differential effects of IT4VAR60 DBLα mutations on live parasite and recombinant protein adhesion.

|  | Live parasites | Recombinant protein |
| --- | --- | --- |

| **Mutant** | Transgenic parasite clone | % of IE PfEMP1+ by flow cytometry | Expected RF (%)<sup>^</sup> | Observed RF (%)<sup>§</sup> | Rosette Inhibition<sup>&</sup> (%) | rMut* reduction in erythrocyte binding (%) |
| --- | --- | --- | --- | --- | --- | --- |
| **Mut A**<br>(Y73A, K263E) | F10 | 84.2 | 78.1 | 24.0 | **69.3** | **100** |
|  | B10 | 93.8 | 87.0 | 28.0 | **67.8** |  |
| **Mut G**<br>(K263E) | G6 | 77.6 | 72.0 | 21.8 | **69.7** | **~50** |
|  | D3 | 81.2 | 75.3 | 25.9 | **65.6** |  |
| **Mut F**<br>(K97A) | E4 | 92.7 | 86.0 | 68.1 | **20.8** | **~75** |
|  | G4 | 92.3 | 85.6 | 85.3 | **0** |  |

<sup>^</sup>The expected rosette frequency (RF) was derived by multiplying the % of PfEMP1+ IEs by a scaling factor (0.93) derived from the rosetting of edited Tg60 wild type parasites as described in the methods
<sup>§</sup>average of four timepoints.
<sup>&</sup>rosette inhibition is expected-observed RF/expected RF x100
\* Recombinant protein mutant (rMut) data are from Angeletti *et al* (2015) and are based on the MFI of each mutant protein in an erythrocyte binding assay compared to the IT4VAR60 DBLα wild type protein control.

To assess whether the introduced mutations affected PfEMP1 surface display on IEs, flow cytometry was performed using antibodies to IT4VAR60 domains (NTS-DBLα, DBLα-CIDR, DBLγ7, DBLε12). All mutant clones expressed IT4VAR60 PfEMP1 on the IE surface at levels comparable to Tg60, indicating that surface expression was unaffected by the mutations (Fig. 6a–c). The effect of the mutations on rosetting was assessed over the course of one month in culture under blasticidin selection (Fig. 6d-f). Rosette inhibition was calculated as described in the methods, representing the percentage reduction in adhesion relative to the expected level given the proportion of IEs with surface PfEMP1. Based on recombinant protein studies (Angeletti et al., 2015), Mut A (Y73A K263E) was predicted to abolish adhesion, Mut G (K263E) to reduce it by ∼50%, and Mut F (K97A) by ∼75%. In the transgenic P. falciparum lines, Mut A and Mut G both reduced adhesion by ∼65-70%, whereas Mut F had minimal effect (Table 1). Therefore, IT4VAR60 Mut G showed similar effects in both recombinant protein and live parasites, reducing adhesion by 50-70%. However, the other two mutations gave different results in live parasites compared to the published recombinant protein data (Angeletti et al., 2015). Mut A was expected to abolish adhesion, but the effects of Y73A and K263E together were no different to K263E alone (Mut G), suggesting that Y73 does not form an essential part of the binding site between PfEMP1 and uninfected erythrocytes. Mut F (K97A) had minimal impact on rosetting in live parasites, despite predictions from the recombinant protein assays of >75% inhibition. These results highlight the need for reverse genetic experiments in P. falciparum parasites to provide insights into PfEMP1 function. The single variant genome editing strategy described here provides the means to carry out such experiments in any PfEMP1 variant in a timely and efficient manner.

**Figure 6:**
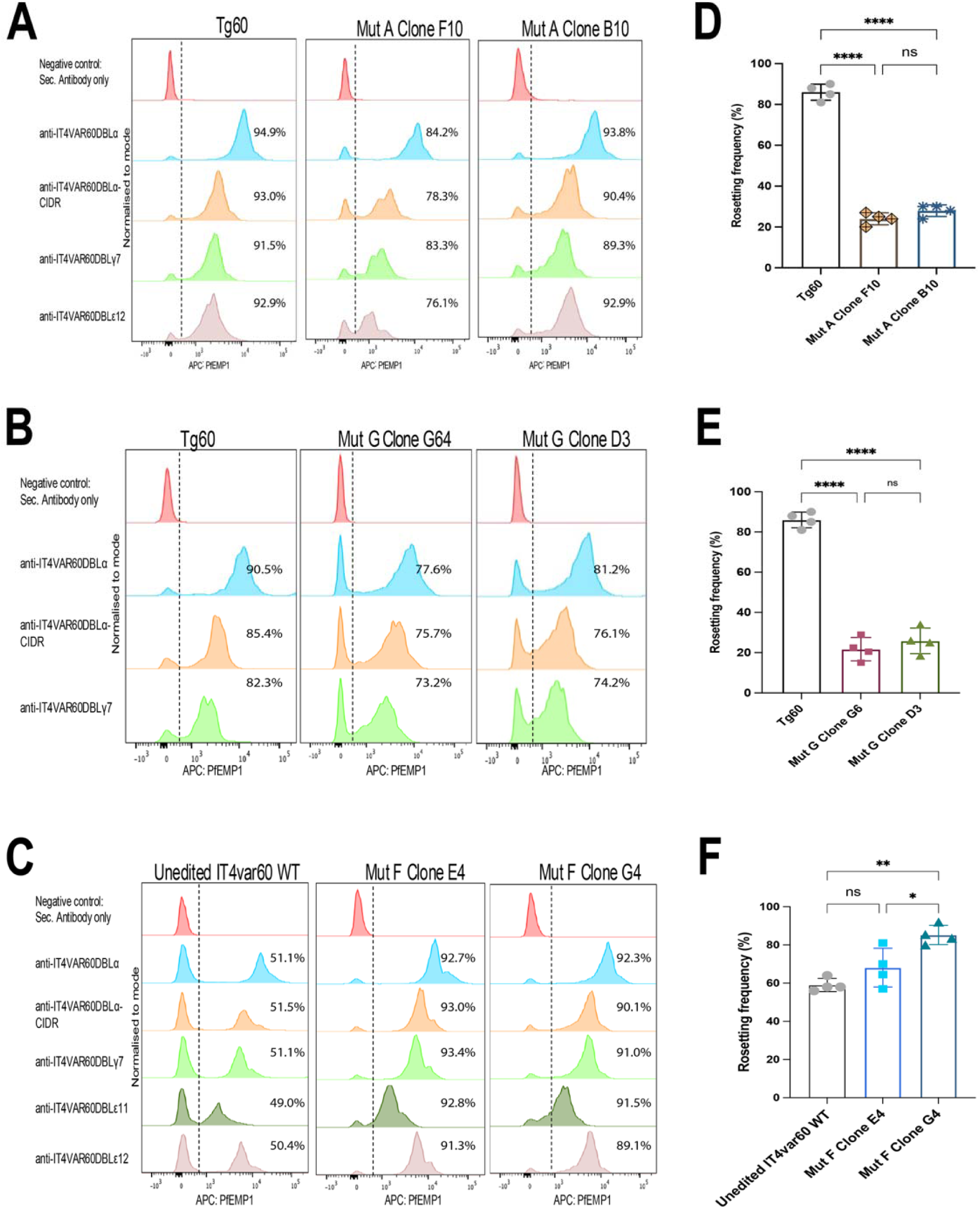
Infected erythrocyte (IE) surface expression of PfEMP1 and rosetting of IT4VAR60 DBLα mutants. (A) Flow cytometry analysis of Mut A clones F10 and B10 stained with domain-specific antibodies (NTS-DBLα, DBLα-CIDR, DBLγ7, DBLε12) compared to a Tg60 clone with the wild-type sequence. (B) Surface expression of Mut G clones G6 and D3 stained with NTS-DBLα, DBLα-CIDR, and DBLγ7 antibodies, compared to a Tg60 clone with the wild-type sequence. (C) Surface expression of Mut F clones E4 and G4 stained with NTS-DBLα, DBLα-CIDR, DBLγ7, DBL4ε11, and DBLε12 antibodies compared to unedited IT4var60 parasites. Flow cytometry data are representative of at least two independent experiments. (D-F) Mean and standard deviation of rosette frequency of Mut A, Mut G and Mut F clones compared to Tg60 or unedited IT4var60 parasites. Statistical comparisons were performed using one-way ANOVA with Tukey’s multiple comparison test; **** p < 0.0001, ns = not significant.

## Discussion

A major barrier to investigating PfEMP1 biology has been the inability to experimentally control mutually exclusive expression of var genes in live P. falciparum (Frank & Deitsch, 2006; Scherf et al., 1998; Voss et al., 2006). Previous studies showed that var transcription can be influenced by linking promoter activity to drug selection (Dzikowski et al., 2006; Dzikowski et al., 2007; Voss et al., 2006), but did not enable simultaneous genetic modification of defined loci. Here, we establish a CRISPR/Cas9-based strategy that combines P. falciparum genome editing (Ghorbal et al., 2014; Wagner et al., 2014; Zhang et al., 2014) with drug selection (Birnbaum et al., 2017), focused on a defined endogenous var gene locus. By linking a bsd cassette to the target locus via a 2A ribosomal skip peptide (de Felipe et al., 2006), parasite survival when exposed to drug is dependent on transcription of that locus, resulting in parasite populations expressing a single PfEMP1 variant. This provides a framework for studying PfEMP1 in its native form, overcoming a long-standing constraint in P. falciparum biology imposed by antigenic variation.

Validation of this approach in it4var60 resulted in efficient CRISPR/Cas9 mediated integration of the bsd-2A cassette, and dominant transcription of the target gene (Fig. 1), confirming that locus-specific control can be achieved in live parasites under drug pressure. The mature transgenic parasites expressed IE surface IT4VAR60 PfEMP1 and showed the IgM-positive rosetting phenotype, consistent with the established role of IT4VAR60 in rosette formation and non-immune IgM-Fc binding (Akhouri et al., 2016; Angeletti et al., 2012; Ghumra et al., 2012; McLean et al., 2025). The successful application of the single variant editing strategy to additional var genes and parasite genotypes, such as the virulence-associated variants IT4VAR19 and HB3VAR03, demonstrates the potential for broad applicability of the approach across diverse variants and parasite genomes. This is particularly significant given the extensive diversity in the var gene family. The main requirement for editing any culture-adapted P. falciparum genotype is the availability of the parasite’s whole genome sequence to allow design of specific gRNAs. Compared with conventional enrichment methods that require time-consuming repeated selection by panning or density gradients over extended periods (Albrecht et al., 2011; Ghumra et al., 2012; Handunnetti et al., 1992; Horrocks et al., 2002), the approach described here enables rapid generation of parasites expressing a single-variant that can be stably maintained under drug selection for months.

Despite the near-complete IE surface expression of PfEMP1 in the transgenic parasites, a small PfEMP1-negative subpopulation was consistently observed, even within clonal lines (Fig. 2). This may reflect dying parasites that have switched to alternative var transcripts but have not yet been eliminated from the culture, or parasites in a transient low-var or “var-null” state with little or no PfEMP1 expression, as described recently (Florini et al., 2025). Attempts to grow the PfEMP1-negative parasites in culture failed, suggesting that the former explanation is most likely. These results, and the reversibility of selection demonstrated by growing the transgenic parasites with and without drug (Fig. 3), emphasise that the editing strategy used here does not eliminate antigenic variation. Parasites continue to switch, but those that do are killed if drug is present. Hence, edited parasites could be a useful tool in future studies of factors affecting var gene switching by providing cultures with homogenous starting var gene transcription that can be released from drug pressure and followed through time.

Epitope tagging further demonstrated the utility and versatility of the system. N-terminal tagging of IT4VAR60 with HA, Myc, and 3xFLAG was confirmed using anti-tag antibodies, and showed correct trafficking to the IE surface, with retention of the rosetting phenotype (Fig. 4). Given the limited availability of variant-specific antibodies, this provides a broadly applicable approach for detection, localisation, and purification using commercially available reagents. This will substantially expand the experimental accessibility of var gene repertoires in diverse P. falciparum genotypes, while simultaneously reducing the need to use animals to generate custom variant-specific antibodies.

A key biological finding of this study is that editing a frameshift into the it4var60 gene completely abolished rosetting (Fig. 5). This establishes a direct requirement for the IT4VAR60 variant in rosetting, providing definitive genetic validation in live IEs. Our editing system also enables targeted interrogation of PfEMP1 function at the level of domains, sub-domains or individual amino acids. We show that targeted mutagenesis of IT4VAR60 DBLα residues (Y73, K97 and K263) previously implicated in erythrocyte binding from recombinant protein studies (Angeletti et al., 2015), produced effects on rosetting in transgenic parasites (Fig. 6), however clear differences were seen compared to the recombinant protein data (Table 1). These discrepancies demonstrate the need to examine PfEMP1-mediated adhesion within the structural and cellular context of the IE. Factors such as domain organisation, intermolecular interactions (Rajan Raghavan et al., 2023) and post-translational modifications (Dorin-Semblat et al., 2019) may influence adhesion phenotypes. Together, these findings emphasise the limitation of recombinant approaches and underscore the necessity of studying PfEMP1 in live parasites.

Some limitations of the work should be considered. Reverse genetic modifications at the 5’ end of the var gene coding sequence are easily incorporated, but modifications further from the transcription start site will require longer HDR regions and additional guide RNAs, making the design and cloning of editing constructs more challenging. Furthermore, any var gene with a very high “off” rate of switching may be difficult to target by this method, as switching may occur too rapidly to allow transgenic parasites to be selected. The method described here selects for transcription and modification of a var gene of interest, but does not guarantee IE surface expression of PfEMP1, which relies on the parasite having intact trafficking pathways, while effective cytoadhesion may also require other parasite genes to be expressed appropriately (Cronshagen et al., 2025). For Tg60 parasites, drug-selection alone maintained high levels of rosetting for months and no additional phenotypic enrichment was required. For other cytoadhesion phenotypes, occasional selection for associated phenotypes that facilitate cytoadhesion such as knob expression (Crabb et al., 1997) may be required.

Future applications of the var gene selection and editing strategy described here include interrogation of PfEMP1-host receptor binding interaction sites by domain-, subdomain- and motif-swaps, incorporation of fluorescent proteins for localisation studies, and inclusion of BIO-ID tags for proximity labelling experiments (Cronshagen et al., 2025; Kimmel et al., 2022) to identify novel host receptors bound by PfEMP1. Single-variant parasite lines will also provide a powerful tool for immunological studies, as PfEMP1 is expressed in its native conformation on the IE surface, allowing direct assessment of antibody binding and functional immune responses without reliance on recombinant proteins or multiplex bead-based assays (Rambhatla et al., 2022; Travassos et al., 2026; Walker et al., 2024). This will offer a more physiologically relevant framework for studying naturally acquired immunity and may facilitate quicker identification of protective immune responses and functionally relevant epitopes for vaccine development.

The combined selection and editing strategy described here could also be applied to other muti-gene families in Plasmodium, such as rifin and stevor, building upon previous SLI approaches for these genes (Omelianczyk et al., 2020), although the lack of mutually exclusive expression in these families (Witmer et al., 2012) would mean that single variant expression may not occur. Similarly, the strategy could open-up the functional investigation of specific variant surface antigens in other pathogenic Plasmodium species such as those encoded by vir genes in P. vivax and SICAvar in P. knowlesi, and related multi-gene families in rodent models (yir, pir and cir genes in respectively, P. yoeli, P.berghei and P. chabaudi). The bsd gene in the editing plasmid could easily be swapped for other drug resistance genes to allow generation of parasite lines in which different drugs select for different multi-gene family members. The approach could also be adapted for multi-gene families from other microbial pathogens with mutually exclusive expression systems such as vsg genes in Trypanosomes (Florini et al., 2022).

Overall, this work establishes a broadly applicable framework for investigating the role of PfEMP1 in malaria parasite biology, enabling section of specific variants combined with precise genetic manipulations. The method offers the potential to transform how PfEMP1 is studied and generate new insights into the molecular basis of antigenic variation, cytoadherence and malaria pathogenesis.

## Materials and Methods

### Ethical permission

Human serum and erythrocytes used in this study were obtained from the Scottish National Blood Transfusion Service (SNBTS), Edinburgh, UK, with ethical approval reference numbers 19∼08, 22∼11 and 25∼10. Experimental work was approved by the University of Edinburgh School of Biological Sciences ethical committee (reference arowe-0002). All genetic modification work was conducted under University of Edinburgh GM Safety Committee approval SBS_642.

### P. falciparum strains and cultures

The parasite lines used in this study were IT4, selected to express it4var60 (Ghumra et al., 2012) and it4var19 (Claessens et al., 2012) and HB3, selected to express hb3var03 (Claessens et al., 2012). Plasmodium falciparum parasites were cultured in fresh O+ erythrocytes at 2% haematocrit in complete RPMI-1640 (Lonza, BE12-167) supplemented with 25 mM HEPES, 2 mM L-glutamine, 16 mM glucose, 25 μg/mL gentamycin, 5% pooled human serum (SNBTS), and 0.25% AlbuMAX II (Gibco, 11021037), adjusted to pH 7.2–7.4. Cultures were maintained at 37 °C in 96% N₂, 5% CO₂, 1% O₂ and kept between 2–10% parasitaemia. The transgenic parasite lines reported here are available after publication from the European Malaria Reagent Repository (https://emrr.bio.ed.ac.uk).

### Donor template construction

All plasmids, primers and sgRNAs used in this study are listed in Extended Data Tables 1-3 and var gene sequences are from PlasmoDB (https://plasmodb.org/plasmo/app). The nucleotide sequence of the editing plasmids is given in the Supplementary information. A homologous donor cassette was made comprising ∼550 bp of promoter sequence upstream of exon I and ∼600 bp from the 51 region of exon I as left and right homology arms flanking a bsd coding sequence linked via a 2A peptide. For it4var60 (PfIT_120060300), the homology arms were amplified from IT4 genomic DNA, incorporating KasI/SpeI sites in the promoter fragment and SalI/SpeI sites in the exon I fragment. The bsd sequence was amplified from pDC2-cam-eGFP-BSD-attP (Straimer et al., 2012) without its stop codon and with NheI/HindIII sites for in-frame 2A fusion. Fragments were assembled into a linearised AmpR backbone using In-Fusion® HD cloning to generate IT4var60prom-BSD-ExonI. A synthetic 2A sequence was inserted via NheI/HindIII digestion and ligation to yield IT4var60prom-BSD-2A-ExonI, which was used as a versatile (modular) plasmid for subsequent homology substitution or exon I modification. Equivalent donor backbones were generated for it4var19 (PfIT_010005000) and hb3var03 (PfHB3_130080100).

### Construction of Cas9-expressing plasmids

For each target locus, the donor cassette was excised using KasI/SpeI and ligated into a pDC2-cam-Cas9-U6-hDHFR (Ng et al., 2016) backbone digested with the same enzymes. For it4var60, this generated PfCas9-IT4var60prom-BSD-2A-IT4var60-ExonI. The redundant hDHFR cassette was subsequently removed by HpaI/SmaI digestion, followed by blunt-end ligation, yielding the reduced-size derivative PfCas9-IT4var60prom-BSD-2A-IT4var60-ExonI(b), which served as the backbone for subsequent constructs. The same cloning strategy was applied to additional var genes (it4var19 and hb3var03). All constructs were verified by diagnostic restriction digestion and Sanger sequencing prior to parasite transfection.

### Generation of derivative transfection vectors

For it4var60 knockout, a 300 bp synthetic repair fragment (Twist Bioscience) containing a single thymine insertion after nucleotide 9 of exon I and synonymous shield mutations was synthesised and cloned via HindIII/MfeI into the versatile donor backbone and subcloned into the PfCas9-BSD(b) backbone using KasI/SpeI. Targeted amino acid substitutions within exon I (MutA, MutG, MutF) were introduced using synthetic repair fragments incorporating the desired nucleotide substitutions together with synonymous shield mutations. Modified fragments were cloned into the donor backbone via HindIII/MfeI and assembled into the Cas9 vector using KasI/SpeI. For N-terminal epitope tagging of it4var60, synthetic fragments encoding the 2A peptide followed by either HA (YPYDVPDYA), myc (EQKLISEEDL), or 3×FLAG (DYKDHDG–DYKDHDI–DYKDDDDK) and the first 125 bp of exon I were synthesised. The fragments were designed with flanking NheI/MfeI restriction sites and incorporated M13F/M13R priming sequences to enable amplification. PCR-amplified fragments were purified, digested with NheI/MfeI, and ligated into the corresponding sites within PfCas9-IT4var60prom-BSD-2A-IT4var60-ExonI(b).

### sgRNA design

sgRNAs targeting exon I of each var gene were designed using Benchling (20 nt guide length; 51-NGG PAM) against the IT4 or HB3 genome. Guides were selected based on predicted on-target and off-target scores (>50 preferred) and proximity to the intended modification site and verified by in silico analysis against strain-specific genomic sequences to confirm locus-specific uniqueness. Complementary guide oligonucleotides were synthesised (IDT) with 51-ATTG and 51-AAAC overhangs compatible with BbsI digestion. ∼2 μg of pDC2-cam-Cas9-U6-hDHFR vector was digested with BbsI-HF, dephosphorylated, and gel-purified. Oligos were phosphorylated and annealed (37 °C for 30 min; 94 °C for 5 min; gradual cooling to 25 °C), diluted 1:200, and ligated to ∼50 ng of digested vector backbone using T4 DNA ligase. Sanger sequencing-validated sgRNA cassette was subsequently incorporated into the appropriate Cas9 donor plasmids for transfection. For mutations separated by >100 bp, dual-guide constructs were generated by cloning a second sgRNA into pDC2-cam-Cas9-U6-hDHFR and subcloning the amplified U6–guide–scaffold cassette into the single-guide Cas9 donor plasmid via SbfI/XbaI. Dual-guide constructs were verified by Sanger sequencing.

### Parasite transfection and drug selection

Transfections were performed by direct electroporation of ring-stage parasites (Fidock & Wellems, 1997). ∼50 µg plasmid DNA was combined with Cytomix buffer (120 mM KCl, 0.15 mM CaCl_2_, 2mM EGTA, 5 mM MgCl_2_, 10 mM K_2_HPO_4_/KH_2_PO_4_, 25 mM HEPES, pH 7.6) to a final volume of 400 µL. A 10 mL parasite culture (2% haematocrit; ∼5-10% parasitaemia) was washed in 37 °C pre-warmed Cytomix buffer, and the pellet resuspended in the DNA-Cytomix mixture. The suspension was transferred to a pre-chilled 0.2 cm electroporation cuvette and electroporated using a Gene Pulser II (0.31 kV, 950 µF). Cells were immediately transferred to a 10 mL pre-warmed complete medium with ∼50 µL PCV fresh erythrocytes, gassed and incubated at 37°C. Blasticidin S (Sigma-Aldrich, 203350-25MG) selection (2.51µg/ml) (Pandey et al., 2016) commenced 48 h post-transfection. Cultures were maintained at 2% haematocrit with daily medium changes for 7 days, then every other day until parasites reappeared. Fresh erythrocytes (∼50 µL PCV) were added at least once a week. Transfectants were typically detectable by Giemsa-stained thin blood smear after 10-14 days.

### Limiting dilution cloning

Clonal parasite lines were generated from bulk cultures by limiting dilution as previously described (Kirkman et al., 1996). Briefly, parasitaemia was determined microscopically, and cultures were serially diluted in complete RPMI at 2% haematocrit with blasticidin S to achieve ∼ 3, 1, or 0.3 infected erythrocytes per well. Aliquots (100 µL) of each dilution were distributed into 96-well plates, placed in a humidified chamber, gassed (96% N_2_, 5% CO_2_ and 1% O_2_), and incubated at 37 °C. After one week, medium was replaced with fresh complete medium (containing 0.4% haematocrit fresh erythrocytes and blasticidin). Following a further week of incubation, wells were screened by Giemsa-stained thick blood smears. Wells yielding ∼50% positivity at the lowest parasite dilution were expanded in six-well plates (2 mL) and subsequently in 10 mL culture flasks.

### Extraction of genomic DNA

Genomic DNA was extracted from parasite cultures using the QIAamp Micro DNA Kit (Qiagen, 56304) according to the manufacturer’s protocol. DNA concentration was estimated using a NanoDrop 2000 Spectrophotometer.

### PCR to confirm genomic integration

Integration at the intended locus was confirmed by PCR using parasite gDNA as template and locus-specific primers spanning the 5’ UTR (F1), bsd cassette (R1) 2A sequence (F2) and var exon I (R2) (Extended Data Table 1). Primer combinations F1R1 and F2R2 confirmed 51 and 31 integration events respectively, while F1R2 spans the full modified and wildtype loci, confirming full-length donor integration or the intact wildtype depending on fragment size differences. PCR reactions were performed with an initial denaturation at 94 °C for 2 min, followed by 35 cycles of 94 °C for 5 s, 60 °C for 15 s, and 65 °C for 1 min 30 s, with a final extension at 65 °C for 10 min. Products were resolved on 1.2% agarose gels, gel-purified and subjected to Sanger sequencing to confirm accurate integration and, where applicable, the presence of intended sequence modifications.

### Assessment of rosette frequency

Rosette frequency was assessed by incubating ∼200 μL of parasite culture suspension (∼2% haematocrit) with ethidium bromide (20 μg/mL) and examining wet preparations by simultaneous fluorescence and bright field microscopy (Leica DM2000, 40x objective). A rosette was defined as an infected erythrocyte binding ≥2 uninfected erythrocytes. Two fields (100 infected erythrocytes per field; 200 total) were counted, and rosette frequency was expressed as the proportion of rosettes among the counted infected erythrocytes. Sample identity was masked during scoring to minimise observer bias.

### RNA extraction and cDNA synthesis

Parasite cultures (two independent transgenic clones and wild-type parasites) were synchronised by 5% D-sorbitol treatment and harvested at the ring stage (3-18 h post-invasion), when var transcription predominates (Kyes et al., 2007). Synchronous cultures (>5% parasitaemia) were pelleted and lysed in 10 volumes of pre-warmed TRIzol (37 °C) for 5 min at 37 °C, then stored at −70 °C until processing. For RNA extraction, thawed TRIzol lysates were mixed with bromochlorporane (0.2 x TRIzol volume), vortexed for 15 s, incubated for 3 min at room temperature, and centrifuged (9,000 xg, 15 min, 4 °C) to achieve phase separation. The aqueous phase (∼500 μL per 1 mL TRIzol) was transferred to a fresh RNase-free tube, mixed 1:1 with 100% ethanol, and applied to RNA Clean & Concentrator-5 columns (Zymo Research, R1013). On-column DNase I treatment (15 min at room temperature) was performed during purification. RNA was eluted in 20 μL DNase/RNase-free water pre-warmed to 65 °C and quantified using a NanoDrop 2000 Spectrophotometer. To ensure complete removal of genomic DNA, ∼2.5 μg RNA was subjected to a second DNase treatment using the TURBO DNase kit (Invitrogen, AM1907) (37 °C, 20 min), followed by DNase inactivation and centrifugation.

First-strand cDNA synthesis was performed using Superscript III First Strand Synthesis Kit (Invitrogen, 180800-51). DNase-treated RNA (0.5–1 μg) was combined with random hexamers and dNTPs, heated at 65 °C for 5 min, and chilled on ice. Reverse transcription reactions were performed according to the manufacturer’s protocol and incubated at 25 °C for 10 min, 50 °C for 50 min, and 85 °C for 5 min. cDNA samples were treated with RNAse H (2 U, 37 °C for 20 min). Successful cDNA synthesis and absence of genomic DNA contamination were confirmed by PCR.

### Profiling of var gene expression

Transcript levels of var genes were analysed by qRT-PCR using IT4 gene-specific primer sets described previously (Viebig et al., 2007). Reactions were performed on a LightCycler 480 Instrument II (Roche, 04729749001) using SYBR Green master mix (Roche, 04707516001) in 384-well plates. Each reaction contained premixed primer pairs, cDNA template, and SYBR Green master mix. Cycling conditions were: pre-incubation at 95°C, 15 min, 40 amplification cycles of 95 °C for 30 s, 50 °C for 40 s, and 65 °C, 50 s), melt curve analysis (95 °C for 5s, 65 °C for 1 min, ramp to 97 °C with 5 acquisitions per °C) and cooling at 40°C for 10 s. Relative transcript levels were calculated using the 2^-ΔCT^ method (Livak & Schmittgen, 2001), normalised to the housekeeping gene adenylosuccinate lyase (adsl). The mean cycle threshold (C_T_) value of adsl was subtracted from the mean C_T_ value of each sample to obtain ΔC_T_ value for each sample. Relative expression levels were plotted using GraphPad Prism.

### PfEMP1 antibodies

Rabbit polyclonal antibodies (purified total IgG) to IT4VAR60 NTS-DBLα (Ghumra et al., 2012), IT4VAR19 NTS-DBLα (Azasi et al., 2018) and HB3VAR03 NTS-DBLα (Claessens et al., 2012) were made as described previously. Similarly, polyclonal antibodies to other domains of IT4VAR60 (DBLα-CIDR, DBLγ7 and DBLε12) were made by expressing the recombinant proteins in E. coli and immunising rabbits as described (Ghumra et al., 2012).

### Flow cytometry

Surface and intracellular PfEMP1 expression were analysed by flow cytometry as previously described (McLean et al., 2024) using either 96-well plates (Corning, 3799) or FACS tubes (Corning, 352058). Parasites cultures at 2% haematocrit (predominantly mature trophozoite/schizont stages) were washed in PBS/1% Ig-free BSA. For surface staining, cells were incubated with 20 μg/mL rabbit IgG against the relevant PfEMP1 domain (45 min at 37 °C), washed twice in PBS and incubated (30 min, protected from light) with Alexa Fluor 647-conjugated anti-rabbit IgG (Invitrogen, A-21244), Vybrant DyeCycle Violet (Thermo Fisher Scientific, V35003, 1/2500) and ethidium bromide (Sigma-Aldrich, 20 μg/mL final concentration). For intracellular staining, cells were fixed in 0.5% paraformaldehyde for 10 min at room temperature, permeabilised with PBS/0.01% Triton X-100 for 10 min, blocked in PBS/0.1% BSA for 30 min, and stained as above. For IgM staining, an anti-human IgM heavy chain antibody (Biorad, MCA1662) was used at 20 μg/ml final concentration. For epitope tagged parasite lines, 20 μg/ml final concentration of commercial monoclonal antibodies were used for staining (anti-HA.11, Biolegend 901501; anti-FLAG M2, Sigma F1804; anti-myc, SinoBiological 100029-MM07). For plate-based assays, wells were pre-blocked with FACS buffer (PBS/0.1% BSA), 20 μL cell suspension added per well (duplicates), with centrifugation (1120 x g, 4 min) between washes. For tube-based assays, 1 mL culture was processed similarly. After staining, cells were washed twice in PBS and once in PBS/0.5% paraformaldehyde, resuspended in FACS buffer, and analysed on a BD LSRFortessa (BD Biosciences) using 405 nm, 561 nm, and 640 nm lasers. For each sample, 100,000 events were acquired from a 100 μL volume at a flow rate of 2 μL/s. Data were analysed with a FlowJo software (v.10.8.2) with the gating shown in Figure 2a.

### Calculation of Rosette Inhibition

to quantify the effect of IT4VAR60 DBLa mutations on rosette formation, the percentage reduction in observed RF relative to the expected RF was determined. The expected RF was defined as the proportion of PfEMP1+ mature IEs. Rosette inhibition values were calculated using the formula:

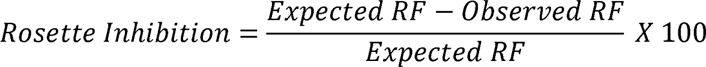

To account for cycle-to-cycle variability and inherent variation in rosetting assay, a scaling factor (0.93), derived from the ratio of mean observed RF (86%) to the mean proportion of mature of IEs expressing surface IT4VAR60 (92.7%) across four independent experiments was applied to standardise the expected RF for each mutant prior to the rosette inhibition calculation.

Structural analysis: Secondary structure PfEMP1 DBL domain was predicted using the Phyre2 tool (http://www.sbg.bio.ic.ac.uk/phyre2). Analysis of the generated structure (.pdb file) was done with PyMOL (The PyMOL Molecular Graphics System, Version 2.0 Schrödinger, LLC).

Data visualisation and statistical analysis: Data visualisation was performed using GraphPad Prism (version 10; GraphPad Software La Jolla, CA). Statistical analyses were conducted to assess differences in RF between IT4VAR60 mutants and wild-type control. Comparisons between two groups were made using unpaired two-tailed t-tests, and multiple group comparisons were analysed using one-way ANOVA with Tukey’s multiple comparisons.

## Acknowledgements

This work was funded by the Darwin Trust of Edinburgh (PhD studentship to SEO and ND), the Royal Society (Newton International Fellowship to HMA) and the Wellcome Trust (Hosts, Pathogens and Global Health PhD studentship to JJ and BRO, and Senior Research Fellowship to JAR). We are grateful to Prof. Marcus Lee (University of Dundee) for kindly providing the pDC2-cam-Cas9-U6-hDHFR plasmid and protocols. We are also grateful to Prof. JJ Lopez-Rubio and Prof. Catherine Merrick for advice on P. falciparum transfection.

## Supph

**Extended Data Figure 1:**
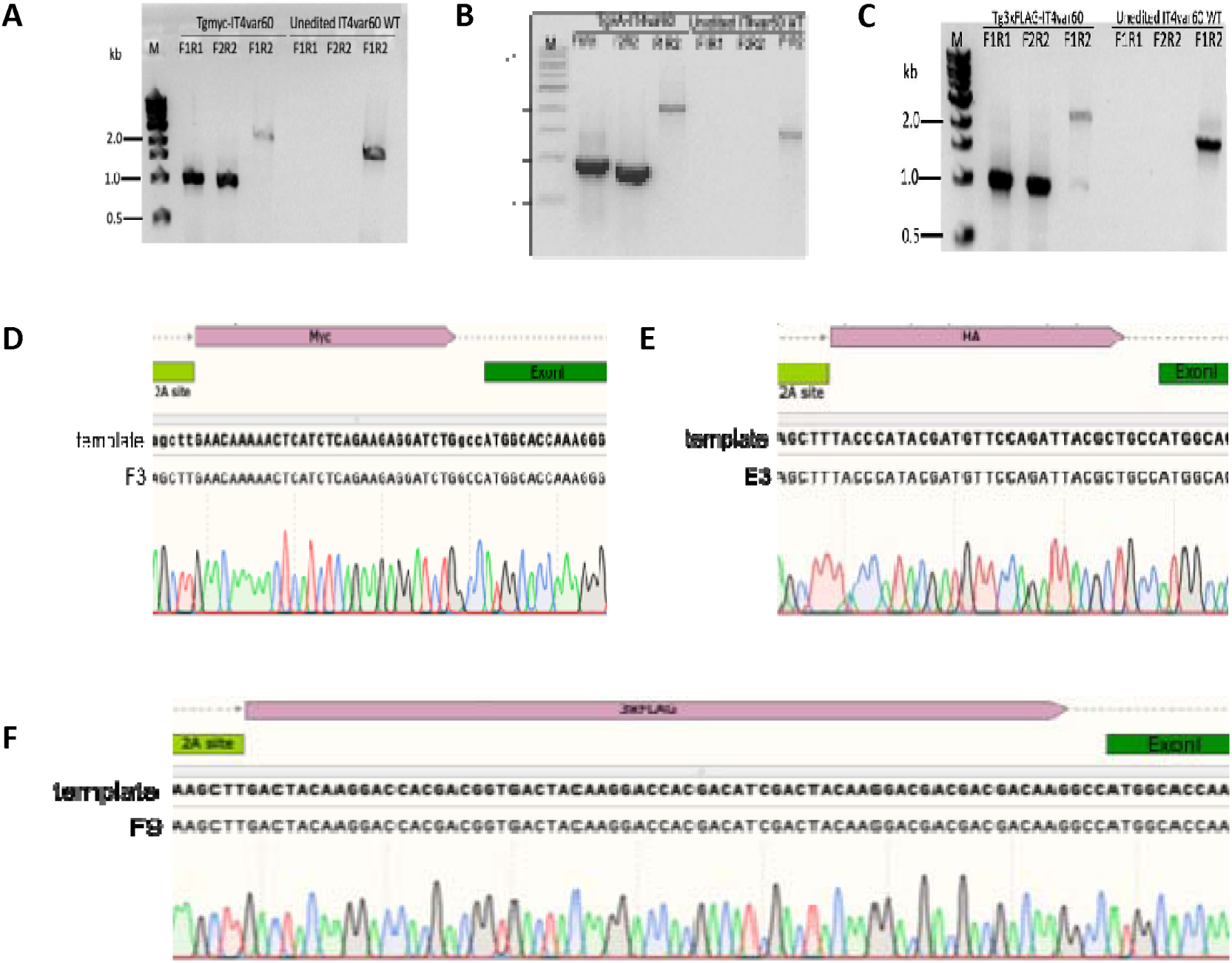
Genetic validation of epitope-tagged IT4VAR60 parasites. (A-C) Integration PCR performed on genomic DNA from bulk cultures of myc-, HA-, and 3×FLAG-tagged IT4VAR60 parasites using primers F1/R1, F2/R2 and F1/R2. (D-F) Representative Sanger sequencing chromatograms of myc-tagged (F3 clone), HA-tagged (E3 clone) and 3×FLAG-tagged (F9 clone) IT4var60 parasites. The 2A peptide is shown in bright green, epitope tag in pink, and IT4var60 exon I in dark green. Alignment of plasmid template and corresponding parasite genomic sequence confirms precise in-frame integration.

**Extended Data Figure 2:**
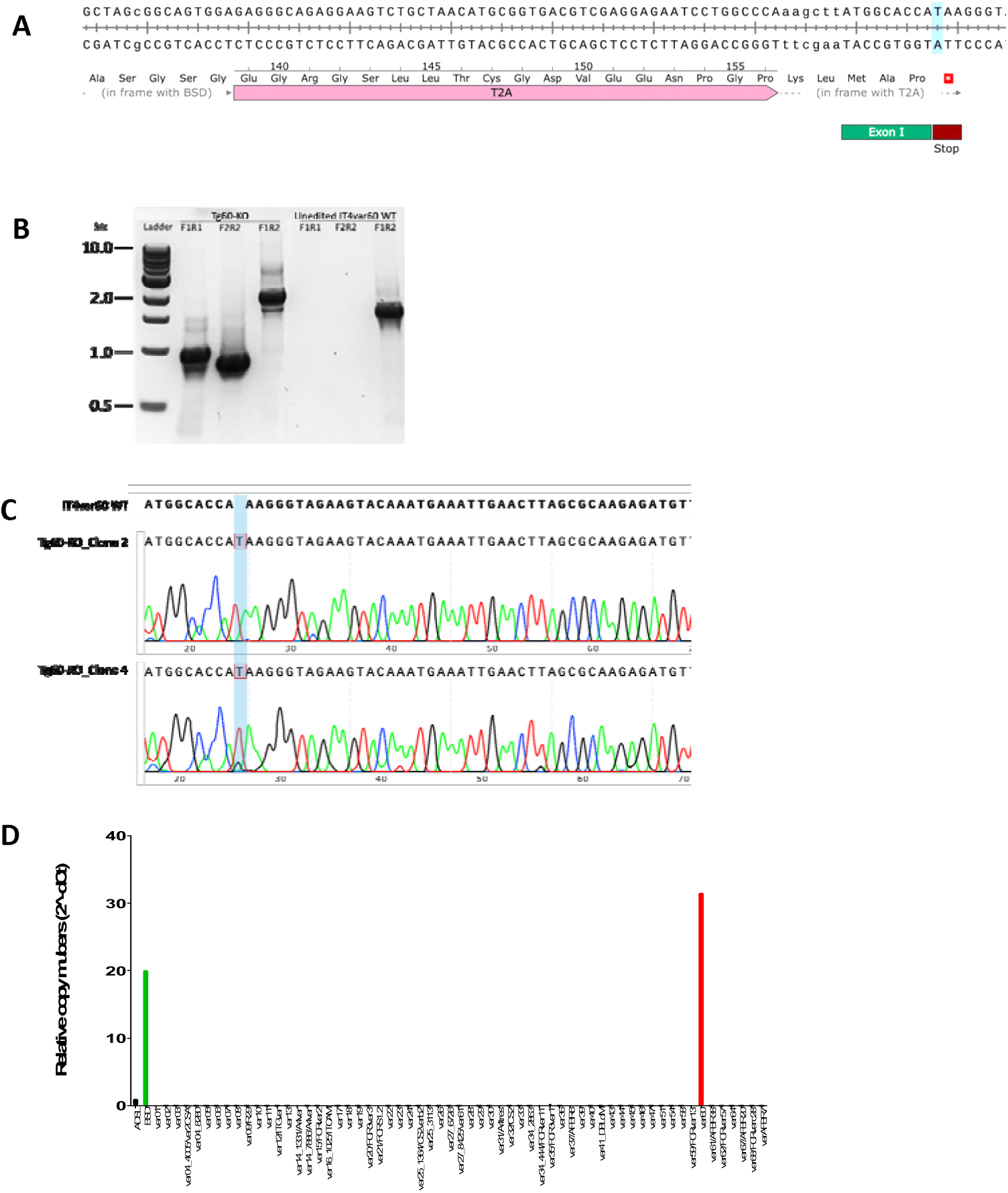
Genetic validation and qRT-PCR of IT4VAR60-KO parasites. (A) Schematic of the IT4var60-KO homology repair template. A single thymine (“T”, highlighted in blue) nucleotide was inserted after the 9^th^ nucleotide of the coding sequence, resulting in a frameshift and premature stop codon after the third codon of IT4var60 exon I. (B) Integration PCR of the IT4var60-KO bulk culture confirming successful integration at the targeted locus. (C) Sanger sequencing chromatograms of IT4var60-KO clones 2 and 4 showing insertion of the single thymine (“T”, highlighted). (D) Representative qRT-PCR graph of IT4var60-KO (clone 4) showing it4var60 (red) as the predominant transcript and detection of bsd, normalised to adsl.

**Extended Data Figure 3:**
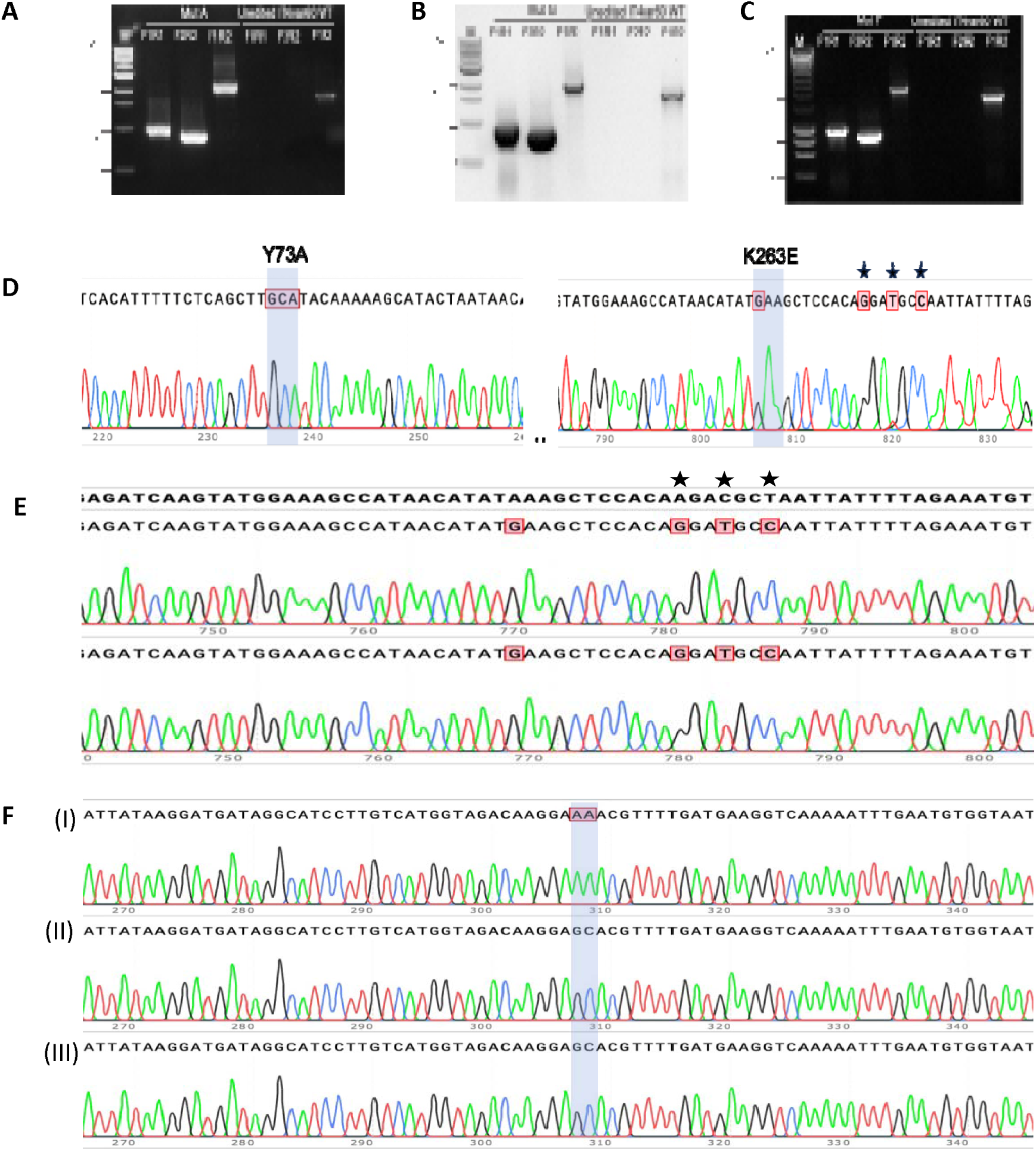
Genetic validation of IT4VAR60 DBLα mutants. (A-C) Integration PCR confirmation of CRISPR/Cas9 edited IT4VAR60 DBLα mutants - Mut A (Y73A K263E), Mut G (K263E) and Mut F (K97A). Integration was verified using it4var60 integration PCR primer pairs F1/R1, F2/R2 and F1/R2, with amplicons of expected sizes observed in all three mutant parasite lines. (D-F) Sanger sequencing validation of IT4VAR60 DBLα mutants. (D) Mut A clone F10 showing substitutions of Y73A (TAT ➔ GCA) and K263E (AAA ➔ GAA); (E) Mut G clone G6 (top) and D3 (bottom) showing substitution of K263 (AAA ➔ GAA); (F) Mut F: (I) unedited IT4VAR60 control and (II – III) Mut F clones E4 and G4 showing substitution of K97A (AAA ➔ GCA). Synonymous shield mutations introduced to prevent Cas9 re-cutting are indicated by Ii.. Shield mutations were not required for Mut F, as insertion of the bsd-2A cassette disrupted the guide target sequence at the promoter-exon I boundary.

**Extended data Table 1:** Plasmids.

| Name | Role | Source |
| --- | --- | --- |
| pDC2-cam-Cas9-U6-hDHFR | All-in-one CRISPR/Cas9 backbone | Gift from Prof Marcus Lee |
| Versatile IT4var60prom-BSD-ExonI | Intermediate donor template plasmid (*it4var60* promoter-*bsd*-2A-*it4var60*_exon I) | This study |
| Cas9-IT4var60prom-BSD-2A-IT4var60_ExonI | CRISPR/Cas9 transfection plasmid with *bsd* and *it4var60* donor template. Used to generate Tg60 parasites. | This study |
| Cas9-IT4var60prom-BSD-2A-IT4var60_ExonI(b) | Optimised smaller CRISPR/Cas9 plasmid (legacy DHFR removed) | This study |
| Versatile IT4var19prom-BSD-ExonI | Intermediate donor template plasmid for *it4var19* | This study |
| Versatile HB3var03prom-BSD-ExonI | Intermediate donor template plasmid for *hb3var03* | This study |
| Cas9-IT4var19prom-BSD-2A-IT4var19_ExonI | Used to generate TgIT4VAR19 parasites | This study |
| Cas9-IT4var19prom-BSD-2A-IT4var19_ExonI | Used to generate TgHB3VAR03 parasites | This study |
| Cas9-IT4var60prom-BSD-2A-IT4var60_ExonI_KO | Used to generate IT4VAR60 knock-out parasites | This study |
| Cas9-IT4var60prom-BSD-2A-IT4var60_ExonI-MutA | Used to generate Mut A | This study |
| Cas9-IT4var60prom-BSD-2A-IT4var60_ExonI-MutF | Used to generate Mut F | This study |
| Cas9-IT4var60prom-BSD-2A-IT4var60_ExonI-MutG | Used to generate Mut G | This study |

**Extended data Table 2:**
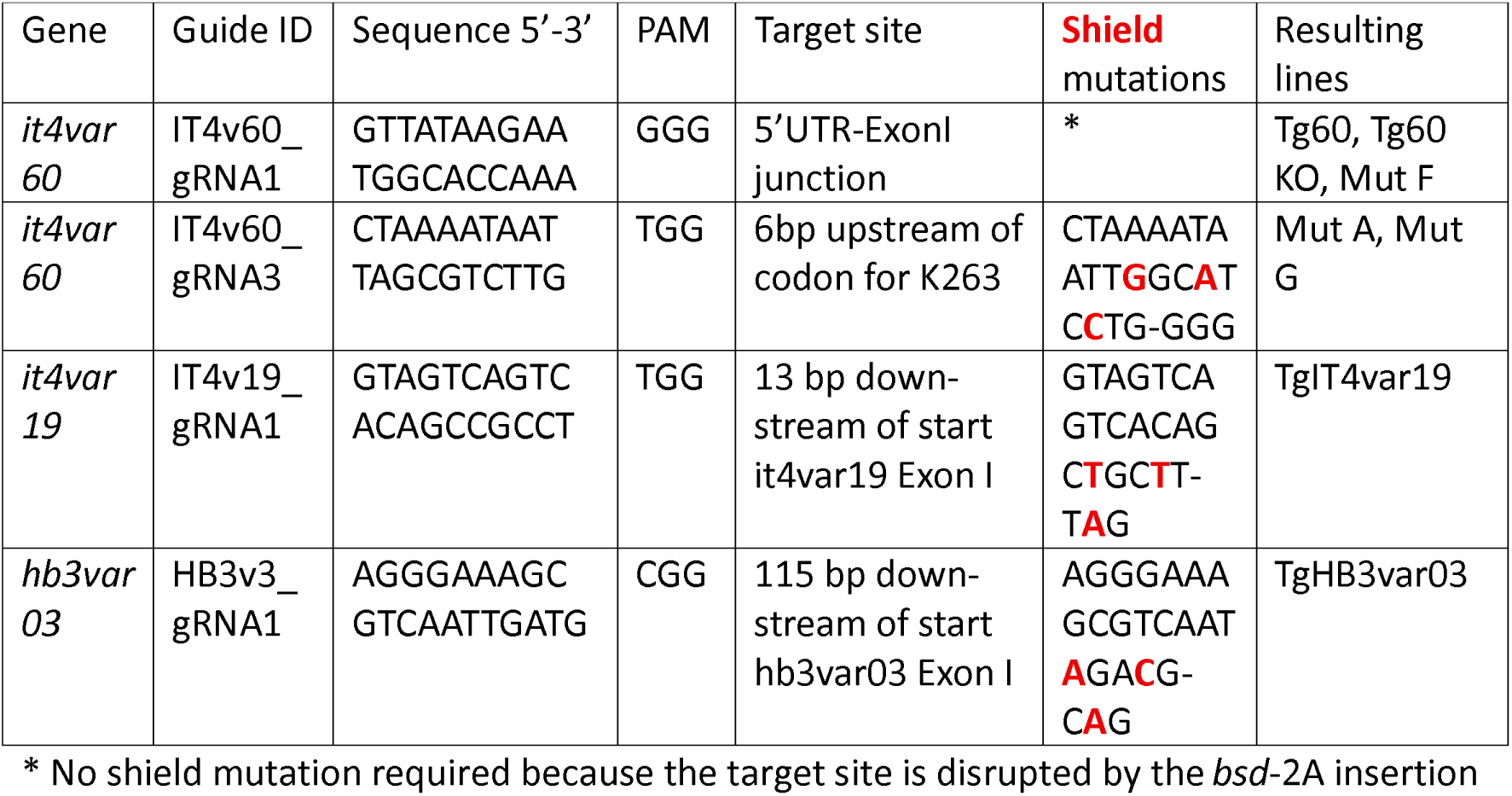
guideRNAs.

**Extended data Table 3:**
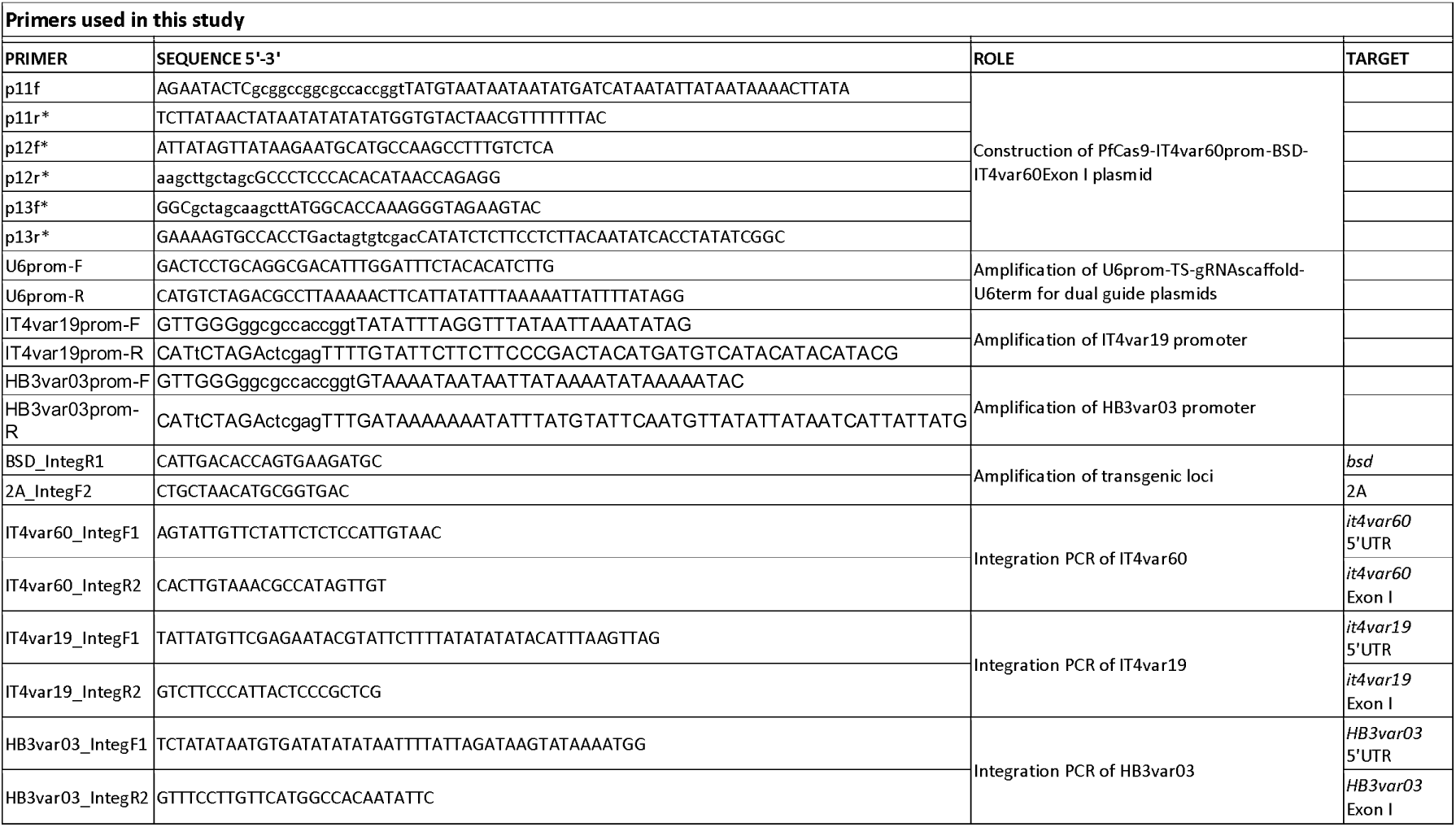
primers.

## References

Akhouri, R. R., Goel, S., Furusho, H., Skoglund, U., & Wahlgren, M. (2016). Architecture of Human IgM in Complex with P. falciparum Erythrocyte Membrane Protein 1. Cell Rep, 14(4), 723–736. 10.1016/j.celrep.2015.12.067

Albrecht, L., Moll, K., Blomqvist, K., Normark, J., Chen, Q., & Wahlgren, M. (2011). var gene transcription and PfEMP1 expression in the rosetting and cytoadhesive Plasmodium falciparum clone FCR3S1.2. Malar J, 10, 17. 10.1186/1475-2875-10-17

Angeletti, D., Albrecht, L., Blomqvist, K., Quintana Mdel, P., Akhter, T., Bächle, S. M., Sawyer, A., Sandalova, T., Achour, A., Wahlgren, M., & Moll, K. (2012). Plasmodium falciparum rosetting epitopes converge in the SD3-loop of PfEMP1-DBL1α. PLoS One, 7(12), e50758. 10.1371/journal.pone.0050758

Angeletti, D., Sandalova, T., Wahlgren, M., & Achour, A. (2015). Binding of subdomains 1/2 of PfEMP1-DBL1α to heparan sulfate or heparin mediates Plasmodium falciparum rosetting. PLoS One, 10(3), e0118898. 10.1371/journal.pone.0118898

Avril, M., Bernabeu, M., Benjamin, M., Brazier, A. J., & Smith, J. D. (2016). Interaction between Endothelial Protein C Receptor and Intercellular Adhesion Molecule 1 to Mediate Binding of Plasmodium falciparum-Infected Erythrocytes to Endothelial Cells. mBio, 7(4). 10.1128/mBio.00615-16

Avril, M., Tripathi, A. K., Brazier, A. J., Andisi, C., Janes, J. H., Soma, V. L., Sullivan, D. J., Jr., Bull, P. C., Stins, M. F., & Smith, J. D. (2012). A restricted subset of var genes mediates adherence of Plasmodium falciparum-infected erythrocytes to brain endothelial cells. Proc Natl Acad Sci U S A, 109(26), E1782–1790. 10.1073/pnas.1120534109

Azasi, Y., Lindergard, G., Ghumra, A., Mu, J., Miller, L. H., & Rowe, J. A. (2018). Infected erythrocytes expressing DC13 PfEMP1 differ from recombinant proteins in EPCR-binding function. Proc Natl Acad Sci U S A, 115(5), 1063–1068. 10.1073/pnas.1712879115

Barry, A. E., Leliwa-Sytek, A., Tavul, L., Imrie, H., Migot-Nabias, F., Brown, S. M., McVean, G. A., & Day, K. P. (2007). Population genomics of the immune evasion (var) genes of Plasmodium falciparum. PLoS Pathog, 3(3), e34. 10.1371/journal.ppat.0030034

Baruch, D. I., Pasloske, B. L., Singh, H. B., Bi, X., Ma, X. C., Feldman, M., Taraschi, T. F., & Howard, R. J. (1995). Cloning the P. falciparum gene encoding PfEMP1, a malarial variant antigen and adherence receptor on the surface of parasitized human erythrocytes. Cell, 82(1), 77–87. 10.1016/0092-8674(95)90054-3

Biggs, B. A., Goozé, L., Wycherley, K., Wollish, W., Southwell, B., Leech, J. H., & Brown, G. V. (1991). Antigenic variation in Plasmodium falciparum. Proc Natl Acad Sci U S A, 88(20), 9171–9174. 10.1073/pnas.88.20.9171

Birnbaum, J., Flemming, S., Reichard, N., Soares, A. B., Mesén-Ramírez, P., Jonscher, E., Bergmann, B., & Spielmann, T. (2017). A genetic system to study Plasmodium falciparum protein function. Nat Methods, 14(4), 450–456. 10.1038/nmeth.4223

Bryant, J. M., Regnault, C., Scheidig-Benatar, C., Baumgarten, S., Guizetti, J., & Scherf, A. (2017). CRISPR/Cas9 Genome Editing Reveals That the Intron Is Not Essential for var2csa Gene Activation or Silencing in Plasmodium falciparum. mBio, 8(4). 10.1128/mBio.00729-17

Claessens, A., Adams, Y., Ghumra, A., Lindergard, G., Buchan, C. C., Andisi, C., Bull, P. C., Mok, S., Gupta, A. P., Wang, C. W., Turner, L., Arman, M., Raza, A., Bozdech, Z., & Rowe, J. A. (2012). A subset of group A-like var genes encodes the malaria parasite ligands for binding to human brain endothelial cells. Proc Natl Acad Sci U S A, 109(26), E1772–1781. 10.1073/pnas.1120461109

Crabb, B. S., Cooke, B. M., Reeder, J. C., Waller, R. F., Caruana, S. R., Davern, K. M., Wickham, M. E., Brown, G. V., Coppel, R. L., & Cowman, A. F. (1997). Targeted gene disruption shows that knobs enable malaria-infected red cells to cytoadhere under physiological shear stress. Cell, 89(2), 287–296. 10.1016/s0092-8674(00)80207-x

Cronshagen, J., Allweier, J., Mesén-Ramírez, J. P., Stäcker, J., Vaaben, A. V., Ramón-Zamorano, G., Naranjo-Prado, I., Graser, M., López-Barona, P., Ofori, S., Jansen, P., Hornebeck, J., Kieferle, F., Murk, A., Martin, E., Castro-Peña, C., Bártfai, R., Lavstsen, T., Bruchhaus, I., & Spielmann, T. (2025). A system for functional studies of the major virulence factor of malaria parasites. Elife, 13. 10.7554/eLife.103542

Crosnier, C., Wanaguru, M., McDade, B., Osier, F. H., Marsh, K., Rayner, J. C., & Wright, G. J. (2013). A library of functional recombinant cell-surface and secreted P. falciparum merozoite proteins. Mol Cell Proteomics, 12(12), 3976–3986. 10.1074/mcp.O113.028357

Dalgaard, N., Olsen, R. W., Adams, Y., Mendez, B. L., Rudbaek, J. J., Bisholm, S. A. G., Mousiliou, A., Walker, M. R., Hviid, L., Tahar, R., Ndam, N. T., Barfod, L., & Jensen, A. R. (2025). A monoclonal antibody selectively recognizing PfEMP1 proteins associated with cerebral malaria. Sci Rep, 15(1), 34732. 10.1038/s41598-025-18465-1

de Felipe, P., Luke, G. A., Hughes, L. E., Gani, D., Halpin, C., & Ryan, M. D. (2006). E unum pluribus: multiple proteins from a self-processing polyprotein. Trends Biotechnol, 24(2), 68–75. 10.1016/j.tibtech.2005.12.006

Dorin-Semblat, D., Tétard, M., Claës, A., Semblat, J. P., Dechavanne, S., Fourati, Z., Hamelin, R., Armand, F., Matesic, G., Nunes-Silva, S., Srivastava, A., Gangnard, S., Lopez-Rubio, J. J., Moniatte, M., Doerig, C., Scherf, A., & Gamain, B. (2019). Phosphorylation of the VAR2CSA extracellular region is associated with enhanced adhesive properties to the placental receptor CSA. PLoS Biol, 17(6), e3000308. 10.1371/journal.pbio.3000308

Doumbo, O. K., Thera, M. A., Koné, A. K., Raza, A., Tempest, L. J., Lyke, K. E., Plowe, C. V., & Rowe, J. A. (2009). High levels of Plasmodium falciparum rosetting in all clinical forms of severe malaria in African children. Am J Trop Med Hyg, 81(6), 987–993. 10.4269/ajtmh.2009.09-0406

Duffy, M. F., Maier, A. G., Byrne, T. J., Marty, A. J., Elliott, S. R., O’Neill, M. T., Payne, P. D., Rogerson, S. J., Cowman, A. F., Crabb, B. S., & Brown, G. V. (2006). VAR2CSA is the principal ligand for chondroitin sulfate A in two allogeneic isolates of Plasmodium falciparum. Mol Biochem Parasitol, 148(2), 117–124. 10.1016/j.molbiopara.2006.03.006

Dzikowski, R., Frank, M., & Deitsch, K. (2006). Mutually exclusive expression of virulence genes by malaria parasites is regulated independently of antigen production. PLoS Pathog, 2(3), e22. 10.1371/journal.ppat.0020022

Dzikowski, R., Li, F., Amulic, B., Eisberg, A., Frank, M., Patel, S., Wellems, T. E., & Deitsch, K. W. (2007). Mechanisms underlying mutually exclusive expression of virulence genes by malaria parasites. EMBO Rep, 8(10), 959–965. 10.1038/sj.embor.7401063

Fidock, D. A., & Wellems, T. E. (1997). Transformation with human dihydrofolate reductase renders malaria parasites insensitive to WR99210 but does not affect the intrinsic activity of proguanil. Proc Natl Acad Sci U S A, 94(20), 10931–10936. 10.1073/pnas.94.20.10931

Florini, F., Visone, J. E., & Deitsch, K. W. (2022). Shared Mechanisms for Mutually Exclusive Expression and Antigenic Variation by Protozoan Parasites. Front Cell Dev Biol, 10, 852239. 10.3389/fcell.2022.852239

Florini, F., Visone, J. E., Hadjimichael, E., Malpotra, S., Nötzel, C., Kafsack, B. F. C., & Deitsch, K. W. (2025). scRNA-seq reveals transcriptional plasticity of var gene expression in Plasmodium falciparum for host immune avoidance. Nat Microbiol, 10(6), 1417–1430. 10.1038/s41564-025-02008-5

Frank, M., & Deitsch, K. (2006). Activation, silencing and mutually exclusive expression within the var gene family of Plasmodium falciparum. Int J Parasitol, 36(9), 975–985. 10.1016/j.ijpara.2006.05.007

Ghorbal, M., Gorman, M., Macpherson, C. R., Martins, R. M., Scherf, A., & Lopez-Rubio, J. J. (2014). Genome editing in the human malaria parasite Plasmodium falciparum using the CRISPR-Cas9 system. Nat Biotechnol, 32(8), 819–821. 10.1038/nbt.2925

Ghumra, A., Semblat, J. P., Ataide, R., Kifude, C., Adams, Y., Claessens, A., Anong, D. N., Bull, P. C., Fennell, C., Arman, M., Amambua-Ngwa, A., Walther, M., Conway, D. J., Kassambara, L., Doumbo, O. K., Raza, A., & Rowe, J. A. (2012). Induction of strain-transcending antibodies against Group A PfEMP1 surface antigens from virulent malaria parasites. PLoS Pathog, 8(4), e1002665. 10.1371/journal.ppat.1002665

Goel, S., Palmkvist, M., Moll, K., Joannin, N., Lara, P., Akhouri, R. R., Moradi, N., Öjemalm, K., Westman, M., Angeletti, D., Kjellin, H., Lehtiö, J., Blixt, O., Ideström, L., Gahmberg, C. G., Storry, J. R., Hult, A. K., Olsson, M. L., von Heijne, G.,…Wahlgren, M. (2015). RIFINs are adhesins implicated in severe Plasmodium falciparum malaria. Nat Med, 21(4), 314–317. 10.1038/nm.3812

Guillotte, M., Nato, F., Juillerat, A., Hessel, A., Marchand, F., Lewit-Bentley, A., Bentley, G. A., Vigan-Womas, I., & Mercereau-Puijalon, O. (2016). Functional analysis of monoclonal antibodies against the Plasmodium falciparum PfEMP1-VarO adhesin. Malar J, 15, 28. 10.1186/s12936-015-1016-5

Handunnetti, S. M., Gilladoga, A. D., van Schravendijk, M. R., Nakamura, K., Aikawa, M., & Howard, R. J. (1992). Purification and in vitro selection of rosette-positive (R+) and rosette-negative (R-) phenotypes of knob-positive Plasmodium falciparum parasites. Am J Trop Med Hyg, 46(4), 371–381. 10.4269/ajtmh.1992.46.371

Hasenkamp, S., Russell, K. T., & Horrocks, P. (2012). Comparison of the absolute and relative efficiencies of electroporation-based transfection protocols for Plasmodium falciparum. Malar J, 11, 210. 10.1186/1475-2875-11-210

Higgins, M. K. (2008). The structure of a chondroitin sulfate-binding domain important in placental malaria. J Biol Chem, 283(32), 21842–21846. 10.1074/jbc.C800086200

Horrocks, P., Pinches, R., Kyes, S., Kriek, N., Lee, S., Christodoulou, Z., & Newbold, C. I. (2002). Effect of var gene disruption on switching in Plasmodium falciparum. Mol Microbiol, 45(4), 1131–1141. 10.1046/j.1365-2958.2002.03085.x

Hsieh, F. L., Turner, L., Bolla, J. R., Robinson, C. V., Lavstsen, T., & Higgins, M. K. (2016). The structural basis for CD36 binding by the malaria parasite. Nat Commun, 7, 12837. 10.1038/ncomms12837

Joergensen, L. M., Salanti, A., Dobrilovic, T., Barfod, L., Hassenkam, T., Theander, T. G., Hviid, L., & Arnot, D. E. (2010). The kinetics of antibody binding to Plasmodium falciparum VAR2CSA PfEMP1 antigen and modelling of PfEMP1 antigen packing on the membrane knobs. Malar J, 9, 100. 10.1186/1475-2875-9-100

Juillerat, A., Lewit-Bentley, A., Guillotte, M., Gangnard, S., Hessel, A., Baron, B., Vigan-Womas, I., England, P., Mercereau-Puijalon, O., & Bentley, G. A. (2011). Structure of a Plasmodium falciparum PfEMP1 rosetting domain reveals a role for the N-terminal segment in heparin-mediated rosette inhibition. Proc Natl Acad Sci U S A, 108(13), 5243–5248. 10.1073/pnas.1018692108

Kaul, D. K., Roth, E. F., Jr., Nagel, R. L., Howard, R. J., & Handunnetti, S. M. (1991). Rosetting of Plasmodium falciparum-infected red blood cells with uninfected red blood cells enhances microvascular obstruction under flow conditions. Blood, 78(3), 812–819.

Kimmel, J., Kehrer, J., Frischknecht, F., & Spielmann, T. (2022). Proximity-dependent biotinylation approaches to study apicomplexan biology. Mol Microbiol, 117(3), 553–568. 10.1111/mmi.14815

Kirkman, L. A., Su, X. Z., & Wellems, T. E. (1996). Plasmodium falciparum: isolation of large numbers of parasite clones from infected blood samples. Exp Parasitol, 83(1), 147–149. 10.1006/expr.1996.0058

Kyes, S., Horrocks, P., & Newbold, C. (2001). Antigenic variation at the infected red cell surface in malaria. Annu Rev Microbiol, 55, 673–707. 10.1146/annurev.micro.55.1.673

Lau, C. K., Turner, L., Jespersen, J. S., Lowe, E. D., Petersen, B., Wang, C. W., Petersen, J. E., Lusingu, J., Theander, T. G., Lavstsen, T., & Higgins, M. K. (2015). Structural conservation despite huge sequence diversity allows EPCR binding by the PfEMP1 family implicated in severe childhood malaria. Cell Host Microbe, 17(1), 118–129. 10.1016/j.chom.2014.11.007

Leech, J. H., Barnwell, J. W., Aikawa, M., Miller, L. H., & Howard, R. J. (1984). Plasmodium falciparum malaria: association of knobs on the surface of infected erythrocytes with a histidine-rich protein and the erythrocyte skeleton. J Cell Biol, 98(4), 1256–1264. 10.1083/jcb.98.4.1256

Lennartz, F., Adams, Y., Bengtsson, A., Olsen, R. W., Turner, L., Ndam, N. T., Ecklu-Mensah, G., Moussiliou, A., Ofori, M. F., Gamain, B., Lusingu, J. P., Petersen, J. E., Wang, C. W., Nunes-Silva, S., Jespersen, J. S., Lau, C. K., Theander, T. G., Lavstsen, T., Hviid, L.,…Jensen, A. T. (2017). Structure-Guided Identification of a Family of Dual Receptor-Binding PfEMP1 that Is Associated with Cerebral Malaria. Cell Host Microbe, 21(3), 403–414. 10.1016/j.chom.2017.02.009

Lennartz, F., Smith, C., Craig, A. G., & Higgins, M. K. (2019). Structural insights into diverse modes of ICAM-1 binding by Plasmodium falciparum-infected erythrocytes. Proc Natl Acad Sci U S A, 116(40), 20124–20134. 10.1073/pnas.1911900116

Livak, K. J., & Schmittgen, T. D. (2001). Analysis of relative gene expression data using real-time quantitative PCR and the 2(-Delta Delta C(T)) Method. Methods, 25(4), 402–408. 10.1006/meth.2001.1262

McLean, F. E., Omondi, B. R., Diallo, N., Otoboh, S., Kifude, C., Abdi, A. I., Lim, R., Otto, T. D., Ghumra, A., & Rowe, J. A. (2025). Identification of novel PfEMP1 variants containing domain cassettes 11, 15 and 8 that mediate the Plasmodium falciparum virulence-associated rosetting phenotype. PLoS Pathog, 21(1), e1012434. 10.1371/journal.ppat.1012434

Miller, L. H., Baruch, D. I., Marsh, K., & Doumbo, O. K. (2002). The pathogenic basis of malaria. Nature, 415(6872), 673–679. 10.1038/415673a

Moll, K., Pettersson, F., Vogt, A. M., Jonsson, C., Rasti, N., Ahuja, S., Spångberg, M., Mercereau-Puijalon, O., Arnot, D. E., Wahlgren, M., & Chen, Q. (2007). Generation of cross-protective antibodies against Plasmodium falciparum sequestration by immunization with an erythrocyte membrane protein 1-duffy binding-like 1 alpha domain. Infect Immun, 75(1), 211–219. 10.1128/iai.00749-06

Ng, C. L., Siciliano, G., Lee, M. C., de Almeida, M. J., Corey, V. C., Bopp, S. E., Bertuccini, L., Wittlin, S., Kasdin, R. G., Le Bihan, A., Clozel, M., Winzeler, E. A., Alano, P., & Fidock, D. A. (2016). CRISPR-Cas9-modified pfmdr1 protects Plasmodium falciparum asexual blood stages and gametocytes against a class of piperazine-containing compounds but potentiates artemisinin-based combination therapy partner drugs. Mol Microbiol, 101(3), 381–393. 10.1111/mmi.13397

O’Donnell, R. A., Freitas-Junior, L. H., Preiser, P. R., Williamson, D. H., Duraisingh, M., McElwain, T. F., Scherf, A., Cowman, A. F., & Crabb, B. S. (2002). A genetic screen for improved plasmid segregation reveals a role for Rep20 in the interaction of Plasmodium falciparum chromosomes. Embo j, 21(5), 1231–1239. 10.1093/emboj/21.5.1231

Omelianczyk, R. I., Loh, H. P., Chew, M., Hoo, R., Baumgarten, S., Renia, L., Chen, J., & Preiser, P. R. (2020). Rapid activation of distinct members of multigene families in Plasmodium spp. Commun Biol, 3(1), 351. 10.1038/s42003-020-1081-3

Pandey, K., Ferreira, P. E., Ishikawa, T., Nagai, T., Kaneko, O., & Yahata, K. (2016). Ca(2+) monitoring in Plasmodium falciparum using the yellow cameleon-Nano biosensor. Sci Rep, 6, 23454. 10.1038/srep23454

Peters, J. M., Fowler, E. V., Krause, D. R., Cheng, Q., & Gatton, M. L. (2007). Differential changes in Plasmodium falciparum var transcription during adaptation to culture. J Infect Dis, 195(5), 748–755. 10.1086/511436

Rajan Raghavan, S. S., Turner, L., Jensen, R. W., Johansen, N. T., Jensen, D. S., Gourdon, P., Zhang, J., Wang, Y., Theander, T. G., Wang, K., & Lavstsen, T. (2023). Endothelial protein C receptor binding induces conformational changes to severe malaria-associated group A PfEMP1. Structure, 31(10), 1174–1183.e1174. 10.1016/j.str.2023.07.011

Rambhatla, J. S., Tonkin-Hill, G. Q., Takashima, E., Tsuboi, T., Noviyanti, R., Trianty, L., Sebayang, B. F., Lampah, D. A., Marfurt, J., Price, R. N., Anstey, N. M., Papenfuss, A. T., Damelang, T., Chung, A. W., Duffy, M. F., & Rogerson, S. J. (2022). Identifying Targets of Protective Antibodies against Severe Malaria in Papua, Indonesia, Using Locally Expressed Domains of Plasmodium falciparum Erythrocyte Membrane Protein 1. Infect Immun, 90(2), e0043521. 10.1128/iai.00435-21

Rask, T. S., Hansen, D. A., Theander, T. G., Gorm Pedersen, A., & Lavstsen, T. (2010). Plasmodium falciparum erythrocyte membrane protein 1 diversity in seven genomes--divide and conquer. PLoS Comput Biol, 6(9). 10.1371/journal.pcbi.1000933

Roberts, D. J., Craig, A. G., Berendt, A. R., Pinches, R., Nash, G., Marsh, K., & Newbold, C. I. (1992). Rapid switching to multiple antigenic and adhesive phenotypes in malaria. Nature, 357(6380), 689–692. 10.1038/357689a0

Rowe, J. A., Claessens, A., Corrigan, R. A., & Arman, M. (2009). Adhesion of Plasmodium falciparum-infected erythrocytes to human cells: molecular mechanisms and therapeutic implications. Expert Rev Mol Med, 11, e16. 10.1017/s1462399409001082

Rowe, J. A., Moulds, J. M., Newbold, C. I., & Miller, L. H. (1997). P. falciparum rosetting mediated by a parasite-variant erythrocyte membrane protein and complement-receptor 1. Nature, 388(6639), 292–295. 10.1038/40888

Scherf, A., Hernandez-Rivas, R., Buffet, P., Bottius, E., Benatar, C., Pouvelle, B., Gysin, J., & Lanzer, M. (1998). Antigenic variation in malaria: in situ switching, relaxed and mutually exclusive transcription of var genes during intra-erythrocytic development in Plasmodium falciparum. Embo j, 17(18), 5418–5426. 10.1093/emboj/17.18.5418

Smith, J. D., Chitnis, C. E., Craig, A. G., Roberts, D. J., Hudson-Taylor, D. E., Peterson, D. S., Pinches, R., Newbold, C. I., & Miller, L. H. (1995). Switches in expression of Plasmodium falciparum var genes correlate with changes in antigenic and cytoadherent phenotypes of infected erythrocytes. Cell, 82(1), 101–110. 10.1016/0092-8674(95)90056-x

Smith, J. D., Subramanian, G., Gamain, B., Baruch, D. I., & Miller, L. H. (2000). Classification of adhesive domains in the Plasmodium falciparum erythrocyte membrane protein 1 family. Mol Biochem Parasitol, 110(2), 293–310. 10.1016/s0166-6851(00)00279-6

Su, X. Z., Heatwole, V. M., Wertheimer, S. P., Guinet, F., Herrfeldt, J. A., Peterson, D. S., Ravetch, J. A., & Wellems, T. E. (1995). The large diverse gene family var encodes proteins involved in cytoadherence and antigenic variation of Plasmodium falciparum-infected erythrocytes. Cell, 82(1), 89–100. 10.1016/0092-8674(95)90055-1

Travassos, M. A., Han, P., Coulibaly, D., Stucke, E. M., Zhou, A. E., Shrestha, B., Nakajima, R., Jain, A., Taghavian, O., Jasinskas, A., Lawton, J. G., Laurens, M. B., Niangaly, A., Pike, A., Ouattara, A., Berry, A. A., Adams, M., Takala-Harrison, S., Kouriba, B.,…Thera, M. A. (2026). Distinct Plasmodium falciparum Erythrocyte Membrane Protein 1 Domain Cassette-Associated Antibody Gaps Specific to Cerebral Malaria and Severe Malarial Anemia. J Infect Dis. 10.1093/infdis/jiag256

Turner, L., Lavstsen, T., Berger, S. S., Wang, C. W., Petersen, J. E., Avril, M., Brazier, A. J., Freeth, J., Jespersen, J. S., Nielsen, M. A., Magistrado, P., Lusingu, J., Smith, J. D., Higgins, M. K., & Theander, T. G. (2013). Severe malaria is associated with parasite binding to endothelial protein C receptor. Nature, 498(7455), 502–505. 10.1038/nature12216

Udomsangpetch, R., Wåhlin, B., Carlson, J., Berzins, K., Torii, M., Aikawa, M., Perlmann, P., & Wahlgren, M. (1989). Plasmodium falciparum-infected erythrocytes form spontaneous erythrocyte rosettes. J Exp Med, 169(5), 1835–1840. 10.1084/jem.169.5.1835

Viebig, N. K., Levin, E., Dechavanne, S., Rogerson, S. J., Gysin, J., Smith, J. D., Scherf, A., & Gamain, B. (2007). Disruption of var2csa gene impairs placental malaria associated adhesion phenotype. PLoS One, 2(9), e910. 10.1371/journal.pone.0000910

Vigan-Womas, I., Guillotte, M., Juillerat, A., Hessel, A., Raynal, B., England, P., Cohen, J. H., Bertrand, O., Peyrard, T., Bentley, G. A., Lewit-Bentley, A., & Mercereau-Puijalon, O. (2012). Structural basis for the ABO blood-group dependence of Plasmodium falciparum rosetting. PLoS Pathog, 8(7), e1002781. 10.1371/journal.ppat.1002781

Voss, T. S., Healer, J., Marty, A. J., Duffy, M. F., Thompson, J. K., Beeson, J. G., Reeder, J. C., Crabb, B. S., & Cowman, A. F. (2006). A var gene promoter controls allelic exclusion of virulence genes in Plasmodium falciparum malaria. Nature, 439(7079), 1004–1008. 10.1038/nature04407

Wagner, J. C., Platt, R. J., Goldfless, S. J., Zhang, F., & Niles, J. C. (2014). Efficient CRISPR-Cas9-mediated genome editing in Plasmodium falciparum. Nat Methods, 11(9), 915–918. 10.1038/nmeth.3063

Walker, I. S., Dini, S., Aitken, E. H., Damelang, T., Hasang, W., Alemu, A., Jensen, A. T. R., Rambhatla, J. S., Opi, D. H., Duffy, M. F., Takashima, E., Harawa, V., Tsuboi, T., Simpson, J. A., Mandala, W., Taylor, T. E., Seydel, K. B., Chung, A. W., & Rogerson, S. J. (2024). A systems serology approach to identifying key antibody correlates of protection from cerebral malaria in Malawian children. BMC Med, 22(1), 388. 10.1186/s12916-024-03604-8

Witmer, K., Schmid, C. D., Brancucci, N. M., Luah, Y. H., Preiser, P. R., Bozdech, Z., & Voss, T. S. (2012). Analysis of subtelomeric virulence gene families in Plasmodium falciparum by comparative transcriptional profiling. Mol Microbiol, 84(2), 243–259. 10.1111/j.1365-2958.2012.08019.x

Zhang, C., Xiao, B., Jiang, Y., Zhao, Y., Li, Z., Gao, H., Ling, Y., Wei, J., Li, S., Lu, M., Su, X. Z., Cui, H., & Yuan, J. (2014). Efficient editing of malaria parasite genome using the CRISPR/Cas9 system. mBio, 5(4), e01414–01414. 10.1128/mBio.01414-14

