## Supplementary Figures 1 and 2 for "CRISPR/Cas9 genome editing to generate single variant *Plasmodium falciparum* lines and enable reverse genetic studies of PfEMP1 function in live parasites"

**D**

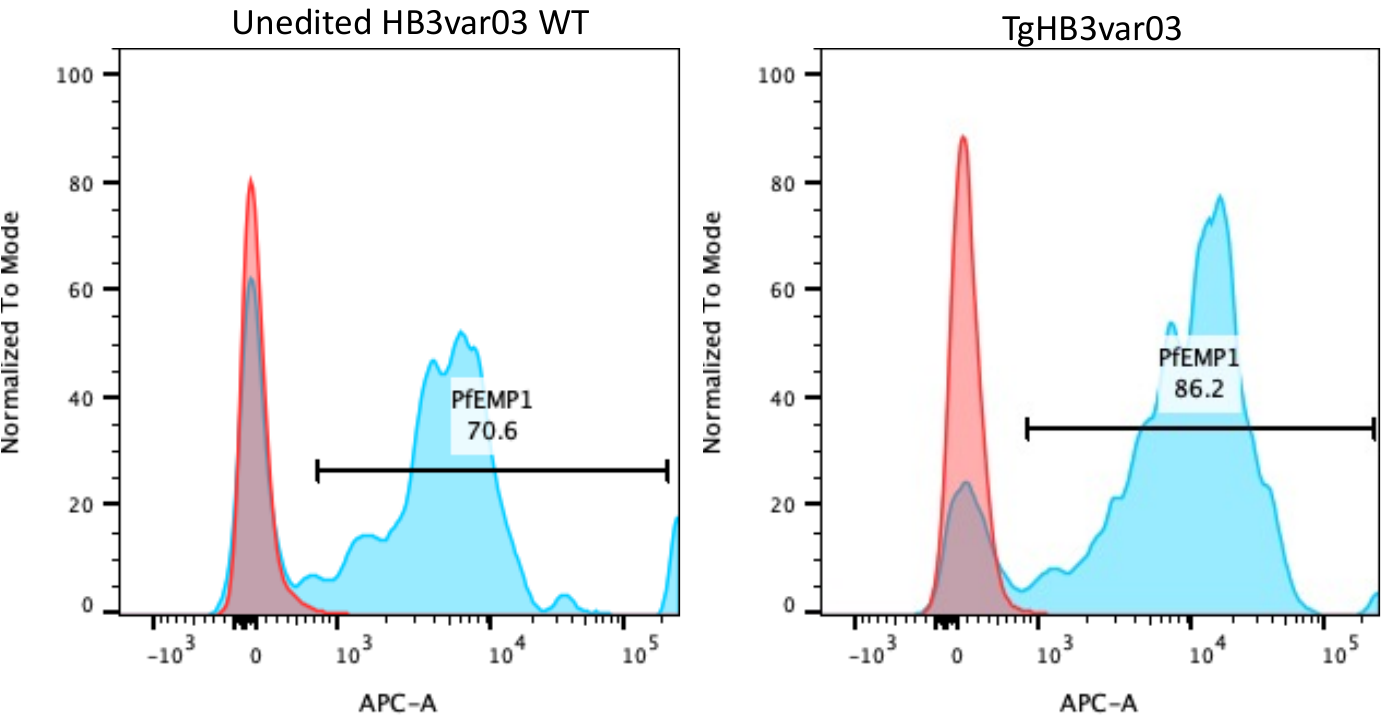

TgIT4var19-P1

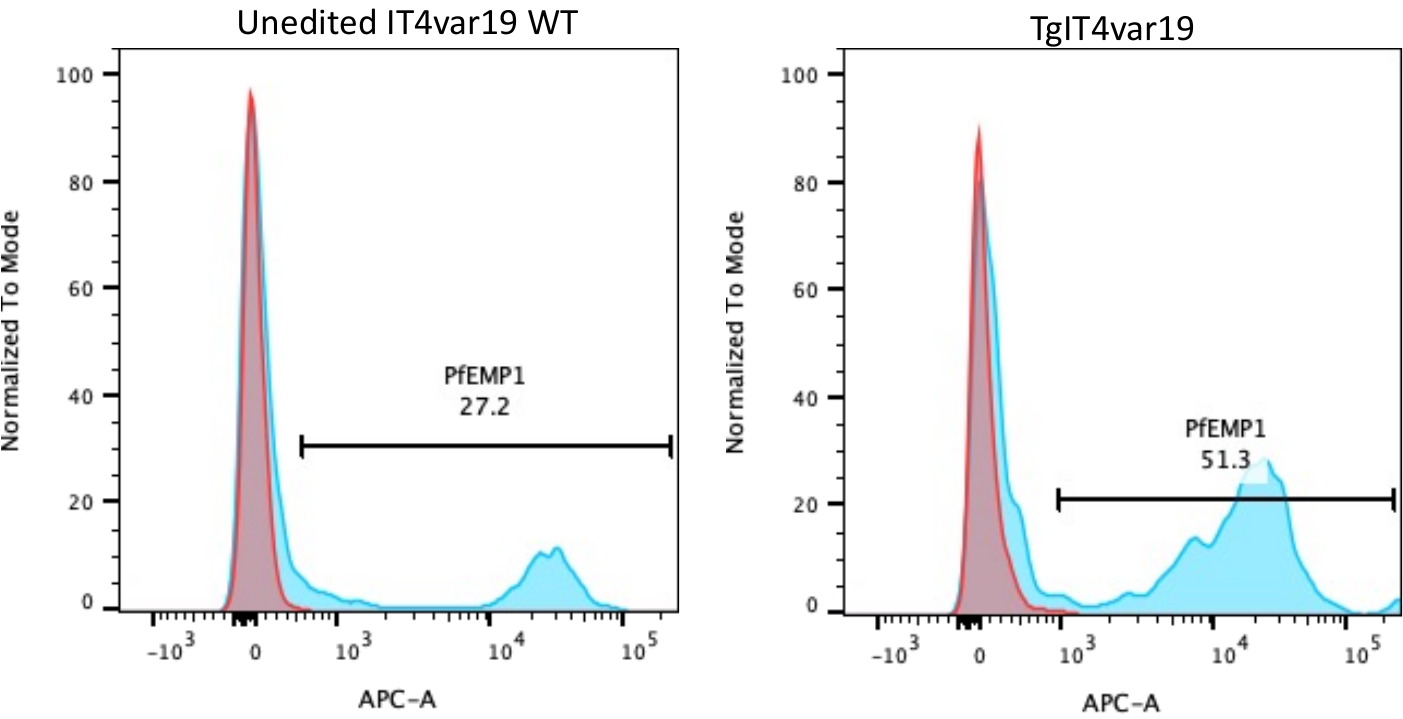

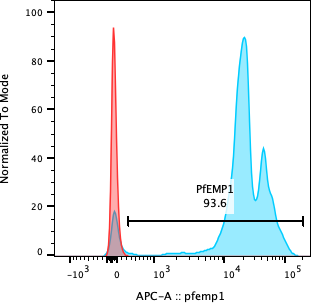

**C**

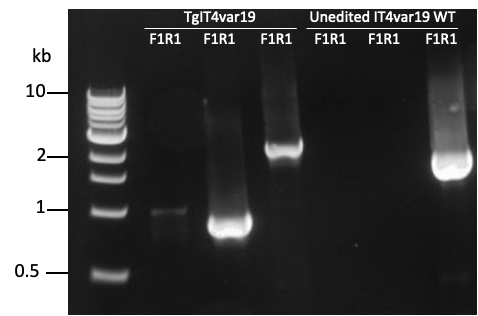

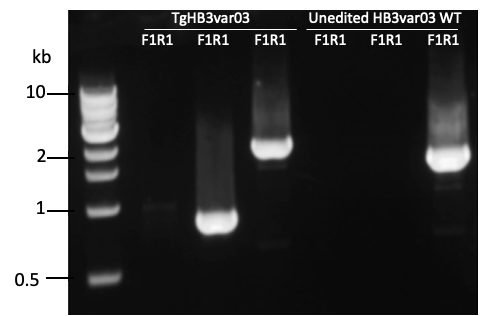

**A**

**B**

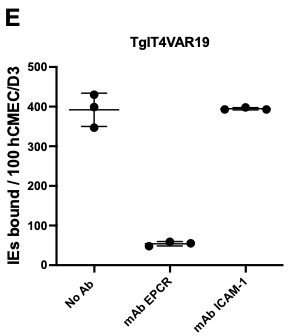

**Supplementary Figure 1: Integration PCR and flow cytometry analysis of PfEMP1 surface expression in transgenic IT4VAR19- and HB3VAR03-expressing parasites.** **(A-B)** Integration PCR confirming successful integration of *bsd*-2A at the targeted loci in TgIT4VAR19 (A) and TgHB3VAR03 (B) bulk cultures. **(C-D)** Flow cytometry histograms of mature infected erythrocytes (IEs) stained with non-immunised rabbit IgG (red, negative control) or variant-specific anti-NTS-DBLα IgG (blue). The percentage of mature live IEs expressing surface PfEMP1 is shown. Surface expression of PfEMP1 in TgIT4Var19 was increased by one round of panning (P1) on human brain endothelial cells (HBEC). Unedited HB3var03 was panned 5x on human brain endothelial cells (HBEC) over two months to achieve 70% surface positive IEs, whereas the Tg line expressed 86% positive IEs without panning. **(E)** As expected (Azasi et al PNAS 2018), binding of TgIT4VAR19-P1 to an HBEC line (hCMEC/D3) was inhibited by pre-incubation of the HBEC with a monoclonal antibody (mAb) to EPCR, whereas a mAb to ICAM-1 had no effect.

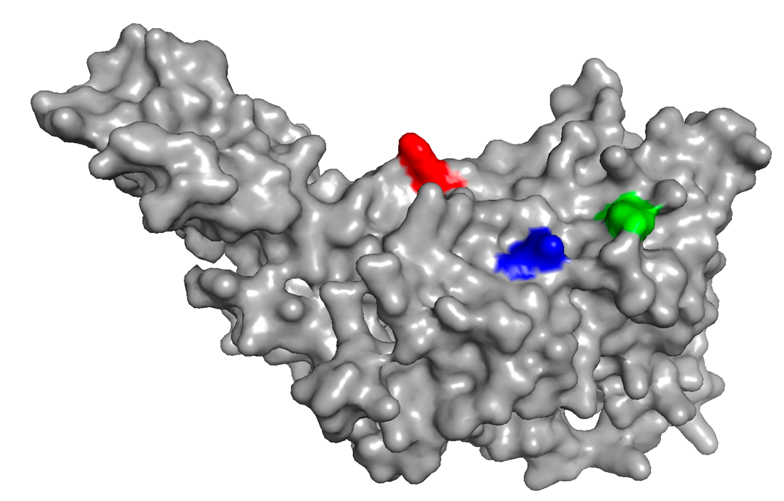

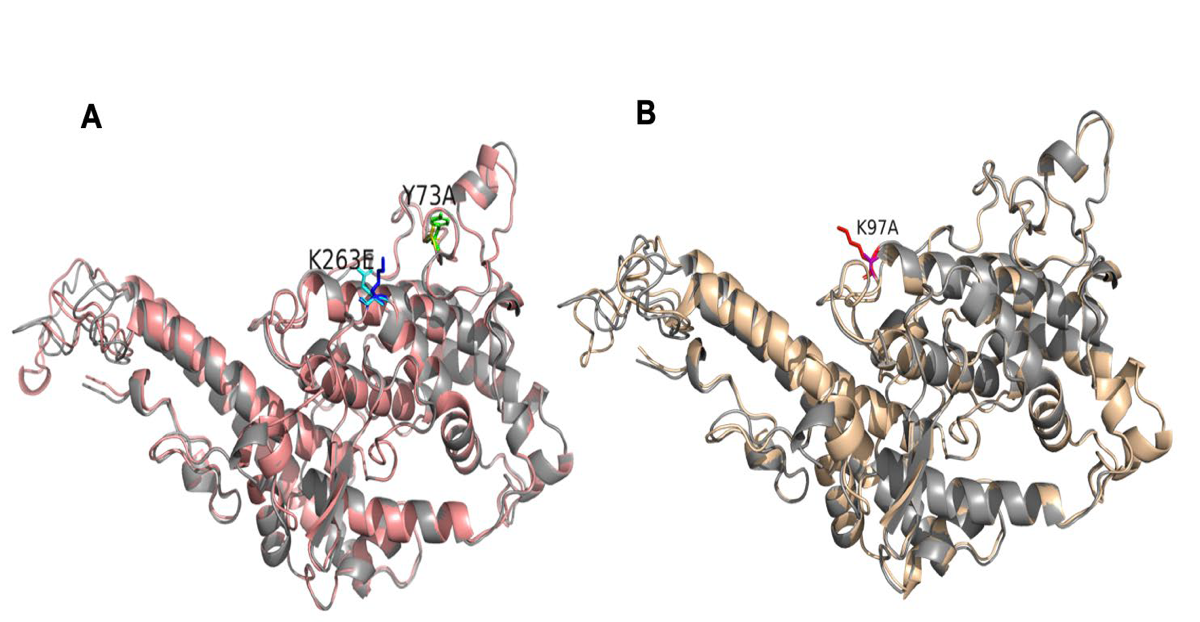

**A**

**B**

**Supplementary Figure 2: Structural model of the IT4VAR60 DBLα domain (A)** Surface model highlighting residues implicated in erythrocyte binding (Angeletti et al., 2015). Y73 (green); K97 (red), and K263 (blue). **(B)** Left**:** Alignment of IT4VAR60 DBLα_Mut A (pink) with IT4VAR60 DBLα wild type (WT) (grey). Right: IT4VAR60 DBLα_Mut F (wheat) aligned to IT4VAR60 DBLα WT (grey). The amino acids changes are shown as sticks and labelled. These changes are predicted to have no effect on the overall structure of the domain. The model is based on the crystal structure of varO NTS-DBLα (PDB: 2yK0) (Vigan-Womas et al., 2012).
