## Supplementary information 1 for "CRISPR/Cas9 genome editing to generate single variant *Plasmodium falciparum* lines and enable reverse genetic studies of PfEMP1 function in live parasites"

**Supplementary Information Otoboh et al:**

Nucleotide sequence of plasmid Cas9_itvar60prom_bsd_2A_exon1(b)

AAGCTTGGGGGGATCCGCCTTAAAAACTTCATTATATTTAAAAATTATTTTATAGGAAATAATAAAAAAAAAAgcaccga

ctcggtgccactttttcaagttgataacggactagccttattttaacttgctaTTTCtagctctaaaacTTTGGTGCCAT

TCTTATAACAATATTATATACTTAATATGAAATATGTGCATATAGGAAAAATTATGCATTTTGGTTACTCTAATATTATA

TATATATATATATATATATATATATTATAATATATTATGTTATATATACATAACATATACATTTTTTAATAATAATTTAC

CCTTTATTTTTACATTATAAAAAATTATATTACAGTAAAAATAAAAGTTTATTATATTAATAGTTTTTTTTTTTTTTTTT

TTAATTTATGAAATATTTAAATATTTAAAATTTTTTAAATGAATAATTATATTTATAATTAGAAAAAAAAAAAAAAAAAA

AAAAAAAAAATATAGCTATTTATATAAATTTCTTTTATTTATCTGAACAAGCAAGAATTTTTTTTTATATTAAATTAGAA

TAAATTATTATTAGTTTATGTATATATTTTTTTTTTTTCATAGTATATAAATATTATATATATTGTACCTTTTTACAATA

TATTTCATATATAGAAGAGAAAAAAAAAAAAAGAAGATATTATTGTAAAACCTCAAGATGTGTAGAAATCCAAATGTCGG

ATCCTCTAGAGTCGACCTGCAGGCATGCTATTTGATGAATTAACTACACTTAAAATAATACAATTATTATTAAATTTTTT

TTTGATTTATTTATTAATTTTTAAACTTAATCATTTGTATTTGGGAGGAATTATATATATCTTTATAATTATTTTATTTT

TTTTTATTTTTTTATTTTTTTATTATTATTATTTTTTTTTATTTTTTTTTTTTACTGTATCAAAGAAAAACCTTTAAAAA

AAAAATTATAATTTCCCCATCTTACTATATTTTTAATACATACGTTTTAAGGAATTAAATTAGACAAAAGCTATATTATG

CTTTACATATAATTAGAATTTATAAACGTTTGGTTATTAGATATTTCATGTCTCAGTAAAGTCTTTCAATACATATGTAA

AAAAATATATATGAATACACATAAGTTGTTAATATATTTTATATGCATAAATGTATAAATATATATATATATATATATAT

ATGTATGTATGTATATGTGTGTATATGAAATTATTTCAATGTTTAATTTTTTAAATTTTAATTTTTTTTTTTTTTTTTTT

TTTTATTATGTATATTGATCTTTATTATTTAAATATTACTTTTTTCGTTTTTTCTTCTTTTTATTATTTTTTTTTTTTTT

TATATTTTATACAAATGGTAATTCAAATAAAAGGTATAAATTTATATTTAATTTTCTTTTATGGATAAATAAAAGAAAAA

TATAAATATATAAAAATATAAAAATATATATATGTATATTGGGGTGATGATAAAATGAAAGATAATATATATATATATAT

ATCTTTATTTTTTTTTTTTTGTAGACCCCATTGTGAGTACATAAATATATTATATAACTCGAGTTACTTTTTCTTTTTTG

CCTGGCCGGCCTTTTTCGTGGCCGCCGGCCTTTTGTCGCCTCCCAGCTGAGACAGGTCGATCCGTGTCTCGTACAGGCCG

GTGATGCTCTGGTGGATCAGGGTGGCGTCCAGCACCTCTTTGGTGCTGGTGTACCTCTTCCGGTCGATGGTGGTGTCAAA

GTACTTGAAGGCGGCAGGGGCTCCCAGATTGGTCAGGGTAAACAGGTGGATGATATTCTCGGCCTGCTCTCTGATGGGCT

TATCCCGGTGCTTGTTGTAGGCGGACAGCACTTTGTCCAGATTAGCGTCGGCCAGGATCACTCTCTTGGAGAACTCGCTG

ATCTGCTCGATGATCTCGTCCAGGTAGTGCTTGTGCTGTTCCACAAACAGCTGTTTCTGCTCATTATCCTCGGGGGAGCC

CTTCAGCTTCTCATAGTGGCTGGCCAGGTACAGGAAGTTCACATATTTGGAGGGCAGGGCCAGTTCGTTTCCCTTCTGCA

GTTCGCCGGCAGAGGCCAGCATTCTCTTCCGGCCGTTTTCCAGCTCGAACAGGGAGTACTTAGGCAGCTTGATGATCAGG

TCCTTTTTCACTTCTTTGTAGCCCTTGGCTTCCAGAAAGTCGATGGGATTCTTCTCGAAGCTGCTTCTTTCCATGATGGT

GATCCCCAGCAGCTCTTTCACACTCTTCAGTTTCTTGGACTTGCCCTTTTCCACTTTGGCCACCACCAGCACAGAATAGG

CCACGGTGGGGCTGTCGAAGCCGCCGTACTTCTTAGGGTCCCAGTCCTTCTTTCTGGCGATCAGCTTATCGCTGTTCCTC

TTGGGCAGGATAGACTCTTTGCTGAAGCCGCCTGTCTGCACCTCGGTCTTTTTCACGATATTCACTTGGGGCATGCTCAG

CACTTTCCGCACGGTGGCAAAATCCCGGCCCTTATCCCACACGATCTCCCCGGTTTCGCCGTTTGTCTCGATCAGAGGCC

GCTTCCGGATCTCGCCGTTGGCCAGGGTAATCTCGGTCTTGAAAAAGTTCATGATGTTGCTGTAGAAGAAGTACTTGGCG

GTAGCCTTGCCGATTTCCTGCTCGCTCTTGGCGATCATCTTCCGCACGTCGTACACCTTGTAGTCGCCGTACACGAACTC

GCTTTCCAGCTTAGGGTACTTTTTGATCAGGGCGGTTCCCACGACGGCGTTCAGGTAGGCGTCGTGGGCGTGGTGGTAGT

TGTTGATCTCGCGCACTTTGTAAAACTGGAAATCCTTCCGGAAATCGGACACCAGCTTGGACTTCAGGGTGATCACTTTC

ACTTCCCGGATCAGCTTGTCATTCTCGTCGTACTTAGTGTTCATCCGGGAGTCCAGGATCTGTGCCACGTGCTTTGTGAT

CTGCCGGGTTTCCACCAGCTGTCTCTTGATGAAGCCGGCCTTATCCAGTTCGCTCAGGCCGCCTCTCTCGGCCTTGGTCA

GATTGTCGAACTTTCTCTGGGTAATCAGCTTGGCGTTCAGCAGCTGCCGCCAGTAGTTCTTCATCTTCTTCACGACCTCT

TCGGAGGGCACGTTGTCGCTCTTGCCCCGGTTCTTGTCGCTTCTGGTCAGCACCTTGTTGTCGATGGAGTCGTCCTTCAG

AAAGCTCTGAGGCACGATATGGTCCACATCGTAGTCGGACAGCCGGTTGATGTCCAGTTCCTGGTCCACGTACATATCCC

GCCCATTCTGCAGGTAGTACAGGTACAGCTTCTCGTTCTGCAGCTGGGTGTTTTCCACGGGGTGTTCTTTCAGGATCTGG

CTGCCCAGCTCTTTGATGCCCTCTTCGATCCGCTTCATTCTCTCGCGGCTGTTCTTCTGTCCCTTCTGGGTGGTCTGGTT

CTCTCTGGCCATTTCGATCACGATGTTCTCGGGCTTGTGCCGGCCCATCACTTTCACGAGCTCGTCCACCACCTTCACTG

TCTGCAGGATGCCCTTCTTAATGGCGGGGCTGCCGGCCAGATTGGCAATGTGCTCGTGCAGGCTATCGCCCTGGCCGGAC

ACCTGGGCTTTCTGGATGTCCTCTTTAAAGGTCAGGCTGTCGTCGTGGATCAGCTGCATGAAGTTTCTGTTGGCGAAGCC

GTCGGACTTCAGGAAATCCAGGATTGTCTTGCCGGACTGCTTGTCCCGGATGCCGTTGATCAGCTTCCGGCTCAGCCTGC

CCCAGCCGGTGTATCTCCGCCGCTTCAGCTGCTTCATCACTTTGTCGTCGAACAGGTGGGCATAGGTTTTCAGCCGTTCC

TCGATCATCTCTCTGTCCTCAAACAGTGTCAGGGTCAGCACGATATCTTCCAGAATGTCCTCGTTTTCCTCATTGTCCAG

GAAGTCCTTGTCCTTGATAATTTTCAGCAGATCGTGGTATGTGCCCAGGGAGGCGTTGAACCGATCTTCCACGCCGGAGA

TTTCCACGGAGTCGAAGCACTCGATTTTCTTGAAGTAGTCCTCTTTCAGCTGCTTCACGGTCACTTTCCGGTTGGTCTTG

AACAGCAGGTCCACGATGGCCTTTTTCTGCTCGCCGCTCAGGAAGGCGGGCTTTCTCATTCCCTCGGTCACGTATTTCAC

TTTGGTCAGCTCGTTATACACGGTGAAGTACTCGTACAGCAGGCTGTGCTTGGGCAGCACCTTCTCGTTGGGCAGGTTCT

TATCGAAGTTGGTCATCCGCTCGATGAAGCTCTGGGCGGAAGCGCCCTTGTCCACCACTTCCTCGAAGTTCCAGGGGGTG

ATGGTTTCCTCGCTCTTTCTGGTCATCCAGGCGAATCTGCTGTTTCCCCTGGCCAGAGGGCCCACGTAGTAGGGGATGCG

GAAGGTCAGGATCTTCTCGATCTTTTCCCGGTTGTCCTTCAGGAATGGGTAAAAATCTTCCTGCCGCCGCAGAATGGCGT

GCAGCTCTCCCAGGTGGATCTGGTGGGGGATGCTGCCGTTGTCGAAGGTCCGCTGCTTCCGCAGCAGGTCCTCTCTGTTC

AGCTTCACGAGCAGTTCCTCGGTGCCGTCCATCTTTTCCAGGATGGGCTTGATGAACTTGTAGAACTCTTCCTGGCTGGC

TCCGCCGTCAATGTAGCCGGCGTAGCCGTTCTTGCTCTGGTCGAAGAAAATCTCTTTGTACTTCTCAGGCAGCTGCTGCC

GCACGAGAGCTTTCAGCAGGGTCAGGTCCTGGTGGTGCTCGTCGTATCTCTTGATCATAGAGGCGCTCAGGGGGGCCTTG

GTGATCTCGGTGTTCACTCTCAGGATGTCGCTCAGCAGGATGGCGTCGGACAGGTTCTTGGCGGCCAGAAACAGGTCGGC

GTACTGGTCGCCGATCTGGGCCAGCAGGTTGTCCAGGTCGTCGTCGTAGGTGTCCTTGCTCAGCTGCAGTTTGGCATCCT

CGGCCAGGTCGAAGTTGCTCTTGAAGTTGGGGGTCAGGCCCAGGCTCAGGGCAATCAGGTTTCCGAACAGGCCATTCTTC

TTCTCGCCGGGCAGCTGGGCGATCAGATTTTCCAGCCGTCTGCTCTTGCTCAGTCTGGCAGACAGGATGGCCTTGGCGTC

CACGCCGCTGGCGTTGATGGGGTTTTCCTCGAACAGCTGGTTGTAGGTCTGCACCAGCTGGATGAACAGCTTGTCCACGT

CGCTGTTGTCGGGGTTCAGGTCGCCCTCGATCAGGAAGTGGCCCCGGAACTTGATCATGTGGGCCAGGGCCAGATAGATC

AGCCGCAGGTCGGCCTTGTCGGTGCTGTCCACCAGTTTCTTTCTCAGGTGGTAGATGGTGGGGTACTTCTCGTGGTAGGC

CACCTCGTCCACGATGTTGCCGAAGATGGGGTGCCGCTCGTGCTTCTTATCCTCTTCCACCAGGAAGGACTCTTCCAGTC

TGTGGAAGAAGCTGTCGTCCACCTTGGCCATCTCGTTGCTGAAGATCTCTTGCAGATAGCAGATCCGGTTCTTCCGTCTG

GTGTATCTTCTTCTGGCGGTTCTCTTCAGCCGGGTGGCCTCGGCTGTTTCGCCGCTGTCGAACAGCAGGGCTCCGATCAG

GTTCTTCTTGATGCTGTGCCGGTCGGTGTTGCCCAGCACCTTGAATTTCTTGCTGGGCACCTTGTACTCGTCGGTGATCA

CGGCCCAGCCCACAGAGTTGGTGCCGATGTCCAGGCCGATGCTGTACTTCTTGTCGGCTGCTGGGACTCCGTGGATACCG

ACCTTCCGCTTCTTCTTTGGGGCCATCTTATCGTCATCGTCTTTGTAATCAATATCATGATCCTTGTAGTCTCCGTCGTG

GTCCTTATAGTCCATcctaggTGATATATTTCTATTAGGTATTTATTATTATAAAATATAAATCTTGAATGATAATAAAT

AAAATATTAGTTATTCCTTTTCTAGTTTAAAATATACATATTATAAATATATATATATATATATATATTTTTATTGTGAC

AAGAATATATAATTATAAATTATATTATTTATTTTTGTATTTTTTTTTTTTTTTTTTTTTTTTTCTTTTTTTGTTTTATT

TTTCTTTTTTTTTATAAATATTATTTTTTTCTTTTATCATGCACATTGGAATAATACATTAATATATATATATATATTAT

ATTATACATATATTGAATAATGTTTATAAAAAATGCATAACTTATATGAATATAATTTTTTTTAAATATGACAAAAAGAA

AAAAAAAAAAAACCAAAAAAAATTAAAATTGAAATGAAATATATAAATATATTATTTATATATATTATACATTGTTTAAT

ACTACTACATGTATATATATATATTATATATATATATATATATCAATTTTTTCAAAAATAAATTAATATAAAAAGAGGGG

AAAAAAAAAAAAAAAAAAAAAAAAAGATAATTAAGTAAGCATTTAAAAATATATAAATTGATAATATATAAAATTAATCA

CATATAAACTAATATAATTTATAAAATAAGGAAAATAAAATATTACCATAAAATAAAAATAAAAATAAAAAAAAAAAAAA

AAAACACCTTTTTTTATATATATTAATATATAATTATCTCTTAGAAAAAATATTGTATAATTATATATGTAATGATTTAT

ATAAAAAAATAAAATTATACAAGTATATATTTTGTTTCTATAAATTGATATCTTAATTATTTATTATTAGAAATAGATAT

TTTTATAATAAACCAATAGATAAAATTTGTAGAGAAAAAAAAATAAAAATAAAAATAAAAATAATATAATATATAATAAA

ATAAAATAATATTATATAAATATATTTTAATTTTTTTTACAAAATGGTTGGGGgcgccaccggtTATGTAATAATAATAT

GATCATAATATTATAATAAAACTTATAAAAAAAATATTAAATATTTCATAAATGATTATTATTTATAATAAGATAGATTC

TTTAATTATTTTAAAATTGTATATTTTTTATGTATATTAATTTATTAATATTAATAAGAATATTATAAAAATTCTATTTT

ATTATTTAATGATATTACCCTAATAAAAATAATATAATTATATTACAATATAAATTTATATATATATATATATTTATATA

TTTAAAGAATATTTTATTTTTCAATAAGAACCTTCATTTTAAATTAACATCAAATTATATATATGTATATATACTTCTTA

GTATTATTAATTAAAATACGGAATAATATATAATATATATAAAATGGCAAAACTTTCCTATAGAAAAAAATATTCCATTT

ATTATATTTGTTGTAGGTAATTCTTATTACCGTTTCCTTCTGTTCGTAATGTATATTGGTATGTACTTTATTTTTGCAAT

TTAATTATATGTAAAAAAACGTTAGTACACCATATATATATTATAGTTATAAGAATGCATGCCAAGCCTTTGTCTCAAGA

AGAATCCACCCTCATTGAAAGAGCAACGGCTACAATCAACAGCATCCCCATCTCTGAAGACTACAGCGTCGCCAGCGCAG

CTCTCTCTAGCGACGGCCGCATCTTCACTGGTGTCAATGTATATCATTTTACTGGGGGACCTTGTGCAGAACTCGTGGTG

CTGGGCACTGCTGCTGCTGCGGCAGCTGGCAACCTGACTTGTATCGTCGCGATCGGAAATGAGAACAGGGGCATCTTGAG

CCCCTGCGGACGGTGCCGACAGGTGCTTCTCGATCTGCATCCTGGGATCAAAGCCATAGTGAAGGACAGTGATGGACAGC

CGACGGCAGTTGGGATTCGTGAATTGCTGCCCTCTGGTTATGTGTGGGAGGGCgCTAGcGGCAGTGGAGAGGGCAGAGGA

AGTCTGCTAACATGCGGTGACGTCGAGGAGAATCCTGGCCCAaagcttATGGCACCAAAGGGTAGAAGTACAAATGAAAT

TGAACTTAGCGCAAGAGATGTTTTGGAAAATATTGGAATAGGAATATATAATCAGGAAAAAATAAAAAAGAATCCATATG

AACAACAATTGAAAGGCACATTATCAAACGCCCGATTTCATGATGGCTTGCACAAGGCAGCTGATTTGGGGGTAATACCT

GGTCCTTCACATTTTTCTCAGCTTTATTACAAAAAGCATACTAATAACACAAAATATTATAAGGATGATAGGCATCCTTG

TCATGGTAGACAAGGAAAACGTTTTGATGAAGGTCAAAAATTTGAATGTGGTAATGATAAAATAATTGGTAATAGCGATA

AATATGGATCCTGTGCTCCACCTAGAAGAAGACATATATGTGATCAAAATTTAGAATTCTTAGATAACAATCATACTGAT

ACTATTCATGATGTATTGGGAAATGTGTTGGTCACAGCAAAATATGAAGGTGAATCTATTGTTAATGATCATCCAGATAA

AAAGAACAATGGTAATAAATCAGGTATATGTACTTCTCTTGCACGAAGTTTTGCCGATATAGGTGATATTGTAAGAGGAA

GAGATATGgtcgacaCTAGTACCGGTACGCGTgacgtCAGGTGGCACTTTTCGGGGAAATGTGCGCGGAACCCCTATTTG

TTTATTTTTCTAAATACATTCAAATATGTATCCGCTCATGAGACAATAACCCTGATAAATGCTTCAATAATATTGAAAAA

GGAAGAGTATGAGTATTCAACATTTCCGTGTCGCCCTTATTCCCTTTTTTGCGGCATTTTGCCTTCCTGTTTTTGCTCAC

CCAGAAACGCTGGTGAAAGTAAAAGATGCTGAAGATCAGTTGGGTGCACGAGTGGGTTACATCGAACTGGATCTCAACAG

CGGTAAGATCCTTGAGAGTTTTCGCCCCGAAGAACGTTTTCCAATGATGAGCACTTTTAAAGTTCTGCTATGTGGCGCGG

TATTATCCCGTATTGACGCCGGGCAAGAGCAACTCGGTCGCCGCATACACTATTCTCAGAATGACTTGGTTGAGTACTCA

CCAGTCACAGAAAAGCATCTTACGGATGGCATGACAGTAAGAGAATTATGCAGTGCTGCCATAACCATGAGTGATAACAC

TGCGGCCAACTTACTTCTGACAACGATCGGAGGACCGAAGGAGCTAACCGCTTTTTTGCACAACATGGGGGATCATGTAA

CTCGCCTTGATCGTTGGGAACCGGAGCTGAATGAAGCCATACCAAACGACGAGCGTGACACCACGATGCCTGTAGCAATG

CCAACAACGTTGCGCAAACTATTAACTGGCGAACTACTTACTCTAGCTTCCCGGCAACAATTAATAGACTGGATGGAGGC

GGATAAAGTTGCAGGACCACTTCTGCGCTCGGCCCTTCCGGCTGGCTGGTTTATTGCTGATAAATCTGGAGCCGGTGAGC

GTGGGTCTCGCGGTATCATTGCAGCACTGGGGCCAGATGGTAAGCCCTCCCGTATCGTAGTTATCTACACGACGGGGAGT

CAGGCAACTATGGATGAACGAAATAGACAGATCGCTGAGATAGGTGCCTCACTGATTAAGCATTGGTAACTGTCAGACCA

AGTTTACTCATATATACTTTAGATTGATTTAAAACTTCATTTTTAATTTAAAAGGATCTAGGTGAAGATCCTTTTTGATA

ATCTCATGACCAAAATCCCTTAACGTGAGTTTTCGTTCCACTGAGCGTCAGACCCCGTAGAAAAGATCAAAGGATCTTCT

TGAGATCCTTTTTTTCTGCGCGTAATCTGCTGCTTGCAAACAAAAAAACCACCGCTACCAGCGGTGGTTTGTTTGCCGGA

TCAAGAGCTACCAACTCTTTTTCCGAAGGTAACTGGCTTCAGCAGAGCGCAGATACCAAATACTGTCCTTCTAGTGTAGC

CGTAGTTAGGCCACCACTTCAAGAACTCTGTAGCACCGCCTACATACCTCGCTCTGCTAATCCTGTTACCAGTGGCTGCT

GCCAGTGGCGATAAGTCGTGTCTTACCGGGTTGGACTCAAGACGATAGTTACCGGATAAGGCGCAGCGGTCGGGCTGAAC

GGGGGGTTCGTGCACACAGCCCAGCTTGGAGCGAACGACCTACACCGAACTGAGATACCTACAGCGTGAGCTATGAGAAA

GCGCCACGCTTCCCGAAGGGAGAAAGGCGGACAGGTATCCGGTAAGCGGCAGGGTCGGAACAGGAGAGCGCACGAGGGAG

CTTCCAGGGGGAAACGCCTGGTATCTTTATAGTCCTGTCGGGTTTCGCCACCTCTGACTTGAGCGTCGATTTTTGTGATG

CTCGTCAGGGGGGCGGAGCCTATCGAAAAACGCCAGCAACGCGGCCTTTTTACGGTTCCTGGCCTTTTGCTGGCCTTTTG

CTCACATGTTCTTTCCTGCGTTATCCCCTGATTCTGTGGATAACCGTATTACCGCCTTTGAGTGAGCTGATACCGCTCGC

CGCAGCCGAACGACCGAGCGCAGCGAGTCAGTGAGCGAGGAAGCGGAAGAGCGCCCAATACGCAAACCGCCTCTCCCCGC

GCGTTGGCCGATTCATTAATGCAGCTGGCACGACAGGTTTCCCGACTGGAAAGCGGGCAGTGAGCGCAACGCAATTAATG

TGAGTTAGCTCACTCATTAGGCACCCCAGGCTTTACACTTTATGCTTCCGGCTCGTATGTTGTGTGGAATTGTGAGCGGA

TAACAATTTCACACAGGAAACAGCTATGACCATGATTACGCCAAGCTATTTAGGTGACACTATAGAATACTC

Nucleotide sequence of plasmid Cas9_itvar19prom_bsd_2A_exonI

AAGCTTGGGGGGATCCGCCTTAAAAACTTCATTATATTTAAAAATTATTTTATAGGAAATAATAAAAAAAAAAgcaccga

ctcggtgccactttttcaagttgataacggactagccttattttaacttgctaTTTCtagctctaaaacCCTTGGGCCCC

ATTTTTGTAAATATTATATACTTAATATGAAATATGTGCATATAGGAAAAATTATGCATTTTGGTTACTCTAATATTATA

TATATATATATATATATATATATATTATAATATATTATGTTATATATACATAACATATACATTTTTTAATAATAATTTAC

CCTTTATTTTTACATTATAAAAAATTATATTACAGTAAAAATAAAAGTTTATTATATTAATAGTTTTTTTTTTTTTTTTT

TTAATTTATGAAATATTTAAATATTTAAAATTTTTTAAATGAATAATTATATTTATAATTAGAAAAAAAAAAAAAAAAAA

AAAAAAAAAATATAGCTATTTATATAAATTTCTTTTATTTATCTGAACAAGCAAGAATTTTTTTTTATATTAAATTAGAA

TAAATTATTATTAGTTTATGTATATATTTTTTTTTTTTCATAGTATATAAATATTATATATATTGTACCTTTTTACAATA

TATTTCATATATAGAAGAGAAAAAAAAAAAAAGAAGATATTATTGTAAAACCTCAAGATGTGTAGAAATCCAAATGTCGG

ATCCTCTAGAGTCGACCTGCAGGCATGCTATTTGATGAATTAACTACACTTAAAATAATACAATTATTATTAAATTTTTT

TTTGATTTATTTATTAATTTTTAAACTTAATCATTTGTATTTGGGAGGAATTATATATATCTTTATAATTATTTTATTTT

TTTTTATTTTTTTATTTTTTTATTATTATTATTTTTTTTTATTTTTTTTTTTTACTGTATCAAAGAAAAACCTTTAAAAA

AAAAATTATAATTTCCCCATCTTACTATATTTTTAATACATACGTTTTAAGGAATTAAATTAGACAAAAGCTATATTATG

CTTTACATATAATTAGAATTTATAAACGTTTGGTTATTAGATATTTCATGTCTCAGTAAAGTCTTTCAATACATATGTAA

AAAAATATATATGAATACACATAAGTTGTTAATATATTTTATATGCATAAATGTATAAATATATATATATATATATATAT

ATGTATGTATGTATATGTGTGTATATGAAATTATTTCAATGTTTAATTTTTTAAATTTTAATTTTTTTTTTTTTTTTTTT

TTTTATTATGTATATTGATCTTTATTATTTAAATATTACTTTTTTCGTTTTTTCTTCTTTTTATTATTTTTTTTTTTTTT

TATATTTTATACAAATGGTAATTCAAATAAAAGGTATAAATTTATATTTAATTTTCTTTTATGGATAAATAAAAGAAAAA

TATAAATATATAAAAATATAAAAATATATATATGTATATTGGGGTGATGATAAAATGAAAGATAATATATATATATATAT

ATCTTTATTTTTTTTTTTTTGTAGACCCCATTGTGAGTACATAAATATATTATATAACTCGAGTTACTTTTTCTTTTTTG

CCTGGCCGGCCTTTTTCGTGGCCGCCGGCCTTTTGTCGCCTCCCAGCTGAGACAGGTCGATCCGTGTCTCGTACAGGCCG

GTGATGCTCTGGTGGATCAGGGTGGCGTCCAGCACCTCTTTGGTGCTGGTGTACCTCTTCCGGTCGATGGTGGTGTCAAA

GTACTTGAAGGCGGCAGGGGCTCCCAGATTGGTCAGGGTAAACAGGTGGATGATATTCTCGGCCTGCTCTCTGATGGGCT

TATCCCGGTGCTTGTTGTAGGCGGACAGCACTTTGTCCAGATTAGCGTCGGCCAGGATCACTCTCTTGGAGAACTCGCTG

ATCTGCTCGATGATCTCGTCCAGGTAGTGCTTGTGCTGTTCCACAAACAGCTGTTTCTGCTCATTATCCTCGGGGGAGCC

CTTCAGCTTCTCATAGTGGCTGGCCAGGTACAGGAAGTTCACATATTTGGAGGGCAGGGCCAGTTCGTTTCCCTTCTGCA

GTTCGCCGGCAGAGGCCAGCATTCTCTTCCGGCCGTTTTCCAGCTCGAACAGGGAGTACTTAGGCAGCTTGATGATCAGG

TCCTTTTTCACTTCTTTGTAGCCCTTGGCTTCCAGAAAGTCGATGGGATTCTTCTCGAAGCTGCTTCTTTCCATGATGGT

GATCCCCAGCAGCTCTTTCACACTCTTCAGTTTCTTGGACTTGCCCTTTTCCACTTTGGCCACCACCAGCACAGAATAGG

CCACGGTGGGGCTGTCGAAGCCGCCGTACTTCTTAGGGTCCCAGTCCTTCTTTCTGGCGATCAGCTTATCGCTGTTCCTC

TTGGGCAGGATAGACTCTTTGCTGAAGCCGCCTGTCTGCACCTCGGTCTTTTTCACGATATTCACTTGGGGCATGCTCAG

CACTTTCCGCACGGTGGCAAAATCCCGGCCCTTATCCCACACGATCTCCCCGGTTTCGCCGTTTGTCTCGATCAGAGGCC

GCTTCCGGATCTCGCCGTTGGCCAGGGTAATCTCGGTCTTGAAAAAGTTCATGATGTTGCTGTAGAAGAAGTACTTGGCG

GTAGCCTTGCCGATTTCCTGCTCGCTCTTGGCGATCATCTTCCGCACGTCGTACACCTTGTAGTCGCCGTACACGAACTC

GCTTTCCAGCTTAGGGTACTTTTTGATCAGGGCGGTTCCCACGACGGCGTTCAGGTAGGCGTCGTGGGCGTGGTGGTAGT

TGTTGATCTCGCGCACTTTGTAAAACTGGAAATCCTTCCGGAAATCGGACACCAGCTTGGACTTCAGGGTGATCACTTTC

ACTTCCCGGATCAGCTTGTCATTCTCGTCGTACTTAGTGTTCATCCGGGAGTCCAGGATCTGTGCCACGTGCTTTGTGAT

CTGCCGGGTTTCCACCAGCTGTCTCTTGATGAAGCCGGCCTTATCCAGTTCGCTCAGGCCGCCTCTCTCGGCCTTGGTCA

GATTGTCGAACTTTCTCTGGGTAATCAGCTTGGCGTTCAGCAGCTGCCGCCAGTAGTTCTTCATCTTCTTCACGACCTCT

TCGGAGGGCACGTTGTCGCTCTTGCCCCGGTTCTTGTCGCTTCTGGTCAGCACCTTGTTGTCGATGGAGTCGTCCTTCAG

AAAGCTCTGAGGCACGATATGGTCCACATCGTAGTCGGACAGCCGGTTGATGTCCAGTTCCTGGTCCACGTACATATCCC

GCCCATTCTGCAGGTAGTACAGGTACAGCTTCTCGTTCTGCAGCTGGGTGTTTTCCACGGGGTGTTCTTTCAGGATCTGG

CTGCCCAGCTCTTTGATGCCCTCTTCGATCCGCTTCATTCTCTCGCGGCTGTTCTTCTGTCCCTTCTGGGTGGTCTGGTT

CTCTCTGGCCATTTCGATCACGATGTTCTCGGGCTTGTGCCGGCCCATCACTTTCACGAGCTCGTCCACCACCTTCACTG

TCTGCAGGATGCCCTTCTTAATGGCGGGGCTGCCGGCCAGATTGGCAATGTGCTCGTGCAGGCTATCGCCCTGGCCGGAC

ACCTGGGCTTTCTGGATGTCCTCTTTAAAGGTCAGGCTGTCGTCGTGGATCAGCTGCATGAAGTTTCTGTTGGCGAAGCC

GTCGGACTTCAGGAAATCCAGGATTGTCTTGCCGGACTGCTTGTCCCGGATGCCGTTGATCAGCTTCCGGCTCAGCCTGC

CCCAGCCGGTGTATCTCCGCCGCTTCAGCTGCTTCATCACTTTGTCGTCGAACAGGTGGGCATAGGTTTTCAGCCGTTCC

TCGATCATCTCTCTGTCCTCAAACAGTGTCAGGGTCAGCACGATATCTTCCAGAATGTCCTCGTTTTCCTCATTGTCCAG

GAAGTCCTTGTCCTTGATAATTTTCAGCAGATCGTGGTATGTGCCCAGGGAGGCGTTGAACCGATCTTCCACGCCGGAGA

TTTCCACGGAGTCGAAGCACTCGATTTTCTTGAAGTAGTCCTCTTTCAGCTGCTTCACGGTCACTTTCCGGTTGGTCTTG

AACAGCAGGTCCACGATGGCCTTTTTCTGCTCGCCGCTCAGGAAGGCGGGCTTTCTCATTCCCTCGGTCACGTATTTCAC

TTTGGTCAGCTCGTTATACACGGTGAAGTACTCGTACAGCAGGCTGTGCTTGGGCAGCACCTTCTCGTTGGGCAGGTTCT

TATCGAAGTTGGTCATCCGCTCGATGAAGCTCTGGGCGGAAGCGCCCTTGTCCACCACTTCCTCGAAGTTCCAGGGGGTG

ATGGTTTCCTCGCTCTTTCTGGTCATCCAGGCGAATCTGCTGTTTCCCCTGGCCAGAGGGCCCACGTAGTAGGGGATGCG

GAAGGTCAGGATCTTCTCGATCTTTTCCCGGTTGTCCTTCAGGAATGGGTAAAAATCTTCCTGCCGCCGCAGAATGGCGT

GCAGCTCTCCCAGGTGGATCTGGTGGGGGATGCTGCCGTTGTCGAAGGTCCGCTGCTTCCGCAGCAGGTCCTCTCTGTTC

AGCTTCACGAGCAGTTCCTCGGTGCCGTCCATCTTTTCCAGGATGGGCTTGATGAACTTGTAGAACTCTTCCTGGCTGGC

TCCGCCGTCAATGTAGCCGGCGTAGCCGTTCTTGCTCTGGTCGAAGAAAATCTCTTTGTACTTCTCAGGCAGCTGCTGCC

GCACGAGAGCTTTCAGCAGGGTCAGGTCCTGGTGGTGCTCGTCGTATCTCTTGATCATAGAGGCGCTCAGGGGGGCCTTG

GTGATCTCGGTGTTCACTCTCAGGATGTCGCTCAGCAGGATGGCGTCGGACAGGTTCTTGGCGGCCAGAAACAGGTCGGC

GTACTGGTCGCCGATCTGGGCCAGCAGGTTGTCCAGGTCGTCGTCGTAGGTGTCCTTGCTCAGCTGCAGTTTGGCATCCT

CGGCCAGGTCGAAGTTGCTCTTGAAGTTGGGGGTCAGGCCCAGGCTCAGGGCAATCAGGTTTCCGAACAGGCCATTCTTC

TTCTCGCCGGGCAGCTGGGCGATCAGATTTTCCAGCCGTCTGCTCTTGCTCAGTCTGGCAGACAGGATGGCCTTGGCGTC

CACGCCGCTGGCGTTGATGGGGTTTTCCTCGAACAGCTGGTTGTAGGTCTGCACCAGCTGGATGAACAGCTTGTCCACGT

CGCTGTTGTCGGGGTTCAGGTCGCCCTCGATCAGGAAGTGGCCCCGGAACTTGATCATGTGGGCCAGGGCCAGATAGATC

AGCCGCAGGTCGGCCTTGTCGGTGCTGTCCACCAGTTTCTTTCTCAGGTGGTAGATGGTGGGGTACTTCTCGTGGTAGGC

CACCTCGTCCACGATGTTGCCGAAGATGGGGTGCCGCTCGTGCTTCTTATCCTCTTCCACCAGGAAGGACTCTTCCAGTC

TGTGGAAGAAGCTGTCGTCCACCTTGGCCATCTCGTTGCTGAAGATCTCTTGCAGATAGCAGATCCGGTTCTTCCGTCTG

GTGTATCTTCTTCTGGCGGTTCTCTTCAGCCGGGTGGCCTCGGCTGTTTCGCCGCTGTCGAACAGCAGGGCTCCGATCAG

GTTCTTCTTGATGCTGTGCCGGTCGGTGTTGCCCAGCACCTTGAATTTCTTGCTGGGCACCTTGTACTCGTCGGTGATCA

CGGCCCAGCCCACAGAGTTGGTGCCGATGTCCAGGCCGATGCTGTACTTCTTGTCGGCTGCTGGGACTCCGTGGATACCG

ACCTTCCGCTTCTTCTTTGGGGCCATCTTATCGTCATCGTCTTTGTAATCAATATCATGATCCTTGTAGTCTCCGTCGTG

GTCCTTATAGTCCATcctaggTGATATATTTCTATTAGGTATTTATTATTATAAAATATAAATCTTGAATGATAATAAAT

AAAATATTAGTTATTCCTTTTCTAGTTTAAAATATACATATTATAAATATATATATATATATATATATTTTTATTGTGAC

AAGAATATATAATTATAAATTATATTATTTATTTTTGTATTTTTTTTTTTTTTTTTTTTTTTTTCTTTTTTTGTTTTATT

TTTCTTTTTTTTTATAAATATTATTTTTTTCTTTTATCATGCACATTGGAATAATACATTAATATATATATATATATTAT

ATTATACATATATTGAATAATGTTTATAAAAAATGCATAACTTATATGAATATAATTTTTTTTAAATATGACAAAAAGAA

AAAAAAAAAAAACCAAAAAAAATTAAAATTGAAATGAAATATATAAATATATTATTTATATATATTATACATTGTTTAAT

ACTACTACATGTATATATATATATTATATATATATATATATATCAATTTTTTCAAAAATAAATTAATATAAAAAGAGGGG

AAAAAAAAAAAAAAAAAAAAAAAAAGATAATTAAGTAAGCATTTAAAAATATATAAATTGATAATATATAAAATTAATCA

CATATAAACTAATATAATTTATAAAATAAGGAAAATAAAATATTACCATAAAATAAAAATAAAAATAAAAAAAAAAAAAA

AAAACACCTTTTTTTATATATATTAATATATAATTATCTCTTAGAAAAAATATTGTATAATTATATATGTAATGATTTAT

ATAAAAAAATAAAATTATACAAGTATATATTTTGTTTCTATAAATTGATATCTTAATTATTTATTATTAGAAATAGATAT

TTTTATAATAAACCAATAGATAAAATTTGTAGAGAAAAAAAAATAAAAATAAAAATAAAAATAATATAATATATAATAAA

ATAAAATAATATTATATAAATATATTTTAATTTTTTTTACAAAATGGTTGGGGgcgccaccggtTATATTTAGGTTTATA

ATTAAATATAGAAACATGTTGTATGTTTTTATATGTATGTATTTTCGTATTTTTTTTTTTTCTCATATATAATTTTACAA

AATATAAAAAACATAAAAAAATAATATATAAAATTAAATATAAAAATAAAGGAATACATGAAATATAATATTTTTCATAA

AATGTAATTGTTGTTTTTTTTTTTGTTAGAATATTTAAATTTATTATACAAGTATTAATATATTTTTTTTTAAAAATATA

TATATAAAACTAATAATTATTATTATATACATATTAAATATTATTTATTAATATATATTATATATATATTATAATATTAC

AACTATTATAATTACTATATATATACAAATATATATATAATACTTATATATATATATATTCCAACACAATACTATTATTA

TTATTCTACCATATCACTATACTCCCATAACATACGCAATACGCCACCGCCACCGCCAACACTTACCATGTATGCCACGA

TATAAACCACGTATGTATGTATGACATCATGTAGTCGGGAAGAAGAATACAAAAATGCATGCCAAGCCTTTGTCTCAAGA

AGAATCCACCCTCATTGAAAGAGCAACGGCTACAATCAACAGCATCCCCATCTCTGAAGACTACAGCGTCGCCAGCGCAG

CTCTCTCTAGCGACGGCCGCATCTTCACTGGTGTCAATGTATATCATTTTACTGGGGGACCTTGTGCAGAACTCGTGGTG

CTGGGCACTGCTGCTGCTGCGGCAGCTGGCAACCTGACTTGTATCGTCGCGATCGGAAATGAGAACAGGGGCATCTTGAG

CCCCTGCGGACGGTGCCGACAGGTGCTTCTCGATCTGCATCCTGGGATCAAAGCCATAGTGAAGGACAGTGATGGACAGC

CGACGGCAGTTGGGATTCGTGAATTGCTGCCCTCTGGTTATGTGTGGGAGGGCgCTAGcGGCAGTGGAGAGGGCAGAGGA

AGTCTGCTAACATGCGGTGACGTCGAGGAGAATCCTGGCCCAaagcttATGGGGCCTAAAGCAGCTGTGACTGACTACAG

TGATGCCAAACATTTATTGGATAGCATAGGGGAAAAAGTGTACAAAGAAAAAGTGGAAAAAAAAGCTGTAGATTATAGAA

GCGCTTTGCAAGGACGTTTGTCAGAAGCATCATTTAAAGATCGTAAGAACATCGAGCGTGGAAAAGCCGAATTATGTAAA

CTTAATCATATATATCATACTAATGTTACGGATGGTTATGGTAGGGAGCATCCTTGTAAAGATAGATGGGACATTCGCTT

TTCTGATAAATATGGTGGTCAATGCACTAATAGTAAAATACATGGTAATGATGATAGTAATGGTAAAGACATTGGAGCCT

GCGCGCCGTTCAGACGATTACATCTATGTGACCAACATTTATCGCACATGAAAGCTGAAAAAATTAATTCTAAAGATAAT

TTGTTGTTAGAAGTGTGTCTTGCAGCACAATATGAAGGAGAATCATTAGTAGAAAAACATAAAGAATTTAAAAAAACACA

TAACGATTCCAATATATGTACTATATTGGCACGAAGTTTTGCAGATATAGGAGATATTATTAGAGGAAAAGATCTGTATC

TTGGTAATgtcgacaCTAGTACCGGTACGCGTgacgtCAGGTGGCACTTTTCGGGGAAATGTGCGCGGAACCCCTATTTG

TTTATTTTTCTAAATACATTCAAATATGTATCCGCTCATGAGACAATAACCCTGATAAATGCTTCAATAATATTGAAAAA

GGAAGAGTATGAGTATTCAACATTTCCGTGTCGCCCTTATTCCCTTTTTTGCGGCATTTTGCCTTCCTGTTTTTGCTCAC

CCAGAAACGCTGGTGAAAGTAAAAGATGCTGAAGATCAGTTGGGTGCACGAGTGGGTTACATCGAACTGGATCTCAACAG

CGGTAAGATCCTTGAGAGTTTTCGCCCCGAAGAACGTTTTCCAATGATGAGCACTTTTAAAGTTCTGCTATGTGGCGCGG

TATTATCCCGTATTGACGCCGGGCAAGAGCAACTCGGTCGCCGCATACACTATTCTCAGAATGACTTGGTTGAGTACTCA

CCAGTCACAGAAAAGCATCTTACGGATGGCATGACAGTAAGAGAATTATGCAGTGCTGCCATAACCATGAGTGATAACAC

TGCGGCCAACTTACTTCTGACAACGATCGGAGGACCGAAGGAGCTAACCGCTTTTTTGCACAACATGGGGGATCATGTAA

CTCGCCTTGATCGTTGGGAACCGGAGCTGAATGAAGCCATACCAAACGACGAGCGTGACACCACGATGCCTGTAGCAATG

CCAACAACGTTGCGCAAACTATTAACTGGCGAACTACTTACTCTAGCTTCCCGGCAACAATTAATAGACTGGATGGAGGC

GGATAAAGTTGCAGGACCACTTCTGCGCTCGGCCCTTCCGGCTGGCTGGTTTATTGCTGATAAATCTGGAGCCGGTGAGC

GTGGGTCTCGCGGTATCATTGCAGCACTGGGGCCAGATGGTAAGCCCTCCCGTATCGTAGTTATCTACACGACGGGGAGT

CAGGCAACTATGGATGAACGAAATAGACAGATCGCTGAGATAGGTGCCTCACTGATTAAGCATTGGTAACTGTCAGACCA

AGTTTACTCATATATACTTTAGATTGATTTAAAACTTCATTTTTAATTTAAAAGGATCTAGGTGAAGATCCTTTTTGATA

ATCTCATGACCAAAATCCCTTAACGTGAGTTTTCGTTCCACTGAGCGTCAGACCCCGTAGAAAAGATCAAAGGATCTTCT

TGAGATCCTTTTTTTCTGCGCGTAATCTGCTGCTTGCAAACAAAAAAACCACCGCTACCAGCGGTGGTTTGTTTGCCGGA

TCAAGAGCTACCAACTCTTTTTCCGAAGGTAACTGGCTTCAGCAGAGCGCAGATACCAAATACTGTCCTTCTAGTGTAGC

CGTAGTTAGGCCACCACTTCAAGAACTCTGTAGCACCGCCTACATACCTCGCTCTGCTAATCCTGTTACCAGTGGCTGCT

GCCAGTGGCGATAAGTCGTGTCTTACCGGGTTGGACTCAAGACGATAGTTACCGGATAAGGCGCAGCGGTCGGGCTGAAC

GGGGGGTTCGTGCACACAGCCCAGCTTGGAGCGAACGACCTACACCGAACTGAGATACCTACAGCGTGAGCTATGAGAAA

GCGCCACGCTTCCCGAAGGGAGAAAGGCGGACAGGTATCCGGTAAGCGGCAGGGTCGGAACAGGAGAGCGCACGAGGGAG

CTTCCAGGGGGAAACGCCTGGTATCTTTATAGTCCTGTCGGGTTTCGCCACCTCTGACTTGAGCGTCGATTTTTGTGATG

CTCGTCAGGGGGGCGGAGCCTATCGAAAAACGCCAGCAACGCGGCCTTTTTACGGTTCCTGGCCTTTTGCTGGCCTTTTG

CTCACATGTTCTTTCCTGCGTTATCCCCTGATTCTGTGGATAACCGTATTACCGCCTTTGAGTGAGCTGATACCGCTCGC

CGCAGCCGAACGACCGAGCGCAGCGAGTCAGTGAGCGAGGAAGCGGAAGAGCGCCCAATACGCAAACCGCCTCTCCCCGC

GCGTTGGCCGATTCATTAATGCAGCTGGCACGACAGGTTTCCCGACTGGAAAGCGGGCAGTGAGCGCAACGCAATTAATG

TGAGTTAGCTCACTCATTAGGCACCCCAGGCTTTACACTTTATGCTTCCGGCTCGTATGTTGTGTGGAATTGTGAGCGGA

TAACAATTTCACACAGGAAACAGCTATGACCATGATTACGCCAAGCTATTTAGGTGACACTATAGAATACTC

Nucleotide sequence of plasmid Cas9_hb3var03prom_bsd_2A_exonI

AAGCTTGGGGGGATCCGCCTTAAAAACTTCATTATATTTAAAAATTATTTTATAGGAAATAATAAAAAAAAAAgcaccga

ctcggtgccactttttcaagttgataacggactagccttattttaacttgctaTTTCtagctctaaaacATTTTGATAAA

AAAATATTTAATATTATATACTTAATATGAAATATGTGCATATAGGAAAAATTATGCATTTTGGTTACTCTAATATTATA

TATATATATATATATATATATATATTATAATATATTATGTTATATATACATAACATATACATTTTTTAATAATAATTTAC

CCTTTATTTTTACATTATAAAAAATTATATTACAGTAAAAATAAAAGTTTATTATATTAATAGTTTTTTTTTTTTTTTTT

TTAATTTATGAAATATTTAAATATTTAAAATTTTTTAAATGAATAATTATATTTATAATTAGAAAAAAAAAAAAAAAAAA

AAAAAAAAAATATAGCTATTTATATAAATTTCTTTTATTTATCTGAACAAGCAAGAATTTTTTTTTATATTAAATTAGAA

TAAATTATTATTAGTTTATGTATATATTTTTTTTTTTTCATAGTATATAAATATTATATATATTGTACCTTTTTACAATA

TATTTCATATATAGAAGAGAAAAAAAAAAAAAGAAGATATTATTGTAAAACCTCAAGATGTGTAGAAATCCAAATGTCGG

ATCCTCTAGAGTCGACCTGCAGGCATGCTATTTGATGAATTAACTACACTTAAAATAATACAATTATTATTAAATTTTTT

TTTGATTTATTTATTAATTTTTAAACTTAATCATTTGTATTTGGGAGGAATTATATATATCTTTATAATTATTTTATTTT

TTTTTATTTTTTTATTTTTTTATTATTATTATTTTTTTTTATTTTTTTTTTTTACTGTATCAAAGAAAAACCTTTAAAAA

AAAAATTATAATTTCCCCATCTTACTATATTTTTAATACATACGTTTTAAGGAATTAAATTAGACAAAAGCTATATTATG

CTTTACATATAATTAGAATTTATAAACGTTTGGTTATTAGATATTTCATGTCTCAGTAAAGTCTTTCAATACATATGTAA

AAAAATATATATGAATACACATAAGTTGTTAATATATTTTATATGCATAAATGTATAAATATATATATATATATATATAT

ATGTATGTATGTATATGTGTGTATATGAAATTATTTCAATGTTTAATTTTTTAAATTTTAATTTTTTTTTTTTTTTTTTT

TTTTATTATGTATATTGATCTTTATTATTTAAATATTACTTTTTTCGTTTTTTCTTCTTTTTATTATTTTTTTTTTTTTT

TATATTTTATACAAATGGTAATTCAAATAAAAGGTATAAATTTATATTTAATTTTCTTTTATGGATAAATAAAAGAAAAA

TATAAATATATAAAAATATAAAAATATATATATGTATATTGGGGTGATGATAAAATGAAAGATAATATATATATATATAT

ATCTTTATTTTTTTTTTTTTGTAGACCCCATTGTGAGTACATAAATATATTATATAACTCGAGTTACTTTTTCTTTTTTG

CCTGGCCGGCCTTTTTCGTGGCCGCCGGCCTTTTGTCGCCTCCCAGCTGAGACAGGTCGATCCGTGTCTCGTACAGGCCG

GTGATGCTCTGGTGGATCAGGGTGGCGTCCAGCACCTCTTTGGTGCTGGTGTACCTCTTCCGGTCGATGGTGGTGTCAAA

GTACTTGAAGGCGGCAGGGGCTCCCAGATTGGTCAGGGTAAACAGGTGGATGATATTCTCGGCCTGCTCTCTGATGGGCT

TATCCCGGTGCTTGTTGTAGGCGGACAGCACTTTGTCCAGATTAGCGTCGGCCAGGATCACTCTCTTGGAGAACTCGCTG

ATCTGCTCGATGATCTCGTCCAGGTAGTGCTTGTGCTGTTCCACAAACAGCTGTTTCTGCTCATTATCCTCGGGGGAGCC

CTTCAGCTTCTCATAGTGGCTGGCCAGGTACAGGAAGTTCACATATTTGGAGGGCAGGGCCAGTTCGTTTCCCTTCTGCA

GTTCGCCGGCAGAGGCCAGCATTCTCTTCCGGCCGTTTTCCAGCTCGAACAGGGAGTACTTAGGCAGCTTGATGATCAGG

TCCTTTTTCACTTCTTTGTAGCCCTTGGCTTCCAGAAAGTCGATGGGATTCTTCTCGAAGCTGCTTCTTTCCATGATGGT

GATCCCCAGCAGCTCTTTCACACTCTTCAGTTTCTTGGACTTGCCCTTTTCCACTTTGGCCACCACCAGCACAGAATAGG

CCACGGTGGGGCTGTCGAAGCCGCCGTACTTCTTAGGGTCCCAGTCCTTCTTTCTGGCGATCAGCTTATCGCTGTTCCTC

TTGGGCAGGATAGACTCTTTGCTGAAGCCGCCTGTCTGCACCTCGGTCTTTTTCACGATATTCACTTGGGGCATGCTCAG

CACTTTCCGCACGGTGGCAAAATCCCGGCCCTTATCCCACACGATCTCCCCGGTTTCGCCGTTTGTCTCGATCAGAGGCC

GCTTCCGGATCTCGCCGTTGGCCAGGGTAATCTCGGTCTTGAAAAAGTTCATGATGTTGCTGTAGAAGAAGTACTTGGCG

GTAGCCTTGCCGATTTCCTGCTCGCTCTTGGCGATCATCTTCCGCACGTCGTACACCTTGTAGTCGCCGTACACGAACTC

GCTTTCCAGCTTAGGGTACTTTTTGATCAGGGCGGTTCCCACGACGGCGTTCAGGTAGGCGTCGTGGGCGTGGTGGTAGT

TGTTGATCTCGCGCACTTTGTAAAACTGGAAATCCTTCCGGAAATCGGACACCAGCTTGGACTTCAGGGTGATCACTTTC

ACTTCCCGGATCAGCTTGTCATTCTCGTCGTACTTAGTGTTCATCCGGGAGTCCAGGATCTGTGCCACGTGCTTTGTGAT

CTGCCGGGTTTCCACCAGCTGTCTCTTGATGAAGCCGGCCTTATCCAGTTCGCTCAGGCCGCCTCTCTCGGCCTTGGTCA

GATTGTCGAACTTTCTCTGGGTAATCAGCTTGGCGTTCAGCAGCTGCCGCCAGTAGTTCTTCATCTTCTTCACGACCTCT

TCGGAGGGCACGTTGTCGCTCTTGCCCCGGTTCTTGTCGCTTCTGGTCAGCACCTTGTTGTCGATGGAGTCGTCCTTCAG

AAAGCTCTGAGGCACGATATGGTCCACATCGTAGTCGGACAGCCGGTTGATGTCCAGTTCCTGGTCCACGTACATATCCC

GCCCATTCTGCAGGTAGTACAGGTACAGCTTCTCGTTCTGCAGCTGGGTGTTTTCCACGGGGTGTTCTTTCAGGATCTGG

CTGCCCAGCTCTTTGATGCCCTCTTCGATCCGCTTCATTCTCTCGCGGCTGTTCTTCTGTCCCTTCTGGGTGGTCTGGTT

CTCTCTGGCCATTTCGATCACGATGTTCTCGGGCTTGTGCCGGCCCATCACTTTCACGAGCTCGTCCACCACCTTCACTG

TCTGCAGGATGCCCTTCTTAATGGCGGGGCTGCCGGCCAGATTGGCAATGTGCTCGTGCAGGCTATCGCCCTGGCCGGAC

ACCTGGGCTTTCTGGATGTCCTCTTTAAAGGTCAGGCTGTCGTCGTGGATCAGCTGCATGAAGTTTCTGTTGGCGAAGCC

GTCGGACTTCAGGAAATCCAGGATTGTCTTGCCGGACTGCTTGTCCCGGATGCCGTTGATCAGCTTCCGGCTCAGCCTGC

CCCAGCCGGTGTATCTCCGCCGCTTCAGCTGCTTCATCACTTTGTCGTCGAACAGGTGGGCATAGGTTTTCAGCCGTTCC

TCGATCATCTCTCTGTCCTCAAACAGTGTCAGGGTCAGCACGATATCTTCCAGAATGTCCTCGTTTTCCTCATTGTCCAG

GAAGTCCTTGTCCTTGATAATTTTCAGCAGATCGTGGTATGTGCCCAGGGAGGCGTTGAACCGATCTTCCACGCCGGAGA

TTTCCACGGAGTCGAAGCACTCGATTTTCTTGAAGTAGTCCTCTTTCAGCTGCTTCACGGTCACTTTCCGGTTGGTCTTG

AACAGCAGGTCCACGATGGCCTTTTTCTGCTCGCCGCTCAGGAAGGCGGGCTTTCTCATTCCCTCGGTCACGTATTTCAC

TTTGGTCAGCTCGTTATACACGGTGAAGTACTCGTACAGCAGGCTGTGCTTGGGCAGCACCTTCTCGTTGGGCAGGTTCT

TATCGAAGTTGGTCATCCGCTCGATGAAGCTCTGGGCGGAAGCGCCCTTGTCCACCACTTCCTCGAAGTTCCAGGGGGTG

ATGGTTTCCTCGCTCTTTCTGGTCATCCAGGCGAATCTGCTGTTTCCCCTGGCCAGAGGGCCCACGTAGTAGGGGATGCG

GAAGGTCAGGATCTTCTCGATCTTTTCCCGGTTGTCCTTCAGGAATGGGTAAAAATCTTCCTGCCGCCGCAGAATGGCGT

GCAGCTCTCCCAGGTGGATCTGGTGGGGGATGCTGCCGTTGTCGAAGGTCCGCTGCTTCCGCAGCAGGTCCTCTCTGTTC

AGCTTCACGAGCAGTTCCTCGGTGCCGTCCATCTTTTCCAGGATGGGCTTGATGAACTTGTAGAACTCTTCCTGGCTGGC

TCCGCCGTCAATGTAGCCGGCGTAGCCGTTCTTGCTCTGGTCGAAGAAAATCTCTTTGTACTTCTCAGGCAGCTGCTGCC

GCACGAGAGCTTTCAGCAGGGTCAGGTCCTGGTGGTGCTCGTCGTATCTCTTGATCATAGAGGCGCTCAGGGGGGCCTTG

GTGATCTCGGTGTTCACTCTCAGGATGTCGCTCAGCAGGATGGCGTCGGACAGGTTCTTGGCGGCCAGAAACAGGTCGGC

GTACTGGTCGCCGATCTGGGCCAGCAGGTTGTCCAGGTCGTCGTCGTAGGTGTCCTTGCTCAGCTGCAGTTTGGCATCCT

CGGCCAGGTCGAAGTTGCTCTTGAAGTTGGGGGTCAGGCCCAGGCTCAGGGCAATCAGGTTTCCGAACAGGCCATTCTTC

TTCTCGCCGGGCAGCTGGGCGATCAGATTTTCCAGCCGTCTGCTCTTGCTCAGTCTGGCAGACAGGATGGCCTTGGCGTC

CACGCCGCTGGCGTTGATGGGGTTTTCCTCGAACAGCTGGTTGTAGGTCTGCACCAGCTGGATGAACAGCTTGTCCACGT

CGCTGTTGTCGGGGTTCAGGTCGCCCTCGATCAGGAAGTGGCCCCGGAACTTGATCATGTGGGCCAGGGCCAGATAGATC

AGCCGCAGGTCGGCCTTGTCGGTGCTGTCCACCAGTTTCTTTCTCAGGTGGTAGATGGTGGGGTACTTCTCGTGGTAGGC

CACCTCGTCCACGATGTTGCCGAAGATGGGGTGCCGCTCGTGCTTCTTATCCTCTTCCACCAGGAAGGACTCTTCCAGTC

TGTGGAAGAAGCTGTCGTCCACCTTGGCCATCTCGTTGCTGAAGATCTCTTGCAGATAGCAGATCCGGTTCTTCCGTCTG

GTGTATCTTCTTCTGGCGGTTCTCTTCAGCCGGGTGGCCTCGGCTGTTTCGCCGCTGTCGAACAGCAGGGCTCCGATCAG

GTTCTTCTTGATGCTGTGCCGGTCGGTGTTGCCCAGCACCTTGAATTTCTTGCTGGGCACCTTGTACTCGTCGGTGATCA

CGGCCCAGCCCACAGAGTTGGTGCCGATGTCCAGGCCGATGCTGTACTTCTTGTCGGCTGCTGGGACTCCGTGGATACCG

ACCTTCCGCTTCTTCTTTGGGGCCATCTTATCGTCATCGTCTTTGTAATCAATATCATGATCCTTGTAGTCTCCGTCGTG

GTCCTTATAGTCCATcctaggTGATATATTTCTATTAGGTATTTATTATTATAAAATATAAATCTTGAATGATAATAAAT

AAAATATTAGTTATTCCTTTTCTAGTTTAAAATATACATATTATAAATATATATATATATATATATATTTTTATTGTGAC

AAGAATATATAATTATAAATTATATTATTTATTTTTGTATTTTTTTTTTTTTTTTTTTTTTTTTCTTTTTTTGTTTTATT

TTTCTTTTTTTTTATAAATATTATTTTTTTCTTTTATCATGCACATTGGAATAATACATTAATATATATATATATATTAT

ATTATACATATATTGAATAATGTTTATAAAAAATGCATAACTTATATGAATATAATTTTTTTTAAATATGACAAAAAGAA

AAAAAAAAAAAACCAAAAAAAATTAAAATTGAAATGAAATATATAAATATATTATTTATATATATTATACATTGTTTAAT

ACTACTACATGTATATATATATATTATATATATATATATATATCAATTTTTTCAAAAATAAATTAATATAAAAAGAGGGG

AAAAAAAAAAAAAAAAAAAAAAAAAGATAATTAAGTAAGCATTTAAAAATATATAAATTGATAATATATAAAATTAATCA

CATATAAACTAATATAATTTATAAAATAAGGAAAATAAAATATTACCATAAAATAAAAATAAAAATAAAAAAAAAAAAAA

AAAACACCTTTTTTTATATATATTAATATATAATTATCTCTTAGAAAAAATATTGTATAATTATATATGTAATGATTTAT

ATAAAAAAATAAAATTATACAAGTATATATTTTGTTTCTATAAATTGATATCTTAATTATTTATTATTAGAAATAGATAT

TTTTATAATAAACCAATAGATAAAATTTGTAGAGAAAAAAAAATAAAAATAAAAATAAAAATAATATAATATATAATAAA

ATAAAATAATATTATATAAATATATTTTAATTTTTTTTACAAAATGGTTGGGGgcgccaccggtGTAAAATAATAATTAT

AAAATATAAAAATACAATTAAAGATATAATTTAGATATATATAAATTGATGATACGTATTTTATAATAATATATGTTCAA

TAATTTTATAAATTATATAATTAATAATAATATGAAACAATAGTACGAGAAAAATATTATTATGAATATTATGTAAAGAA

TAATAAATATTTCATAAATGATTCTTTTTAATAATATGATAAGTTCATTTATTATTTTATTATAATATTATGTTATTTAT

ATCATATTTATATTATATAACTACAATAATTTGTAGATATTGTTACATAATATTATGATATTACAATATTTATACATACA

TATATATATATATATATATATATATACACTACTTAGCACTATTGAATAAAATATAGCATAAAAAAAATATACATATAATG

GCAAACCTTTGGTATAGAAAAAAATATTTAATTTATTACATTTGTTGTAGGTGAAAAATATATTTGGATGAAATAAATTG

TTCATAATAATGATTATAATATAACATTGAATACATAAATATTTTTTTATCAAAATGCATGCCAAGCCTTTGTCTCAAGA

AGAATCCACCCTCATTGAAAGAGCAACGGCTACAATCAACAGCATCCCCATCTCTGAAGACTACAGCGTCGCCAGCGCAG

CTCTCTCTAGCGACGGCCGCATCTTCACTGGTGTCAATGTATATCATTTTACTGGGGGACCTTGTGCAGAACTCGTGGTG

CTGGGCACTGCTGCTGCTGCGGCAGCTGGCAACCTGACTTGTATCGTCGCGATCGGAAATGAGAACAGGGGCATCTTGAG

CCCCTGCGGACGGTGCCGACAGGTGCTTCTCGATCTGCATCCTGGGATCAAAGCCATAGTGAAGGACAGTGATGGACAGC

CGACGGCAGTTGGGATTCGTGAATTGCTGCCCTCTGGTTATGTGTGGGAGGGCgCTAGcGGCAGTGGAGAGGGCAGAGGA

AGTCTGCTAACATGCGGTGACGTCGAGGAGAATCCTGGCCCAaagcttATGGGGTCAAGCGCATCAAAATTTTCTAAAAT

TGTTGTTGGAAATGAAACACACAAGAGTGCCAGAAATGTTTTGGAAGGTTTTGCAAAAGATATAAAAGGGAAAGCGTCAA

TAGACGCAGAAAAACATGCATATTCGTTAAAAGGAAATTTGAAAGACGCAAAATTTAATCATGATTTTTTTAAAATAAAA

AGTGACATGCCTGGAAATCCATGTTATCTTGATTTTGCTTTTCATTCTAATACTCCTGGAAATCAAAGAGAATATAGACA

TCCTTGTGCTCGTAGTATGAACAAAAATTTGTTTAATTTGGAAGGAGCTGTATGTACTAATAGTAAAATAAAGGGTAATG

AAGAAAAAATAAATGGCGCTGGAGCATGTGCCCCATATAGAAGAAGACATATATGTGACTTAAATTTGGAACACATAGAT

GTACATAATGTACAAAATATTCATGACTTATTGGGAAATGTATTAGTTACAGCAAAATATGAAGGCGAATCTATTGTTGA

GAAACATCCAAATAGAGGATCTTCAGAAGTATGTACTGCCCTTGCACGAAGTTTTGCAGATATAGGTGATATTATACGAG

GAAAAGATgtcgacaCTAGTACCGGTACGCGTgacgtCAGGTGGCACTTTTCGGGGAAATGTGCGCGGAACCCCTATTTG

TTTATTTTTCTAAATACATTCAAATATGTATCCGCTCATGAGACAATAACCCTGATAAATGCTTCAATAATATTGAAAAA

GGAAGAGTATGAGTATTCAACATTTCCGTGTCGCCCTTATTCCCTTTTTTGCGGCATTTTGCCTTCCTGTTTTTGCTCAC

CCAGAAACGCTGGTGAAAGTAAAAGATGCTGAAGATCAGTTGGGTGCACGAGTGGGTTACATCGAACTGGATCTCAACAG

CGGTAAGATCCTTGAGAGTTTTCGCCCCGAAGAACGTTTTCCAATGATGAGCACTTTTAAAGTTCTGCTATGTGGCGCGG

TATTATCCCGTATTGACGCCGGGCAAGAGCAACTCGGTCGCCGCATACACTATTCTCAGAATGACTTGGTTGAGTACTCA

CCAGTCACAGAAAAGCATCTTACGGATGGCATGACAGTAAGAGAATTATGCAGTGCTGCCATAACCATGAGTGATAACAC

TGCGGCCAACTTACTTCTGACAACGATCGGAGGACCGAAGGAGCTAACCGCTTTTTTGCACAACATGGGGGATCATGTAA

CTCGCCTTGATCGTTGGGAACCGGAGCTGAATGAAGCCATACCAAACGACGAGCGTGACACCACGATGCCTGTAGCAATG

CCAACAACGTTGCGCAAACTATTAACTGGCGAACTACTTACTCTAGCTTCCCGGCAACAATTAATAGACTGGATGGAGGC

GGATAAAGTTGCAGGACCACTTCTGCGCTCGGCCCTTCCGGCTGGCTGGTTTATTGCTGATAAATCTGGAGCCGGTGAGC

GTGGGTCTCGCGGTATCATTGCAGCACTGGGGCCAGATGGTAAGCCCTCCCGTATCGTAGTTATCTACACGACGGGGAGT

CAGGCAACTATGGATGAACGAAATAGACAGATCGCTGAGATAGGTGCCTCACTGATTAAGCATTGGTAACTGTCAGACCA

AGTTTACTCATATATACTTTAGATTGATTTAAAACTTCATTTTTAATTTAAAAGGATCTAGGTGAAGATCCTTTTTGATA

ATCTCATGACCAAAATCCCTTAACGTGAGTTTTCGTTCCACTGAGCGTCAGACCCCGTAGAAAAGATCAAAGGATCTTCT

TGAGATCCTTTTTTTCTGCGCGTAATCTGCTGCTTGCAAACAAAAAAACCACCGCTACCAGCGGTGGTTTGTTTGCCGGA

TCAAGAGCTACCAACTCTTTTTCCGAAGGTAACTGGCTTCAGCAGAGCGCAGATACCAAATACTGTCCTTCTAGTGTAGC

CGTAGTTAGGCCACCACTTCAAGAACTCTGTAGCACCGCCTACATACCTCGCTCTGCTAATCCTGTTACCAGTGGCTGCT

GCCAGTGGCGATAAGTCGTGTCTTACCGGGTTGGACTCAAGACGATAGTTACCGGATAAGGCGCAGCGGTCGGGCTGAAC

GGGGGGTTCGTGCACACAGCCCAGCTTGGAGCGAACGACCTACACCGAACTGAGATACCTACAGCGTGAGCTATGAGAAA

GCGCCACGCTTCCCGAAGGGAGAAAGGCGGACAGGTATCCGGTAAGCGGCAGGGTCGGAACAGGAGAGCGCACGAGGGAG

CTTCCAGGGGGAAACGCCTGGTATCTTTATAGTCCTGTCGGGTTTCGCCACCTCTGACTTGAGCGTCGATTTTTGTGATG

CTCGTCAGGGGGGCGGAGCCTATCGAAAAACGCCAGCAACGCGGCCTTTTTACGGTTCCTGGCCTTTTGCTGGCCTTTTG

CTCACATGTTCTTTCCTGCGTTATCCCCTGATTCTGTGGATAACCGTATTACCGCCTTTGAGTGAGCTGATACCGCTCGC

CGCAGCCGAACGACCGAGCGCAGCGAGTCAGTGAGCGAGGAAGCGGAAGAGCGCCCAATACGCAAACCGCCTCTCCCCGC

GCGTTGGCCGATTCATTAATGCAGCTGGCACGACAGGTTTCCCGACTGGAAAGCGGGCAGTGAGCGCAACGCAATTAATG

TGAGTTAGCTCACTCATTAGGCACCCCAGGCTTTACACTTTATGCTTCCGGCTCGTATGTTGTGTGGAATTGTGAGCGGA

TAACAATTTCACACAGGAAACAGCTATGACCATGATTACGCCAAGCTATTTAGGTGACACTATAGAATACTC

Nucleotide sequence of plasmid Cas9_itvar60prom_bsd_2A_HA_exonI

AAGCTTGGGGGGATCCGCCTTAAAAACTTCATTATATTTAAAAATTATTTTATAGGAAATAATAAAAAAAAAAgcaccga

ctcggtgccactttttcaagttgataacggactagccttattttaacttgctaTTTCtagctctaaaacTTTGGTGCCAT

TCTTATAACAATATTATATACTTAATATGAAATATGTGCATATAGGAAAAATTATGCATTTTGGTTACTCTAATATTATA

TATATATATATATATATATATATATTATAATATATTATGTTATATATACATAACATATACATTTTTTAATAATAATTTAC

CCTTTATTTTTACATTATAAAAAATTATATTACAGTAAAAATAAAAGTTTATTATATTAATAGTTTTTTTTTTTTTTTTT

TTAATTTATGAAATATTTAAATATTTAAAATTTTTTAAATGAATAATTATATTTATAATTAGAAAAAAAAAAAAAAAAAA

AAAAAAAAAATATAGCTATTTATATAAATTTCTTTTATTTATCTGAACAAGCAAGAATTTTTTTTTATATTAAATTAGAA

TAAATTATTATTAGTTTATGTATATATTTTTTTTTTTTCATAGTATATAAATATTATATATATTGTACCTTTTTACAATA

TATTTCATATATAGAAGAGAAAAAAAAAAAAAGAAGATATTATTGTAAAACCTCAAGATGTGTAGAAATCCAAATGTCGG

ATCCTCTAGAGTCGACCTGCAGGCATGCTATTTGATGAATTAACTACACTTAAAATAATACAATTATTATTAAATTTTTT

TTTGATTTATTTATTAATTTTTAAACTTAATCATTTGTATTTGGGAGGAATTATATATATCTTTATAATTATTTTATTTT

TTTTTATTTTTTTATTTTTTTATTATTATTATTTTTTTTTATTTTTTTTTTTTACTGTATCAAAGAAAAACCTTTAAAAA

AAAAATTATAATTTCCCCATCTTACTATATTTTTAATACATACGTTTTAAGGAATTAAATTAGACAAAAGCTATATTATG

CTTTACATATAATTAGAATTTATAAACGTTTGGTTATTAGATATTTCATGTCTCAGTAAAGTCTTTCAATACATATGTAA

AAAAATATATATGAATACACATAAGTTGTTAATATATTTTATATGCATAAATGTATAAATATATATATATATATATATAT

ATGTATGTATGTATATGTGTGTATATGAAATTATTTCAATGTTTAATTTTTTAAATTTTAATTTTTTTTTTTTTTTTTTT

TTTTATTATGTATATTGATCTTTATTATTTAAATATTACTTTTTTCGTTTTTTCTTCTTTTTATTATTTTTTTTTTTTTT

TATATTTTATACAAATGGTAATTCAAATAAAAGGTATAAATTTATATTTAATTTTCTTTTATGGATAAATAAAAGAAAAA

TATAAATATATAAAAATATAAAAATATATATATGTATATTGGGGTGATGATAAAATGAAAGATAATATATATATATATAT

ATCTTTATTTTTTTTTTTTTGTAGACCCCATTGTGAGTACATAAATATATTATATAACTCGAGTTACTTTTTCTTTTTTG

CCTGGCCGGCCTTTTTCGTGGCCGCCGGCCTTTTGTCGCCTCCCAGCTGAGACAGGTCGATCCGTGTCTCGTACAGGCCG

GTGATGCTCTGGTGGATCAGGGTGGCGTCCAGCACCTCTTTGGTGCTGGTGTACCTCTTCCGGTCGATGGTGGTGTCAAA

GTACTTGAAGGCGGCAGGGGCTCCCAGATTGGTCAGGGTAAACAGGTGGATGATATTCTCGGCCTGCTCTCTGATGGGCT

TATCCCGGTGCTTGTTGTAGGCGGACAGCACTTTGTCCAGATTAGCGTCGGCCAGGATCACTCTCTTGGAGAACTCGCTG

ATCTGCTCGATGATCTCGTCCAGGTAGTGCTTGTGCTGTTCCACAAACAGCTGTTTCTGCTCATTATCCTCGGGGGAGCC

CTTCAGCTTCTCATAGTGGCTGGCCAGGTACAGGAAGTTCACATATTTGGAGGGCAGGGCCAGTTCGTTTCCCTTCTGCA

GTTCGCCGGCAGAGGCCAGCATTCTCTTCCGGCCGTTTTCCAGCTCGAACAGGGAGTACTTAGGCAGCTTGATGATCAGG

TCCTTTTTCACTTCTTTGTAGCCCTTGGCTTCCAGAAAGTCGATGGGATTCTTCTCGAAGCTGCTTCTTTCCATGATGGT

GATCCCCAGCAGCTCTTTCACACTCTTCAGTTTCTTGGACTTGCCCTTTTCCACTTTGGCCACCACCAGCACAGAATAGG

CCACGGTGGGGCTGTCGAAGCCGCCGTACTTCTTAGGGTCCCAGTCCTTCTTTCTGGCGATCAGCTTATCGCTGTTCCTC

TTGGGCAGGATAGACTCTTTGCTGAAGCCGCCTGTCTGCACCTCGGTCTTTTTCACGATATTCACTTGGGGCATGCTCAG

CACTTTCCGCACGGTGGCAAAATCCCGGCCCTTATCCCACACGATCTCCCCGGTTTCGCCGTTTGTCTCGATCAGAGGCC

GCTTCCGGATCTCGCCGTTGGCCAGGGTAATCTCGGTCTTGAAAAAGTTCATGATGTTGCTGTAGAAGAAGTACTTGGCG

GTAGCCTTGCCGATTTCCTGCTCGCTCTTGGCGATCATCTTCCGCACGTCGTACACCTTGTAGTCGCCGTACACGAACTC

GCTTTCCAGCTTAGGGTACTTTTTGATCAGGGCGGTTCCCACGACGGCGTTCAGGTAGGCGTCGTGGGCGTGGTGGTAGT

TGTTGATCTCGCGCACTTTGTAAAACTGGAAATCCTTCCGGAAATCGGACACCAGCTTGGACTTCAGGGTGATCACTTTC

ACTTCCCGGATCAGCTTGTCATTCTCGTCGTACTTAGTGTTCATCCGGGAGTCCAGGATCTGTGCCACGTGCTTTGTGAT

CTGCCGGGTTTCCACCAGCTGTCTCTTGATGAAGCCGGCCTTATCCAGTTCGCTCAGGCCGCCTCTCTCGGCCTTGGTCA

GATTGTCGAACTTTCTCTGGGTAATCAGCTTGGCGTTCAGCAGCTGCCGCCAGTAGTTCTTCATCTTCTTCACGACCTCT

TCGGAGGGCACGTTGTCGCTCTTGCCCCGGTTCTTGTCGCTTCTGGTCAGCACCTTGTTGTCGATGGAGTCGTCCTTCAG

AAAGCTCTGAGGCACGATATGGTCCACATCGTAGTCGGACAGCCGGTTGATGTCCAGTTCCTGGTCCACGTACATATCCC

GCCCATTCTGCAGGTAGTACAGGTACAGCTTCTCGTTCTGCAGCTGGGTGTTTTCCACGGGGTGTTCTTTCAGGATCTGG

CTGCCCAGCTCTTTGATGCCCTCTTCGATCCGCTTCATTCTCTCGCGGCTGTTCTTCTGTCCCTTCTGGGTGGTCTGGTT

CTCTCTGGCCATTTCGATCACGATGTTCTCGGGCTTGTGCCGGCCCATCACTTTCACGAGCTCGTCCACCACCTTCACTG

TCTGCAGGATGCCCTTCTTAATGGCGGGGCTGCCGGCCAGATTGGCAATGTGCTCGTGCAGGCTATCGCCCTGGCCGGAC

ACCTGGGCTTTCTGGATGTCCTCTTTAAAGGTCAGGCTGTCGTCGTGGATCAGCTGCATGAAGTTTCTGTTGGCGAAGCC

GTCGGACTTCAGGAAATCCAGGATTGTCTTGCCGGACTGCTTGTCCCGGATGCCGTTGATCAGCTTCCGGCTCAGCCTGC

CCCAGCCGGTGTATCTCCGCCGCTTCAGCTGCTTCATCACTTTGTCGTCGAACAGGTGGGCATAGGTTTTCAGCCGTTCC

TCGATCATCTCTCTGTCCTCAAACAGTGTCAGGGTCAGCACGATATCTTCCAGAATGTCCTCGTTTTCCTCATTGTCCAG

GAAGTCCTTGTCCTTGATAATTTTCAGCAGATCGTGGTATGTGCCCAGGGAGGCGTTGAACCGATCTTCCACGCCGGAGA

TTTCCACGGAGTCGAAGCACTCGATTTTCTTGAAGTAGTCCTCTTTCAGCTGCTTCACGGTCACTTTCCGGTTGGTCTTG

AACAGCAGGTCCACGATGGCCTTTTTCTGCTCGCCGCTCAGGAAGGCGGGCTTTCTCATTCCCTCGGTCACGTATTTCAC

TTTGGTCAGCTCGTTATACACGGTGAAGTACTCGTACAGCAGGCTGTGCTTGGGCAGCACCTTCTCGTTGGGCAGGTTCT

TATCGAAGTTGGTCATCCGCTCGATGAAGCTCTGGGCGGAAGCGCCCTTGTCCACCACTTCCTCGAAGTTCCAGGGGGTG

ATGGTTTCCTCGCTCTTTCTGGTCATCCAGGCGAATCTGCTGTTTCCCCTGGCCAGAGGGCCCACGTAGTAGGGGATGCG

GAAGGTCAGGATCTTCTCGATCTTTTCCCGGTTGTCCTTCAGGAATGGGTAAAAATCTTCCTGCCGCCGCAGAATGGCGT

GCAGCTCTCCCAGGTGGATCTGGTGGGGGATGCTGCCGTTGTCGAAGGTCCGCTGCTTCCGCAGCAGGTCCTCTCTGTTC

AGCTTCACGAGCAGTTCCTCGGTGCCGTCCATCTTTTCCAGGATGGGCTTGATGAACTTGTAGAACTCTTCCTGGCTGGC

TCCGCCGTCAATGTAGCCGGCGTAGCCGTTCTTGCTCTGGTCGAAGAAAATCTCTTTGTACTTCTCAGGCAGCTGCTGCC

GCACGAGAGCTTTCAGCAGGGTCAGGTCCTGGTGGTGCTCGTCGTATCTCTTGATCATAGAGGCGCTCAGGGGGGCCTTG

GTGATCTCGGTGTTCACTCTCAGGATGTCGCTCAGCAGGATGGCGTCGGACAGGTTCTTGGCGGCCAGAAACAGGTCGGC

GTACTGGTCGCCGATCTGGGCCAGCAGGTTGTCCAGGTCGTCGTCGTAGGTGTCCTTGCTCAGCTGCAGTTTGGCATCCT

CGGCCAGGTCGAAGTTGCTCTTGAAGTTGGGGGTCAGGCCCAGGCTCAGGGCAATCAGGTTTCCGAACAGGCCATTCTTC

TTCTCGCCGGGCAGCTGGGCGATCAGATTTTCCAGCCGTCTGCTCTTGCTCAGTCTGGCAGACAGGATGGCCTTGGCGTC

CACGCCGCTGGCGTTGATGGGGTTTTCCTCGAACAGCTGGTTGTAGGTCTGCACCAGCTGGATGAACAGCTTGTCCACGT

CGCTGTTGTCGGGGTTCAGGTCGCCCTCGATCAGGAAGTGGCCCCGGAACTTGATCATGTGGGCCAGGGCCAGATAGATC

AGCCGCAGGTCGGCCTTGTCGGTGCTGTCCACCAGTTTCTTTCTCAGGTGGTAGATGGTGGGGTACTTCTCGTGGTAGGC

CACCTCGTCCACGATGTTGCCGAAGATGGGGTGCCGCTCGTGCTTCTTATCCTCTTCCACCAGGAAGGACTCTTCCAGTC

TGTGGAAGAAGCTGTCGTCCACCTTGGCCATCTCGTTGCTGAAGATCTCTTGCAGATAGCAGATCCGGTTCTTCCGTCTG

GTGTATCTTCTTCTGGCGGTTCTCTTCAGCCGGGTGGCCTCGGCTGTTTCGCCGCTGTCGAACAGCAGGGCTCCGATCAG

GTTCTTCTTGATGCTGTGCCGGTCGGTGTTGCCCAGCACCTTGAATTTCTTGCTGGGCACCTTGTACTCGTCGGTGATCA

CGGCCCAGCCCACAGAGTTGGTGCCGATGTCCAGGCCGATGCTGTACTTCTTGTCGGCTGCTGGGACTCCGTGGATACCG

ACCTTCCGCTTCTTCTTTGGGGCCATCTTATCGTCATCGTCTTTGTAATCAATATCATGATCCTTGTAGTCTCCGTCGTG

GTCCTTATAGTCCATcctaggTGATATATTTCTATTAGGTATTTATTATTATAAAATATAAATCTTGAATGATAATAAAT

AAAATATTAGTTATTCCTTTTCTAGTTTAAAATATACATATTATAAATATATATATATATATATATATTTTTATTGTGAC

AAGAATATATAATTATAAATTATATTATTTATTTTTGTATTTTTTTTTTTTTTTTTTTTTTTTTCTTTTTTTGTTTTATT

TTTCTTTTTTTTTATAAATATTATTTTTTTCTTTTATCATGCACATTGGAATAATACATTAATATATATATATATATTAT

ATTATACATATATTGAATAATGTTTATAAAAAATGCATAACTTATATGAATATAATTTTTTTTAAATATGACAAAAAGAA

AAAAAAAAAAAACCAAAAAAAATTAAAATTGAAATGAAATATATAAATATATTATTTATATATATTATACATTGTTTAAT

ACTACTACATGTATATATATATATTATATATATATATATATATCAATTTTTTCAAAAATAAATTAATATAAAAAGAGGGG

AAAAAAAAAAAAAAAAAAAAAAAAAGATAATTAAGTAAGCATTTAAAAATATATAAATTGATAATATATAAAATTAATCA

CATATAAACTAATATAATTTATAAAATAAGGAAAATAAAATATTACCATAAAATAAAAATAAAAATAAAAAAAAAAAAAA

AAAACACCTTTTTTTATATATATTAATATATAATTATCTCTTAGAAAAAATATTGTATAATTATATATGTAATGATTTAT

ATAAAAAAATAAAATTATACAAGTATATATTTTGTTTCTATAAATTGATATCTTAATTATTTATTATTAGAAATAGATAT

TTTTATAATAAACCAATAGATAAAATTTGTAGAGAAAAAAAAATAAAAATAAAAATAAAAATAATATAATATATAATAAA

ATAAAATAATATTATATAAATATATTTTAATTTTTTTTACAAAATGGTTGGGGgcgccaccggtTATGTAATAATAATAT

GATCATAATATTATAATAAAACTTATAAAAAAAATATTAAATATTTCATAAATGATTATTATTTATAATAAGATAGATTC

TTTAATTATTTTAAAATTGTATATTTTTTATGTATATTAATTTATTAATATTAATAAGAATATTATAAAAATTCTATTTT

ATTATTTAATGATATTACCCTAATAAAAATAATATAATTATATTACAATATAAATTTATATATATATATATATTTATATA

TTTAAAGAATATTTTATTTTTCAATAAGAACCTTCATTTTAAATTAACATCAAATTATATATATGTATATATACTTCTTA

GTATTATTAATTAAAATACGGAATAATATATAATATATATAAAATGGCAAAACTTTCCTATAGAAAAAAATATTCCATTT

ATTATATTTGTTGTAGGTAATTCTTATTACCGTTTCCTTCTGTTCGTAATGTATATTGGTATGTACTTTATTTTTGCAAT

TTAATTATATGTAAAAAAACGTTAGTACACCATATATATATTATAGTTATAAGAATGCATGCCAAGCCTTTGTCTCAAGA

AGAATCCACCCTCATTGAAAGAGCAACGGCTACAATCAACAGCATCCCCATCTCTGAAGACTACAGCGTCGCCAGCGCAG

CTCTCTCTAGCGACGGCCGCATCTTCACTGGTGTCAATGTATATCATTTTACTGGGGGACCTTGTGCAGAACTCGTGGTG

CTGGGCACTGCTGCTGCTGCGGCAGCTGGCAACCTGACTTGTATCGTCGCGATCGGAAATGAGAACAGGGGCATCTTGAG

CCCCTGCGGACGGTGCCGACAGGTGCTTCTCGATCTGCATCCTGGGATCAAAGCCATAGTGAAGGACAGTGATGGACAGC

CGACGGCAGTTGGGATTCGTGAATTGCTGCCCTCTGGTTATGTGTGGGAGGGCgCTAGcGGCAGTGGAGAGGGCAGAGGA

AGTCTGCTAACATGCGGTGACGTCGAGGAGAATCCTGGCCCAaagcttTACCCATACGATGTTCCAGATTACGCTgccAT

GGCACCAAAGGGTAGAAGTACAAATGAAATTGAACTTAGCGCAAGAGATGTTTTGGAAAATATTGGAATAGGAATATATA

ATCAGGAAAAAATAAAAAAGAATCCATATGAACAACAATTGAAAGGCACATTATCAAACGCCCGATTTCATGATGGCTTG

CACAAGGCAGCTGATTTGGGGGTAATACCTGGTCCTTCACATTTTTCTCAGCTTTATTACAAAAAGCATACTAATAACAC

AAAATATTATAAGGATGATAGGCATCCTTGTCATGGTAGACAAGGAAAACGTTTTGATGAAGGTCAAAAATTTGAATGTG

GTAATGATAAAATAATTGGTAATAGCGATAAATATGGATCCTGTGCTCCACCTAGAAGAAGACATATATGTGATCAAAAT

TTAGAATTCTTAGATAACAATCATACTGATACTATTCATGATGTATTGGGAAATGTGTTGGTCACAGCAAAATATGAAGG

TGAATCTATTGTTAATGATCATCCAGATAAAAAGAACAATGGTAATAAATCAGGTATATGTACTTCTCTTGCACGAAGTT

TTGCCGATATAGGTGATATTGTAAGAGGAAGAGATATGgtcgacaCTAGTACCGGTACGCGTgacgtCAGGTGGCACTTT

TCGGGGAAATGTGCGCGGAACCCCTATTTGTTTATTTTTCTAAATACATTCAAATATGTATCCGCTCATGAGACAATAAC

CCTGATAAATGCTTCAATAATATTGAAAAAGGAAGAGTATGAGTATTCAACATTTCCGTGTCGCCCTTATTCCCTTTTTT

GCGGCATTTTGCCTTCCTGTTTTTGCTCACCCAGAAACGCTGGTGAAAGTAAAAGATGCTGAAGATCAGTTGGGTGCACG

AGTGGGTTACATCGAACTGGATCTCAACAGCGGTAAGATCCTTGAGAGTTTTCGCCCCGAAGAACGTTTTCCAATGATGA

GCACTTTTAAAGTTCTGCTATGTGGCGCGGTATTATCCCGTATTGACGCCGGGCAAGAGCAACTCGGTCGCCGCATACAC

TATTCTCAGAATGACTTGGTTGAGTACTCACCAGTCACAGAAAAGCATCTTACGGATGGCATGACAGTAAGAGAATTATG

CAGTGCTGCCATAACCATGAGTGATAACACTGCGGCCAACTTACTTCTGACAACGATCGGAGGACCGAAGGAGCTAACCG

CTTTTTTGCACAACATGGGGGATCATGTAACTCGCCTTGATCGTTGGGAACCGGAGCTGAATGAAGCCATACCAAACGAC

GAGCGTGACACCACGATGCCTGTAGCAATGCCAACAACGTTGCGCAAACTATTAACTGGCGAACTACTTACTCTAGCTTC

CCGGCAACAATTAATAGACTGGATGGAGGCGGATAAAGTTGCAGGACCACTTCTGCGCTCGGCCCTTCCGGCTGGCTGGT

TTATTGCTGATAAATCTGGAGCCGGTGAGCGTGGGTCTCGCGGTATCATTGCAGCACTGGGGCCAGATGGTAAGCCCTCC

CGTATCGTAGTTATCTACACGACGGGGAGTCAGGCAACTATGGATGAACGAAATAGACAGATCGCTGAGATAGGTGCCTC

ACTGATTAAGCATTGGTAACTGTCAGACCAAGTTTACTCATATATACTTTAGATTGATTTAAAACTTCATTTTTAATTTA

AAAGGATCTAGGTGAAGATCCTTTTTGATAATCTCATGACCAAAATCCCTTAACGTGAGTTTTCGTTCCACTGAGCGTCA

GACCCCGTAGAAAAGATCAAAGGATCTTCTTGAGATCCTTTTTTTCTGCGCGTAATCTGCTGCTTGCAAACAAAAAAACC

ACCGCTACCAGCGGTGGTTTGTTTGCCGGATCAAGAGCTACCAACTCTTTTTCCGAAGGTAACTGGCTTCAGCAGAGCGC

AGATACCAAATACTGTCCTTCTAGTGTAGCCGTAGTTAGGCCACCACTTCAAGAACTCTGTAGCACCGCCTACATACCTC

GCTCTGCTAATCCTGTTACCAGTGGCTGCTGCCAGTGGCGATAAGTCGTGTCTTACCGGGTTGGACTCAAGACGATAGTT

ACCGGATAAGGCGCAGCGGTCGGGCTGAACGGGGGGTTCGTGCACACAGCCCAGCTTGGAGCGAACGACCTACACCGAAC

TGAGATACCTACAGCGTGAGCTATGAGAAAGCGCCACGCTTCCCGAAGGGAGAAAGGCGGACAGGTATCCGGTAAGCGGC

AGGGTCGGAACAGGAGAGCGCACGAGGGAGCTTCCAGGGGGAAACGCCTGGTATCTTTATAGTCCTGTCGGGTTTCGCCA

CCTCTGACTTGAGCGTCGATTTTTGTGATGCTCGTCAGGGGGGCGGAGCCTATCGAAAAACGCCAGCAACGCGGCCTTTT

TACGGTTCCTGGCCTTTTGCTGGCCTTTTGCTCACATGTTCTTTCCTGCGTTATCCCCTGATTCTGTGGATAACCGTATT

ACCGCCTTTGAGTGAGCTGATACCGCTCGCCGCAGCCGAACGACCGAGCGCAGCGAGTCAGTGAGCGAGGAAGCGGAAGA

GCGCCCAATACGCAAACCGCCTCTCCCCGCGCGTTGGCCGATTCATTAATGCAGCTGGCACGACAGGTTTCCCGACTGGA

AAGCGGGCAGTGAGCGCAACGCAATTAATGTGAGTTAGCTCACTCATTAGGCACCCCAGGCTTTACACTTTATGCTTCCG

GCTCGTATGTTGTGTGGAATTGTGAGCGGATAACAATTTCACACAGGAAACAGCTATGACCATGATTACGCCAAGCTATT

TAGGTGACACTATAGAATACTC

Nucleotide sequence of plasmid Cas9_itvar60prom_bsd_2A_myc_exonI

AAGCTTGGGGGGATCCGCCTTAAAAACTTCATTATATTTAAAAATTATTTTATAGGAAATAATAAAAAAAAAAgcaccga

ctcggtgccactttttcaagttgataacggactagccttattttaacttgctaTTTCtagctctaaaacTTTGGTGCCAT

TCTTATAACAATATTATATACTTAATATGAAATATGTGCATATAGGAAAAATTATGCATTTTGGTTACTCTAATATTATA

TATATATATATATATATATATATATTATAATATATTATGTTATATATACATAACATATACATTTTTTAATAATAATTTAC

CCTTTATTTTTACATTATAAAAAATTATATTACAGTAAAAATAAAAGTTTATTATATTAATAGTTTTTTTTTTTTTTTTT

TTAATTTATGAAATATTTAAATATTTAAAATTTTTTAAATGAATAATTATATTTATAATTAGAAAAAAAAAAAAAAAAAA

AAAAAAAAAATATAGCTATTTATATAAATTTCTTTTATTTATCTGAACAAGCAAGAATTTTTTTTTATATTAAATTAGAA

TAAATTATTATTAGTTTATGTATATATTTTTTTTTTTTCATAGTATATAAATATTATATATATTGTACCTTTTTACAATA

TATTTCATATATAGAAGAGAAAAAAAAAAAAAGAAGATATTATTGTAAAACCTCAAGATGTGTAGAAATCCAAATGTCGG

ATCCTCTAGAGTCGACCTGCAGGCATGCTATTTGATGAATTAACTACACTTAAAATAATACAATTATTATTAAATTTTTT

TTTGATTTATTTATTAATTTTTAAACTTAATCATTTGTATTTGGGAGGAATTATATATATCTTTATAATTATTTTATTTT

TTTTTATTTTTTTATTTTTTTATTATTATTATTTTTTTTTATTTTTTTTTTTTACTGTATCAAAGAAAAACCTTTAAAAA

AAAAATTATAATTTCCCCATCTTACTATATTTTTAATACATACGTTTTAAGGAATTAAATTAGACAAAAGCTATATTATG

CTTTACATATAATTAGAATTTATAAACGTTTGGTTATTAGATATTTCATGTCTCAGTAAAGTCTTTCAATACATATGTAA

AAAAATATATATGAATACACATAAGTTGTTAATATATTTTATATGCATAAATGTATAAATATATATATATATATATATAT

ATGTATGTATGTATATGTGTGTATATGAAATTATTTCAATGTTTAATTTTTTAAATTTTAATTTTTTTTTTTTTTTTTTT

TTTTATTATGTATATTGATCTTTATTATTTAAATATTACTTTTTTCGTTTTTTCTTCTTTTTATTATTTTTTTTTTTTTT

TATATTTTATACAAATGGTAATTCAAATAAAAGGTATAAATTTATATTTAATTTTCTTTTATGGATAAATAAAAGAAAAA

TATAAATATATAAAAATATAAAAATATATATATGTATATTGGGGTGATGATAAAATGAAAGATAATATATATATATATAT

ATCTTTATTTTTTTTTTTTTGTAGACCCCATTGTGAGTACATAAATATATTATATAACTCGAGTTACTTTTTCTTTTTTG

CCTGGCCGGCCTTTTTCGTGGCCGCCGGCCTTTTGTCGCCTCCCAGCTGAGACAGGTCGATCCGTGTCTCGTACAGGCCG

GTGATGCTCTGGTGGATCAGGGTGGCGTCCAGCACCTCTTTGGTGCTGGTGTACCTCTTCCGGTCGATGGTGGTGTCAAA

GTACTTGAAGGCGGCAGGGGCTCCCAGATTGGTCAGGGTAAACAGGTGGATGATATTCTCGGCCTGCTCTCTGATGGGCT

TATCCCGGTGCTTGTTGTAGGCGGACAGCACTTTGTCCAGATTAGCGTCGGCCAGGATCACTCTCTTGGAGAACTCGCTG

ATCTGCTCGATGATCTCGTCCAGGTAGTGCTTGTGCTGTTCCACAAACAGCTGTTTCTGCTCATTATCCTCGGGGGAGCC

CTTCAGCTTCTCATAGTGGCTGGCCAGGTACAGGAAGTTCACATATTTGGAGGGCAGGGCCAGTTCGTTTCCCTTCTGCA

GTTCGCCGGCAGAGGCCAGCATTCTCTTCCGGCCGTTTTCCAGCTCGAACAGGGAGTACTTAGGCAGCTTGATGATCAGG

TCCTTTTTCACTTCTTTGTAGCCCTTGGCTTCCAGAAAGTCGATGGGATTCTTCTCGAAGCTGCTTCTTTCCATGATGGT

GATCCCCAGCAGCTCTTTCACACTCTTCAGTTTCTTGGACTTGCCCTTTTCCACTTTGGCCACCACCAGCACAGAATAGG

CCACGGTGGGGCTGTCGAAGCCGCCGTACTTCTTAGGGTCCCAGTCCTTCTTTCTGGCGATCAGCTTATCGCTGTTCCTC

TTGGGCAGGATAGACTCTTTGCTGAAGCCGCCTGTCTGCACCTCGGTCTTTTTCACGATATTCACTTGGGGCATGCTCAG

CACTTTCCGCACGGTGGCAAAATCCCGGCCCTTATCCCACACGATCTCCCCGGTTTCGCCGTTTGTCTCGATCAGAGGCC

GCTTCCGGATCTCGCCGTTGGCCAGGGTAATCTCGGTCTTGAAAAAGTTCATGATGTTGCTGTAGAAGAAGTACTTGGCG

GTAGCCTTGCCGATTTCCTGCTCGCTCTTGGCGATCATCTTCCGCACGTCGTACACCTTGTAGTCGCCGTACACGAACTC

GCTTTCCAGCTTAGGGTACTTTTTGATCAGGGCGGTTCCCACGACGGCGTTCAGGTAGGCGTCGTGGGCGTGGTGGTAGT

TGTTGATCTCGCGCACTTTGTAAAACTGGAAATCCTTCCGGAAATCGGACACCAGCTTGGACTTCAGGGTGATCACTTTC

ACTTCCCGGATCAGCTTGTCATTCTCGTCGTACTTAGTGTTCATCCGGGAGTCCAGGATCTGTGCCACGTGCTTTGTGAT

CTGCCGGGTTTCCACCAGCTGTCTCTTGATGAAGCCGGCCTTATCCAGTTCGCTCAGGCCGCCTCTCTCGGCCTTGGTCA

GATTGTCGAACTTTCTCTGGGTAATCAGCTTGGCGTTCAGCAGCTGCCGCCAGTAGTTCTTCATCTTCTTCACGACCTCT

TCGGAGGGCACGTTGTCGCTCTTGCCCCGGTTCTTGTCGCTTCTGGTCAGCACCTTGTTGTCGATGGAGTCGTCCTTCAG

AAAGCTCTGAGGCACGATATGGTCCACATCGTAGTCGGACAGCCGGTTGATGTCCAGTTCCTGGTCCACGTACATATCCC

GCCCATTCTGCAGGTAGTACAGGTACAGCTTCTCGTTCTGCAGCTGGGTGTTTTCCACGGGGTGTTCTTTCAGGATCTGG

CTGCCCAGCTCTTTGATGCCCTCTTCGATCCGCTTCATTCTCTCGCGGCTGTTCTTCTGTCCCTTCTGGGTGGTCTGGTT

CTCTCTGGCCATTTCGATCACGATGTTCTCGGGCTTGTGCCGGCCCATCACTTTCACGAGCTCGTCCACCACCTTCACTG

TCTGCAGGATGCCCTTCTTAATGGCGGGGCTGCCGGCCAGATTGGCAATGTGCTCGTGCAGGCTATCGCCCTGGCCGGAC

ACCTGGGCTTTCTGGATGTCCTCTTTAAAGGTCAGGCTGTCGTCGTGGATCAGCTGCATGAAGTTTCTGTTGGCGAAGCC

GTCGGACTTCAGGAAATCCAGGATTGTCTTGCCGGACTGCTTGTCCCGGATGCCGTTGATCAGCTTCCGGCTCAGCCTGC

CCCAGCCGGTGTATCTCCGCCGCTTCAGCTGCTTCATCACTTTGTCGTCGAACAGGTGGGCATAGGTTTTCAGCCGTTCC

TCGATCATCTCTCTGTCCTCAAACAGTGTCAGGGTCAGCACGATATCTTCCAGAATGTCCTCGTTTTCCTCATTGTCCAG

GAAGTCCTTGTCCTTGATAATTTTCAGCAGATCGTGGTATGTGCCCAGGGAGGCGTTGAACCGATCTTCCACGCCGGAGA

TTTCCACGGAGTCGAAGCACTCGATTTTCTTGAAGTAGTCCTCTTTCAGCTGCTTCACGGTCACTTTCCGGTTGGTCTTG

AACAGCAGGTCCACGATGGCCTTTTTCTGCTCGCCGCTCAGGAAGGCGGGCTTTCTCATTCCCTCGGTCACGTATTTCAC

TTTGGTCAGCTCGTTATACACGGTGAAGTACTCGTACAGCAGGCTGTGCTTGGGCAGCACCTTCTCGTTGGGCAGGTTCT

TATCGAAGTTGGTCATCCGCTCGATGAAGCTCTGGGCGGAAGCGCCCTTGTCCACCACTTCCTCGAAGTTCCAGGGGGTG

ATGGTTTCCTCGCTCTTTCTGGTCATCCAGGCGAATCTGCTGTTTCCCCTGGCCAGAGGGCCCACGTAGTAGGGGATGCG

GAAGGTCAGGATCTTCTCGATCTTTTCCCGGTTGTCCTTCAGGAATGGGTAAAAATCTTCCTGCCGCCGCAGAATGGCGT

GCAGCTCTCCCAGGTGGATCTGGTGGGGGATGCTGCCGTTGTCGAAGGTCCGCTGCTTCCGCAGCAGGTCCTCTCTGTTC

AGCTTCACGAGCAGTTCCTCGGTGCCGTCCATCTTTTCCAGGATGGGCTTGATGAACTTGTAGAACTCTTCCTGGCTGGC

TCCGCCGTCAATGTAGCCGGCGTAGCCGTTCTTGCTCTGGTCGAAGAAAATCTCTTTGTACTTCTCAGGCAGCTGCTGCC

GCACGAGAGCTTTCAGCAGGGTCAGGTCCTGGTGGTGCTCGTCGTATCTCTTGATCATAGAGGCGCTCAGGGGGGCCTTG

GTGATCTCGGTGTTCACTCTCAGGATGTCGCTCAGCAGGATGGCGTCGGACAGGTTCTTGGCGGCCAGAAACAGGTCGGC

GTACTGGTCGCCGATCTGGGCCAGCAGGTTGTCCAGGTCGTCGTCGTAGGTGTCCTTGCTCAGCTGCAGTTTGGCATCCT

CGGCCAGGTCGAAGTTGCTCTTGAAGTTGGGGGTCAGGCCCAGGCTCAGGGCAATCAGGTTTCCGAACAGGCCATTCTTC

TTCTCGCCGGGCAGCTGGGCGATCAGATTTTCCAGCCGTCTGCTCTTGCTCAGTCTGGCAGACAGGATGGCCTTGGCGTC

CACGCCGCTGGCGTTGATGGGGTTTTCCTCGAACAGCTGGTTGTAGGTCTGCACCAGCTGGATGAACAGCTTGTCCACGT

CGCTGTTGTCGGGGTTCAGGTCGCCCTCGATCAGGAAGTGGCCCCGGAACTTGATCATGTGGGCCAGGGCCAGATAGATC

AGCCGCAGGTCGGCCTTGTCGGTGCTGTCCACCAGTTTCTTTCTCAGGTGGTAGATGGTGGGGTACTTCTCGTGGTAGGC

CACCTCGTCCACGATGTTGCCGAAGATGGGGTGCCGCTCGTGCTTCTTATCCTCTTCCACCAGGAAGGACTCTTCCAGTC

TGTGGAAGAAGCTGTCGTCCACCTTGGCCATCTCGTTGCTGAAGATCTCTTGCAGATAGCAGATCCGGTTCTTCCGTCTG

GTGTATCTTCTTCTGGCGGTTCTCTTCAGCCGGGTGGCCTCGGCTGTTTCGCCGCTGTCGAACAGCAGGGCTCCGATCAG

GTTCTTCTTGATGCTGTGCCGGTCGGTGTTGCCCAGCACCTTGAATTTCTTGCTGGGCACCTTGTACTCGTCGGTGATCA

CGGCCCAGCCCACAGAGTTGGTGCCGATGTCCAGGCCGATGCTGTACTTCTTGTCGGCTGCTGGGACTCCGTGGATACCG

ACCTTCCGCTTCTTCTTTGGGGCCATCTTATCGTCATCGTCTTTGTAATCAATATCATGATCCTTGTAGTCTCCGTCGTG

GTCCTTATAGTCCATcctaggTGATATATTTCTATTAGGTATTTATTATTATAAAATATAAATCTTGAATGATAATAAAT

AAAATATTAGTTATTCCTTTTCTAGTTTAAAATATACATATTATAAATATATATATATATATATATATTTTTATTGTGAC

AAGAATATATAATTATAAATTATATTATTTATTTTTGTATTTTTTTTTTTTTTTTTTTTTTTTTCTTTTTTTGTTTTATT

TTTCTTTTTTTTTATAAATATTATTTTTTTCTTTTATCATGCACATTGGAATAATACATTAATATATATATATATATTAT

ATTATACATATATTGAATAATGTTTATAAAAAATGCATAACTTATATGAATATAATTTTTTTTAAATATGACAAAAAGAA

AAAAAAAAAAAACCAAAAAAAATTAAAATTGAAATGAAATATATAAATATATTATTTATATATATTATACATTGTTTAAT

ACTACTACATGTATATATATATATTATATATATATATATATATCAATTTTTTCAAAAATAAATTAATATAAAAAGAGGGG

AAAAAAAAAAAAAAAAAAAAAAAAAGATAATTAAGTAAGCATTTAAAAATATATAAATTGATAATATATAAAATTAATCA

CATATAAACTAATATAATTTATAAAATAAGGAAAATAAAATATTACCATAAAATAAAAATAAAAATAAAAAAAAAAAAAA

AAAACACCTTTTTTTATATATATTAATATATAATTATCTCTTAGAAAAAATATTGTATAATTATATATGTAATGATTTAT

ATAAAAAAATAAAATTATACAAGTATATATTTTGTTTCTATAAATTGATATCTTAATTATTTATTATTAGAAATAGATAT

TTTTATAATAAACCAATAGATAAAATTTGTAGAGAAAAAAAAATAAAAATAAAAATAAAAATAATATAATATATAATAAA

ATAAAATAATATTATATAAATATATTTTAATTTTTTTTACAAAATGGTTGGGGgcgccaccggtTATGTAATAATAATAT

GATCATAATATTATAATAAAACTTATAAAAAAAATATTAAATATTTCATAAATGATTATTATTTATAATAAGATAGATTC

TTTAATTATTTTAAAATTGTATATTTTTTATGTATATTAATTTATTAATATTAATAAGAATATTATAAAAATTCTATTTT

ATTATTTAATGATATTACCCTAATAAAAATAATATAATTATATTACAATATAAATTTATATATATATATATATTTATATA

TTTAAAGAATATTTTATTTTTCAATAAGAACCTTCATTTTAAATTAACATCAAATTATATATATGTATATATACTTCTTA

GTATTATTAATTAAAATACGGAATAATATATAATATATATAAAATGGCAAAACTTTCCTATAGAAAAAAATATTCCATTT

ATTATATTTGTTGTAGGTAATTCTTATTACCGTTTCCTTCTGTTCGTAATGTATATTGGTATGTACTTTATTTTTGCAAT

TTAATTATATGTAAAAAAACGTTAGTACACCATATATATATTATAGTTATAAGAATGCATGCCAAGCCTTTGTCTCAAGA

AGAATCCACCCTCATTGAAAGAGCAACGGCTACAATCAACAGCATCCCCATCTCTGAAGACTACAGCGTCGCCAGCGCAG

CTCTCTCTAGCGACGGCCGCATCTTCACTGGTGTCAATGTATATCATTTTACTGGGGGACCTTGTGCAGAACTCGTGGTG

CTGGGCACTGCTGCTGCTGCGGCAGCTGGCAACCTGACTTGTATCGTCGCGATCGGAAATGAGAACAGGGGCATCTTGAG

CCCCTGCGGACGGTGCCGACAGGTGCTTCTCGATCTGCATCCTGGGATCAAAGCCATAGTGAAGGACAGTGATGGACAGC

CGACGGCAGTTGGGATTCGTGAATTGCTGCCCTCTGGTTATGTGTGGGAGGGCgCTAGcGGCAGTGGAGAGGGCAGAGGA

AGTCTGCTAACATGCGGTGACGTCGAGGAGAATCCTGGCCCAaagcttGAACAAAAACTCATCTCAGAAGAGGATCTGgc

cATGGCACCAAAGGGTAGAAGTACAAATGAAATTGAACTTAGCGCAAGAGATGTTTTGGAAAATATTGGAATAGGAATAT

ATAATCAGGAAAAAATAAAAAAGAATCCATATGAACAACAATTGAAAGGCACATTATCAAACGCCCGATTTCATGATGGC

TTGCACAAGGCAGCTGATTTGGGGGTAATACCTGGTCCTTCACATTTTTCTCAGCTTTATTACAAAAAGCATACTAATAA

CACAAAATATTATAAGGATGATAGGCATCCTTGTCATGGTAGACAAGGAAAACGTTTTGATGAAGGTCAAAAATTTGAAT

GTGGTAATGATAAAATAATTGGTAATAGCGATAAATATGGATCCTGTGCTCCACCTAGAAGAAGACATATATGTGATCAA

AATTTAGAATTCTTAGATAACAATCATACTGATACTATTCATGATGTATTGGGAAATGTGTTGGTCACAGCAAAATATGA

AGGTGAATCTATTGTTAATGATCATCCAGATAAAAAGAACAATGGTAATAAATCAGGTATATGTACTTCTCTTGCACGAA

GTTTTGCCGATATAGGTGATATTGTAAGAGGAAGAGATATGgtcgacaCTAGTACCGGTACGCGTgacgtCAGGTGGCAC

TTTTCGGGGAAATGTGCGCGGAACCCCTATTTGTTTATTTTTCTAAATACATTCAAATATGTATCCGCTCATGAGACAAT

AACCCTGATAAATGCTTCAATAATATTGAAAAAGGAAGAGTATGAGTATTCAACATTTCCGTGTCGCCCTTATTCCCTTT

TTTGCGGCATTTTGCCTTCCTGTTTTTGCTCACCCAGAAACGCTGGTGAAAGTAAAAGATGCTGAAGATCAGTTGGGTGC

ACGAGTGGGTTACATCGAACTGGATCTCAACAGCGGTAAGATCCTTGAGAGTTTTCGCCCCGAAGAACGTTTTCCAATGA

TGAGCACTTTTAAAGTTCTGCTATGTGGCGCGGTATTATCCCGTATTGACGCCGGGCAAGAGCAACTCGGTCGCCGCATA

CACTATTCTCAGAATGACTTGGTTGAGTACTCACCAGTCACAGAAAAGCATCTTACGGATGGCATGACAGTAAGAGAATT

ATGCAGTGCTGCCATAACCATGAGTGATAACACTGCGGCCAACTTACTTCTGACAACGATCGGAGGACCGAAGGAGCTAA

CCGCTTTTTTGCACAACATGGGGGATCATGTAACTCGCCTTGATCGTTGGGAACCGGAGCTGAATGAAGCCATACCAAAC

GACGAGCGTGACACCACGATGCCTGTAGCAATGCCAACAACGTTGCGCAAACTATTAACTGGCGAACTACTTACTCTAGC

TTCCCGGCAACAATTAATAGACTGGATGGAGGCGGATAAAGTTGCAGGACCACTTCTGCGCTCGGCCCTTCCGGCTGGCT

GGTTTATTGCTGATAAATCTGGAGCCGGTGAGCGTGGGTCTCGCGGTATCATTGCAGCACTGGGGCCAGATGGTAAGCCC

TCCCGTATCGTAGTTATCTACACGACGGGGAGTCAGGCAACTATGGATGAACGAAATAGACAGATCGCTGAGATAGGTGC

CTCACTGATTAAGCATTGGTAACTGTCAGACCAAGTTTACTCATATATACTTTAGATTGATTTAAAACTTCATTTTTAAT

TTAAAAGGATCTAGGTGAAGATCCTTTTTGATAATCTCATGACCAAAATCCCTTAACGTGAGTTTTCGTTCCACTGAGCG

TCAGACCCCGTAGAAAAGATCAAAGGATCTTCTTGAGATCCTTTTTTTCTGCGCGTAATCTGCTGCTTGCAAACAAAAAA

ACCACCGCTACCAGCGGTGGTTTGTTTGCCGGATCAAGAGCTACCAACTCTTTTTCCGAAGGTAACTGGCTTCAGCAGAG

CGCAGATACCAAATACTGTCCTTCTAGTGTAGCCGTAGTTAGGCCACCACTTCAAGAACTCTGTAGCACCGCCTACATAC

CTCGCTCTGCTAATCCTGTTACCAGTGGCTGCTGCCAGTGGCGATAAGTCGTGTCTTACCGGGTTGGACTCAAGACGATA

GTTACCGGATAAGGCGCAGCGGTCGGGCTGAACGGGGGGTTCGTGCACACAGCCCAGCTTGGAGCGAACGACCTACACCG

AACTGAGATACCTACAGCGTGAGCTATGAGAAAGCGCCACGCTTCCCGAAGGGAGAAAGGCGGACAGGTATCCGGTAAGC

GGCAGGGTCGGAACAGGAGAGCGCACGAGGGAGCTTCCAGGGGGAAACGCCTGGTATCTTTATAGTCCTGTCGGGTTTCG

CCACCTCTGACTTGAGCGTCGATTTTTGTGATGCTCGTCAGGGGGGCGGAGCCTATCGAAAAACGCCAGCAACGCGGCCT

TTTTACGGTTCCTGGCCTTTTGCTGGCCTTTTGCTCACATGTTCTTTCCTGCGTTATCCCCTGATTCTGTGGATAACCGT

ATTACCGCCTTTGAGTGAGCTGATACCGCTCGCCGCAGCCGAACGACCGAGCGCAGCGAGTCAGTGAGCGAGGAAGCGGA

AGAGCGCCCAATACGCAAACCGCCTCTCCCCGCGCGTTGGCCGATTCATTAATGCAGCTGGCACGACAGGTTTCCCGACT

GGAAAGCGGGCAGTGAGCGCAACGCAATTAATGTGAGTTAGCTCACTCATTAGGCACCCCAGGCTTTACACTTTATGCTT

CCGGCTCGTATGTTGTGTGGAATTGTGAGCGGATAACAATTTCACACAGGAAACAGCTATGACCATGATTACGCCAAGCT

ATTTAGGTGACACTATAGAATACTC

Nucleotide sequence of plasmid Cas9_itvar60prom_bsd_2A_3xFLAG_exonI

AAGCTTGGGGGGATCCGCCTTAAAAACTTCATTATATTTAAAAATTATTTTATAGGAAATAATAAAAAAAAAAgcaccga

ctcggtgccactttttcaagttgataacggactagccttattttaacttgctaTTTCtagctctaaaacTTTGGTGCCAT

TCTTATAACAATATTATATACTTAATATGAAATATGTGCATATAGGAAAAATTATGCATTTTGGTTACTCTAATATTATA

TATATATATATATATATATATATATTATAATATATTATGTTATATATACATAACATATACATTTTTTAATAATAATTTAC

CCTTTATTTTTACATTATAAAAAATTATATTACAGTAAAAATAAAAGTTTATTATATTAATAGTTTTTTTTTTTTTTTTT

TTAATTTATGAAATATTTAAATATTTAAAATTTTTTAAATGAATAATTATATTTATAATTAGAAAAAAAAAAAAAAAAAA

AAAAAAAAAATATAGCTATTTATATAAATTTCTTTTATTTATCTGAACAAGCAAGAATTTTTTTTTATATTAAATTAGAA

TAAATTATTATTAGTTTATGTATATATTTTTTTTTTTTCATAGTATATAAATATTATATATATTGTACCTTTTTACAATA

TATTTCATATATAGAAGAGAAAAAAAAAAAAAGAAGATATTATTGTAAAACCTCAAGATGTGTAGAAATCCAAATGTCGG

ATCCTCTAGAGTCGACCTGCAGGCATGCTATTTGATGAATTAACTACACTTAAAATAATACAATTATTATTAAATTTTTT

TTTGATTTATTTATTAATTTTTAAACTTAATCATTTGTATTTGGGAGGAATTATATATATCTTTATAATTATTTTATTTT

TTTTTATTTTTTTATTTTTTTATTATTATTATTTTTTTTTATTTTTTTTTTTTACTGTATCAAAGAAAAACCTTTAAAAA

AAAAATTATAATTTCCCCATCTTACTATATTTTTAATACATACGTTTTAAGGAATTAAATTAGACAAAAGCTATATTATG

CTTTACATATAATTAGAATTTATAAACGTTTGGTTATTAGATATTTCATGTCTCAGTAAAGTCTTTCAATACATATGTAA

AAAAATATATATGAATACACATAAGTTGTTAATATATTTTATATGCATAAATGTATAAATATATATATATATATATATAT

ATGTATGTATGTATATGTGTGTATATGAAATTATTTCAATGTTTAATTTTTTAAATTTTAATTTTTTTTTTTTTTTTTTT

TTTTATTATGTATATTGATCTTTATTATTTAAATATTACTTTTTTCGTTTTTTCTTCTTTTTATTATTTTTTTTTTTTTT

TATATTTTATACAAATGGTAATTCAAATAAAAGGTATAAATTTATATTTAATTTTCTTTTATGGATAAATAAAAGAAAAA

TATAAATATATAAAAATATAAAAATATATATATGTATATTGGGGTGATGATAAAATGAAAGATAATATATATATATATAT

ATCTTTATTTTTTTTTTTTTGTAGACCCCATTGTGAGTACATAAATATATTATATAACTCGAGTTACTTTTTCTTTTTTG

CCTGGCCGGCCTTTTTCGTGGCCGCCGGCCTTTTGTCGCCTCCCAGCTGAGACAGGTCGATCCGTGTCTCGTACAGGCCG

GTGATGCTCTGGTGGATCAGGGTGGCGTCCAGCACCTCTTTGGTGCTGGTGTACCTCTTCCGGTCGATGGTGGTGTCAAA

GTACTTGAAGGCGGCAGGGGCTCCCAGATTGGTCAGGGTAAACAGGTGGATGATATTCTCGGCCTGCTCTCTGATGGGCT

TATCCCGGTGCTTGTTGTAGGCGGACAGCACTTTGTCCAGATTAGCGTCGGCCAGGATCACTCTCTTGGAGAACTCGCTG

ATCTGCTCGATGATCTCGTCCAGGTAGTGCTTGTGCTGTTCCACAAACAGCTGTTTCTGCTCATTATCCTCGGGGGAGCC

CTTCAGCTTCTCATAGTGGCTGGCCAGGTACAGGAAGTTCACATATTTGGAGGGCAGGGCCAGTTCGTTTCCCTTCTGCA

GTTCGCCGGCAGAGGCCAGCATTCTCTTCCGGCCGTTTTCCAGCTCGAACAGGGAGTACTTAGGCAGCTTGATGATCAGG

TCCTTTTTCACTTCTTTGTAGCCCTTGGCTTCCAGAAAGTCGATGGGATTCTTCTCGAAGCTGCTTCTTTCCATGATGGT

GATCCCCAGCAGCTCTTTCACACTCTTCAGTTTCTTGGACTTGCCCTTTTCCACTTTGGCCACCACCAGCACAGAATAGG

CCACGGTGGGGCTGTCGAAGCCGCCGTACTTCTTAGGGTCCCAGTCCTTCTTTCTGGCGATCAGCTTATCGCTGTTCCTC

TTGGGCAGGATAGACTCTTTGCTGAAGCCGCCTGTCTGCACCTCGGTCTTTTTCACGATATTCACTTGGGGCATGCTCAG

CACTTTCCGCACGGTGGCAAAATCCCGGCCCTTATCCCACACGATCTCCCCGGTTTCGCCGTTTGTCTCGATCAGAGGCC

GCTTCCGGATCTCGCCGTTGGCCAGGGTAATCTCGGTCTTGAAAAAGTTCATGATGTTGCTGTAGAAGAAGTACTTGGCG

GTAGCCTTGCCGATTTCCTGCTCGCTCTTGGCGATCATCTTCCGCACGTCGTACACCTTGTAGTCGCCGTACACGAACTC

GCTTTCCAGCTTAGGGTACTTTTTGATCAGGGCGGTTCCCACGACGGCGTTCAGGTAGGCGTCGTGGGCGTGGTGGTAGT

TGTTGATCTCGCGCACTTTGTAAAACTGGAAATCCTTCCGGAAATCGGACACCAGCTTGGACTTCAGGGTGATCACTTTC

ACTTCCCGGATCAGCTTGTCATTCTCGTCGTACTTAGTGTTCATCCGGGAGTCCAGGATCTGTGCCACGTGCTTTGTGAT

CTGCCGGGTTTCCACCAGCTGTCTCTTGATGAAGCCGGCCTTATCCAGTTCGCTCAGGCCGCCTCTCTCGGCCTTGGTCA

GATTGTCGAACTTTCTCTGGGTAATCAGCTTGGCGTTCAGCAGCTGCCGCCAGTAGTTCTTCATCTTCTTCACGACCTCT

TCGGAGGGCACGTTGTCGCTCTTGCCCCGGTTCTTGTCGCTTCTGGTCAGCACCTTGTTGTCGATGGAGTCGTCCTTCAG

AAAGCTCTGAGGCACGATATGGTCCACATCGTAGTCGGACAGCCGGTTGATGTCCAGTTCCTGGTCCACGTACATATCCC

GCCCATTCTGCAGGTAGTACAGGTACAGCTTCTCGTTCTGCAGCTGGGTGTTTTCCACGGGGTGTTCTTTCAGGATCTGG

CTGCCCAGCTCTTTGATGCCCTCTTCGATCCGCTTCATTCTCTCGCGGCTGTTCTTCTGTCCCTTCTGGGTGGTCTGGTT

CTCTCTGGCCATTTCGATCACGATGTTCTCGGGCTTGTGCCGGCCCATCACTTTCACGAGCTCGTCCACCACCTTCACTG

TCTGCAGGATGCCCTTCTTAATGGCGGGGCTGCCGGCCAGATTGGCAATGTGCTCGTGCAGGCTATCGCCCTGGCCGGAC

ACCTGGGCTTTCTGGATGTCCTCTTTAAAGGTCAGGCTGTCGTCGTGGATCAGCTGCATGAAGTTTCTGTTGGCGAAGCC

GTCGGACTTCAGGAAATCCAGGATTGTCTTGCCGGACTGCTTGTCCCGGATGCCGTTGATCAGCTTCCGGCTCAGCCTGC

CCCAGCCGGTGTATCTCCGCCGCTTCAGCTGCTTCATCACTTTGTCGTCGAACAGGTGGGCATAGGTTTTCAGCCGTTCC

TCGATCATCTCTCTGTCCTCAAACAGTGTCAGGGTCAGCACGATATCTTCCAGAATGTCCTCGTTTTCCTCATTGTCCAG

GAAGTCCTTGTCCTTGATAATTTTCAGCAGATCGTGGTATGTGCCCAGGGAGGCGTTGAACCGATCTTCCACGCCGGAGA

TTTCCACGGAGTCGAAGCACTCGATTTTCTTGAAGTAGTCCTCTTTCAGCTGCTTCACGGTCACTTTCCGGTTGGTCTTG

AACAGCAGGTCCACGATGGCCTTTTTCTGCTCGCCGCTCAGGAAGGCGGGCTTTCTCATTCCCTCGGTCACGTATTTCAC

TTTGGTCAGCTCGTTATACACGGTGAAGTACTCGTACAGCAGGCTGTGCTTGGGCAGCACCTTCTCGTTGGGCAGGTTCT

TATCGAAGTTGGTCATCCGCTCGATGAAGCTCTGGGCGGAAGCGCCCTTGTCCACCACTTCCTCGAAGTTCCAGGGGGTG

ATGGTTTCCTCGCTCTTTCTGGTCATCCAGGCGAATCTGCTGTTTCCCCTGGCCAGAGGGCCCACGTAGTAGGGGATGCG

GAAGGTCAGGATCTTCTCGATCTTTTCCCGGTTGTCCTTCAGGAATGGGTAAAAATCTTCCTGCCGCCGCAGAATGGCGT

GCAGCTCTCCCAGGTGGATCTGGTGGGGGATGCTGCCGTTGTCGAAGGTCCGCTGCTTCCGCAGCAGGTCCTCTCTGTTC

AGCTTCACGAGCAGTTCCTCGGTGCCGTCCATCTTTTCCAGGATGGGCTTGATGAACTTGTAGAACTCTTCCTGGCTGGC

TCCGCCGTCAATGTAGCCGGCGTAGCCGTTCTTGCTCTGGTCGAAGAAAATCTCTTTGTACTTCTCAGGCAGCTGCTGCC

GCACGAGAGCTTTCAGCAGGGTCAGGTCCTGGTGGTGCTCGTCGTATCTCTTGATCATAGAGGCGCTCAGGGGGGCCTTG

GTGATCTCGGTGTTCACTCTCAGGATGTCGCTCAGCAGGATGGCGTCGGACAGGTTCTTGGCGGCCAGAAACAGGTCGGC

GTACTGGTCGCCGATCTGGGCCAGCAGGTTGTCCAGGTCGTCGTCGTAGGTGTCCTTGCTCAGCTGCAGTTTGGCATCCT

CGGCCAGGTCGAAGTTGCTCTTGAAGTTGGGGGTCAGGCCCAGGCTCAGGGCAATCAGGTTTCCGAACAGGCCATTCTTC

TTCTCGCCGGGCAGCTGGGCGATCAGATTTTCCAGCCGTCTGCTCTTGCTCAGTCTGGCAGACAGGATGGCCTTGGCGTC

CACGCCGCTGGCGTTGATGGGGTTTTCCTCGAACAGCTGGTTGTAGGTCTGCACCAGCTGGATGAACAGCTTGTCCACGT

CGCTGTTGTCGGGGTTCAGGTCGCCCTCGATCAGGAAGTGGCCCCGGAACTTGATCATGTGGGCCAGGGCCAGATAGATC

AGCCGCAGGTCGGCCTTGTCGGTGCTGTCCACCAGTTTCTTTCTCAGGTGGTAGATGGTGGGGTACTTCTCGTGGTAGGC

CACCTCGTCCACGATGTTGCCGAAGATGGGGTGCCGCTCGTGCTTCTTATCCTCTTCCACCAGGAAGGACTCTTCCAGTC

TGTGGAAGAAGCTGTCGTCCACCTTGGCCATCTCGTTGCTGAAGATCTCTTGCAGATAGCAGATCCGGTTCTTCCGTCTG

GTGTATCTTCTTCTGGCGGTTCTCTTCAGCCGGGTGGCCTCGGCTGTTTCGCCGCTGTCGAACAGCAGGGCTCCGATCAG

GTTCTTCTTGATGCTGTGCCGGTCGGTGTTGCCCAGCACCTTGAATTTCTTGCTGGGCACCTTGTACTCGTCGGTGATCA

CGGCCCAGCCCACAGAGTTGGTGCCGATGTCCAGGCCGATGCTGTACTTCTTGTCGGCTGCTGGGACTCCGTGGATACCG

ACCTTCCGCTTCTTCTTTGGGGCCATCTTATCGTCATCGTCTTTGTAATCAATATCATGATCCTTGTAGTCTCCGTCGTG

GTCCTTATAGTCCATcctaggTGATATATTTCTATTAGGTATTTATTATTATAAAATATAAATCTTGAATGATAATAAAT

AAAATATTAGTTATTCCTTTTCTAGTTTAAAATATACATATTATAAATATATATATATATATATATATTTTTATTGTGAC

AAGAATATATAATTATAAATTATATTATTTATTTTTGTATTTTTTTTTTTTTTTTTTTTTTTTTCTTTTTTTGTTTTATT

TTTCTTTTTTTTTATAAATATTATTTTTTTCTTTTATCATGCACATTGGAATAATACATTAATATATATATATATATTAT

ATTATACATATATTGAATAATGTTTATAAAAAATGCATAACTTATATGAATATAATTTTTTTTAAATATGACAAAAAGAA

AAAAAAAAAAAACCAAAAAAAATTAAAATTGAAATGAAATATATAAATATATTATTTATATATATTATACATTGTTTAAT

ACTACTACATGTATATATATATATTATATATATATATATATATCAATTTTTTCAAAAATAAATTAATATAAAAAGAGGGG

AAAAAAAAAAAAAAAAAAAAAAAAAGATAATTAAGTAAGCATTTAAAAATATATAAATTGATAATATATAAAATTAATCA

CATATAAACTAATATAATTTATAAAATAAGGAAAATAAAATATTACCATAAAATAAAAATAAAAATAAAAAAAAAAAAAA

AAAACACCTTTTTTTATATATATTAATATATAATTATCTCTTAGAAAAAATATTGTATAATTATATATGTAATGATTTAT

ATAAAAAAATAAAATTATACAAGTATATATTTTGTTTCTATAAATTGATATCTTAATTATTTATTATTAGAAATAGATAT

TTTTATAATAAACCAATAGATAAAATTTGTAGAGAAAAAAAAATAAAAATAAAAATAAAAATAATATAATATATAATAAA

ATAAAATAATATTATATAAATATATTTTAATTTTTTTTACAAAATGGTTGGGGgcgccaccggtTATGTAATAATAATAT

GATCATAATATTATAATAAAACTTATAAAAAAAATATTAAATATTTCATAAATGATTATTATTTATAATAAGATAGATTC

TTTAATTATTTTAAAATTGTATATTTTTTATGTATATTAATTTATTAATATTAATAAGAATATTATAAAAATTCTATTTT

ATTATTTAATGATATTACCCTAATAAAAATAATATAATTATATTACAATATAAATTTATATATATATATATATTTATATA

TTTAAAGAATATTTTATTTTTCAATAAGAACCTTCATTTTAAATTAACATCAAATTATATATATGTATATATACTTCTTA

GTATTATTAATTAAAATACGGAATAATATATAATATATATAAAATGGCAAAACTTTCCTATAGAAAAAAATATTCCATTT

ATTATATTTGTTGTAGGTAATTCTTATTACCGTTTCCTTCTGTTCGTAATGTATATTGGTATGTACTTTATTTTTGCAAT

TTAATTATATGTAAAAAAACGTTAGTACACCATATATATATTATAGTTATAAGAATGCATGCCAAGCCTTTGTCTCAAGA

AGAATCCACCCTCATTGAAAGAGCAACGGCTACAATCAACAGCATCCCCATCTCTGAAGACTACAGCGTCGCCAGCGCAG

CTCTCTCTAGCGACGGCCGCATCTTCACTGGTGTCAATGTATATCATTTTACTGGGGGACCTTGTGCAGAACTCGTGGTG

CTGGGCACTGCTGCTGCTGCGGCAGCTGGCAACCTGACTTGTATCGTCGCGATCGGAAATGAGAACAGGGGCATCTTGAG

CCCCTGCGGACGGTGCCGACAGGTGCTTCTCGATCTGCATCCTGGGATCAAAGCCATAGTGAAGGACAGTGATGGACAGC

CGACGGCAGTTGGGATTCGTGAATTGCTGCCCTCTGGTTATGTGTGGGAGGGCgCTAGcGGCAGTGGAGAGGGCAGAGGA

AGTCTGCTAACATGCGGTGACGTCGAGGAGAATCCTGGCCCAaagcttGACTACAAGGACCACGACGGTGACTACAAGGA

CCACGACATCGACTACAAGGACGACGACGACAAGgccATGGCACCAAAGGGTAGAAGTACAAATGAAATTGAACTTAGCG

CAAGAGATGTTTTGGAAAATATTGGAATAGGAATATATAATCAGGAAAAAATAAAAAAGAATCCATATGAACAACAATTG

AAAGGCACATTATCAAACGCCCGATTTCATGATGGCTTGCACAAGGCAGCTGATTTGGGGGTAATACCTGGTCCTTCACA

TTTTTCTCAGCTTTATTACAAAAAGCATACTAATAACACAAAATATTATAAGGATGATAGGCATCCTTGTCATGGTAGAC

AAGGAAAACGTTTTGATGAAGGTCAAAAATTTGAATGTGGTAATGATAAAATAATTGGTAATAGCGATAAATATGGATCC

TGTGCTCCACCTAGAAGAAGACATATATGTGATCAAAATTTAGAATTCTTAGATAACAATCATACTGATACTATTCATGA

TGTATTGGGAAATGTGTTGGTCACAGCAAAATATGAAGGTGAATCTATTGTTAATGATCATCCAGATAAAAAGAACAATG

GTAATAAATCAGGTATATGTACTTCTCTTGCACGAAGTTTTGCCGATATAGGTGATATTGTAAGAGGAAGAGATATGgtc

gacaCTAGTACCGGTACGCGTgacgtCAGGTGGCACTTTTCGGGGAAATGTGCGCGGAACCCCTATTTGTTTATTTTTCT

AAATACATTCAAATATGTATCCGCTCATGAGACAATAACCCTGATAAATGCTTCAATAATATTGAAAAAGGAAGAGTATG

AGTATTCAACATTTCCGTGTCGCCCTTATTCCCTTTTTTGCGGCATTTTGCCTTCCTGTTTTTGCTCACCCAGAAACGCT

GGTGAAAGTAAAAGATGCTGAAGATCAGTTGGGTGCACGAGTGGGTTACATCGAACTGGATCTCAACAGCGGTAAGATCC

TTGAGAGTTTTCGCCCCGAAGAACGTTTTCCAATGATGAGCACTTTTAAAGTTCTGCTATGTGGCGCGGTATTATCCCGT

ATTGACGCCGGGCAAGAGCAACTCGGTCGCCGCATACACTATTCTCAGAATGACTTGGTTGAGTACTCACCAGTCACAGA

AAAGCATCTTACGGATGGCATGACAGTAAGAGAATTATGCAGTGCTGCCATAACCATGAGTGATAACACTGCGGCCAACT

TACTTCTGACAACGATCGGAGGACCGAAGGAGCTAACCGCTTTTTTGCACAACATGGGGGATCATGTAACTCGCCTTGAT

CGTTGGGAACCGGAGCTGAATGAAGCCATACCAAACGACGAGCGTGACACCACGATGCCTGTAGCAATGCCAACAACGTT

GCGCAAACTATTAACTGGCGAACTACTTACTCTAGCTTCCCGGCAACAATTAATAGACTGGATGGAGGCGGATAAAGTTG

CAGGACCACTTCTGCGCTCGGCCCTTCCGGCTGGCTGGTTTATTGCTGATAAATCTGGAGCCGGTGAGCGTGGGTCTCGC

GGTATCATTGCAGCACTGGGGCCAGATGGTAAGCCCTCCCGTATCGTAGTTATCTACACGACGGGGAGTCAGGCAACTAT

GGATGAACGAAATAGACAGATCGCTGAGATAGGTGCCTCACTGATTAAGCATTGGTAACTGTCAGACCAAGTTTACTCAT

ATATACTTTAGATTGATTTAAAACTTCATTTTTAATTTAAAAGGATCTAGGTGAAGATCCTTTTTGATAATCTCATGACC

AAAATCCCTTAACGTGAGTTTTCGTTCCACTGAGCGTCAGACCCCGTAGAAAAGATCAAAGGATCTTCTTGAGATCCTTT

TTTTCTGCGCGTAATCTGCTGCTTGCAAACAAAAAAACCACCGCTACCAGCGGTGGTTTGTTTGCCGGATCAAGAGCTAC

CAACTCTTTTTCCGAAGGTAACTGGCTTCAGCAGAGCGCAGATACCAAATACTGTCCTTCTAGTGTAGCCGTAGTTAGGC

CACCACTTCAAGAACTCTGTAGCACCGCCTACATACCTCGCTCTGCTAATCCTGTTACCAGTGGCTGCTGCCAGTGGCGA

TAAGTCGTGTCTTACCGGGTTGGACTCAAGACGATAGTTACCGGATAAGGCGCAGCGGTCGGGCTGAACGGGGGGTTCGT

GCACACAGCCCAGCTTGGAGCGAACGACCTACACCGAACTGAGATACCTACAGCGTGAGCTATGAGAAAGCGCCACGCTT

CCCGAAGGGAGAAAGGCGGACAGGTATCCGGTAAGCGGCAGGGTCGGAACAGGAGAGCGCACGAGGGAGCTTCCAGGGGG

AAACGCCTGGTATCTTTATAGTCCTGTCGGGTTTCGCCACCTCTGACTTGAGCGTCGATTTTTGTGATGCTCGTCAGGGG

GGCGGAGCCTATCGAAAAACGCCAGCAACGCGGCCTTTTTACGGTTCCTGGCCTTTTGCTGGCCTTTTGCTCACATGTTC

TTTCCTGCGTTATCCCCTGATTCTGTGGATAACCGTATTACCGCCTTTGAGTGAGCTGATACCGCTCGCCGCAGCCGAAC

GACCGAGCGCAGCGAGTCAGTGAGCGAGGAAGCGGAAGAGCGCCCAATACGCAAACCGCCTCTCCCCGCGCGTTGGCCGA

TTCATTAATGCAGCTGGCACGACAGGTTTCCCGACTGGAAAGCGGGCAGTGAGCGCAACGCAATTAATGTGAGTTAGCTC

ACTCATTAGGCACCCCAGGCTTTACACTTTATGCTTCCGGCTCGTATGTTGTGTGGAATTGTGAGCGGATAACAATTTCA

CACAGGAAACAGCTATGACCATGATTACGCCAAGCTATTTAGGTGACACTATAGAATACTC

Nucleotide sequence of plasmid Cas9_it4var60prom_bsd_2A_exonI_KO

AAGCTTGGGGGGATCCGCCTTAAAAACTTCATTATATTTAAAAATTATTTTATAGGAAATAATAAAAAAAAAAgcaccga

ctcggtgccactttttcaagttgataacggactagccttattttaacttgctaTTTCtagctctaaaacTTTGGTGCCAT

TCTTATAACAATATTATATACTTAATATGAAATATGTGCATATAGGAAAAATTATGCATTTTGGTTACTCTAATATTATA

TATATATATATATATATATATATATTATAATATATTATGTTATATATACATAACATATACATTTTTTAATAATAATTTAC

CCTTTATTTTTACATTATAAAAAATTATATTACAGTAAAAATAAAAGTTTATTATATTAATAGTTTTTTTTTTTTTTTTT

TTAATTTATGAAATATTTAAATATTTAAAATTTTTTAAATGAATAATTATATTTATAATTAGAAAAAAAAAAAAAAAAAA

AAAAAAAAAATATAGCTATTTATATAAATTTCTTTTATTTATCTGAACAAGCAAGAATTTTTTTTTATATTAAATTAGAA

TAAATTATTATTAGTTTATGTATATATTTTTTTTTTTTCATAGTATATAAATATTATATATATTGTACCTTTTTACAATA

TATTTCATATATAGAAGAGAAAAAAAAAAAAAGAAGATATTATTGTAAAACCTCAAGATGTGTAGAAATCCAAATGTCGG

ATCCTCTAGAGTCGACCTGCAGGCATGCTATTTGATGAATTAACTACACTTAAAATAATACAATTATTATTAAATTTTTT

TTTGATTTATTTATTAATTTTTAAACTTAATCATTTGTATTTGGGAGGAATTATATATATCTTTATAATTATTTTATTTT

TTTTTATTTTTTTATTTTTTTATTATTATTATTTTTTTTTATTTTTTTTTTTTACTGTATCAAAGAAAAACCTTTAAAAA

AAAAATTATAATTTCCCCATCTTACTATATTTTTAATACATACGTTTTAAGGAATTAAATTAGACAAAAGCTATATTATG

CTTTACATATAATTAGAATTTATAAACGTTTGGTTATTAGATATTTCATGTCTCAGTAAAGTCTTTCAATACATATGTAA

AAAAATATATATGAATACACATAAGTTGTTAATATATTTTATATGCATAAATGTATAAATATATATATATATATATATAT

ATGTATGTATGTATATGTGTGTATATGAAATTATTTCAATGTTTAATTTTTTAAATTTTAATTTTTTTTTTTTTTTTTTT

TTTTATTATGTATATTGATCTTTATTATTTAAATATTACTTTTTTCGTTTTTTCTTCTTTTTATTATTTTTTTTTTTTTT

TATATTTTATACAAATGGTAATTCAAATAAAAGGTATAAATTTATATTTAATTTTCTTTTATGGATAAATAAAAGAAAAA

TATAAATATATAAAAATATAAAAATATATATATGTATATTGGGGTGATGATAAAATGAAAGATAATATATATATATATAT

ATCTTTATTTTTTTTTTTTTGTAGACCCCATTGTGAGTACATAAATATATTATATAACTCGAGTTACTTTTTCTTTTTTG

CCTGGCCGGCCTTTTTCGTGGCCGCCGGCCTTTTGTCGCCTCCCAGCTGAGACAGGTCGATCCGTGTCTCGTACAGGCCG

GTGATGCTCTGGTGGATCAGGGTGGCGTCCAGCACCTCTTTGGTGCTGGTGTACCTCTTCCGGTCGATGGTGGTGTCAAA

GTACTTGAAGGCGGCAGGGGCTCCCAGATTGGTCAGGGTAAACAGGTGGATGATATTCTCGGCCTGCTCTCTGATGGGCT

TATCCCGGTGCTTGTTGTAGGCGGACAGCACTTTGTCCAGATTAGCGTCGGCCAGGATCACTCTCTTGGAGAACTCGCTG

ATCTGCTCGATGATCTCGTCCAGGTAGTGCTTGTGCTGTTCCACAAACAGCTGTTTCTGCTCATTATCCTCGGGGGAGCC

CTTCAGCTTCTCATAGTGGCTGGCCAGGTACAGGAAGTTCACATATTTGGAGGGCAGGGCCAGTTCGTTTCCCTTCTGCA

GTTCGCCGGCAGAGGCCAGCATTCTCTTCCGGCCGTTTTCCAGCTCGAACAGGGAGTACTTAGGCAGCTTGATGATCAGG

TCCTTTTTCACTTCTTTGTAGCCCTTGGCTTCCAGAAAGTCGATGGGATTCTTCTCGAAGCTGCTTCTTTCCATGATGGT

GATCCCCAGCAGCTCTTTCACACTCTTCAGTTTCTTGGACTTGCCCTTTTCCACTTTGGCCACCACCAGCACAGAATAGG

CCACGGTGGGGCTGTCGAAGCCGCCGTACTTCTTAGGGTCCCAGTCCTTCTTTCTGGCGATCAGCTTATCGCTGTTCCTC

TTGGGCAGGATAGACTCTTTGCTGAAGCCGCCTGTCTGCACCTCGGTCTTTTTCACGATATTCACTTGGGGCATGCTCAG

CACTTTCCGCACGGTGGCAAAATCCCGGCCCTTATCCCACACGATCTCCCCGGTTTCGCCGTTTGTCTCGATCAGAGGCC

GCTTCCGGATCTCGCCGTTGGCCAGGGTAATCTCGGTCTTGAAAAAGTTCATGATGTTGCTGTAGAAGAAGTACTTGGCG

GTAGCCTTGCCGATTTCCTGCTCGCTCTTGGCGATCATCTTCCGCACGTCGTACACCTTGTAGTCGCCGTACACGAACTC

GCTTTCCAGCTTAGGGTACTTTTTGATCAGGGCGGTTCCCACGACGGCGTTCAGGTAGGCGTCGTGGGCGTGGTGGTAGT

TGTTGATCTCGCGCACTTTGTAAAACTGGAAATCCTTCCGGAAATCGGACACCAGCTTGGACTTCAGGGTGATCACTTTC

ACTTCCCGGATCAGCTTGTCATTCTCGTCGTACTTAGTGTTCATCCGGGAGTCCAGGATCTGTGCCACGTGCTTTGTGAT

CTGCCGGGTTTCCACCAGCTGTCTCTTGATGAAGCCGGCCTTATCCAGTTCGCTCAGGCCGCCTCTCTCGGCCTTGGTCA

GATTGTCGAACTTTCTCTGGGTAATCAGCTTGGCGTTCAGCAGCTGCCGCCAGTAGTTCTTCATCTTCTTCACGACCTCT

TCGGAGGGCACGTTGTCGCTCTTGCCCCGGTTCTTGTCGCTTCTGGTCAGCACCTTGTTGTCGATGGAGTCGTCCTTCAG

AAAGCTCTGAGGCACGATATGGTCCACATCGTAGTCGGACAGCCGGTTGATGTCCAGTTCCTGGTCCACGTACATATCCC

GCCCATTCTGCAGGTAGTACAGGTACAGCTTCTCGTTCTGCAGCTGGGTGTTTTCCACGGGGTGTTCTTTCAGGATCTGG

CTGCCCAGCTCTTTGATGCCCTCTTCGATCCGCTTCATTCTCTCGCGGCTGTTCTTCTGTCCCTTCTGGGTGGTCTGGTT

CTCTCTGGCCATTTCGATCACGATGTTCTCGGGCTTGTGCCGGCCCATCACTTTCACGAGCTCGTCCACCACCTTCACTG

TCTGCAGGATGCCCTTCTTAATGGCGGGGCTGCCGGCCAGATTGGCAATGTGCTCGTGCAGGCTATCGCCCTGGCCGGAC

ACCTGGGCTTTCTGGATGTCCTCTTTAAAGGTCAGGCTGTCGTCGTGGATCAGCTGCATGAAGTTTCTGTTGGCGAAGCC

GTCGGACTTCAGGAAATCCAGGATTGTCTTGCCGGACTGCTTGTCCCGGATGCCGTTGATCAGCTTCCGGCTCAGCCTGC

CCCAGCCGGTGTATCTCCGCCGCTTCAGCTGCTTCATCACTTTGTCGTCGAACAGGTGGGCATAGGTTTTCAGCCGTTCC

TCGATCATCTCTCTGTCCTCAAACAGTGTCAGGGTCAGCACGATATCTTCCAGAATGTCCTCGTTTTCCTCATTGTCCAG

GAAGTCCTTGTCCTTGATAATTTTCAGCAGATCGTGGTATGTGCCCAGGGAGGCGTTGAACCGATCTTCCACGCCGGAGA

TTTCCACGGAGTCGAAGCACTCGATTTTCTTGAAGTAGTCCTCTTTCAGCTGCTTCACGGTCACTTTCCGGTTGGTCTTG

AACAGCAGGTCCACGATGGCCTTTTTCTGCTCGCCGCTCAGGAAGGCGGGCTTTCTCATTCCCTCGGTCACGTATTTCAC

TTTGGTCAGCTCGTTATACACGGTGAAGTACTCGTACAGCAGGCTGTGCTTGGGCAGCACCTTCTCGTTGGGCAGGTTCT

TATCGAAGTTGGTCATCCGCTCGATGAAGCTCTGGGCGGAAGCGCCCTTGTCCACCACTTCCTCGAAGTTCCAGGGGGTG

ATGGTTTCCTCGCTCTTTCTGGTCATCCAGGCGAATCTGCTGTTTCCCCTGGCCAGAGGGCCCACGTAGTAGGGGATGCG

GAAGGTCAGGATCTTCTCGATCTTTTCCCGGTTGTCCTTCAGGAATGGGTAAAAATCTTCCTGCCGCCGCAGAATGGCGT

GCAGCTCTCCCAGGTGGATCTGGTGGGGGATGCTGCCGTTGTCGAAGGTCCGCTGCTTCCGCAGCAGGTCCTCTCTGTTC

AGCTTCACGAGCAGTTCCTCGGTGCCGTCCATCTTTTCCAGGATGGGCTTGATGAACTTGTAGAACTCTTCCTGGCTGGC

TCCGCCGTCAATGTAGCCGGCGTAGCCGTTCTTGCTCTGGTCGAAGAAAATCTCTTTGTACTTCTCAGGCAGCTGCTGCC

GCACGAGAGCTTTCAGCAGGGTCAGGTCCTGGTGGTGCTCGTCGTATCTCTTGATCATAGAGGCGCTCAGGGGGGCCTTG

GTGATCTCGGTGTTCACTCTCAGGATGTCGCTCAGCAGGATGGCGTCGGACAGGTTCTTGGCGGCCAGAAACAGGTCGGC

GTACTGGTCGCCGATCTGGGCCAGCAGGTTGTCCAGGTCGTCGTCGTAGGTGTCCTTGCTCAGCTGCAGTTTGGCATCCT

CGGCCAGGTCGAAGTTGCTCTTGAAGTTGGGGGTCAGGCCCAGGCTCAGGGCAATCAGGTTTCCGAACAGGCCATTCTTC

TTCTCGCCGGGCAGCTGGGCGATCAGATTTTCCAGCCGTCTGCTCTTGCTCAGTCTGGCAGACAGGATGGCCTTGGCGTC

CACGCCGCTGGCGTTGATGGGGTTTTCCTCGAACAGCTGGTTGTAGGTCTGCACCAGCTGGATGAACAGCTTGTCCACGT

CGCTGTTGTCGGGGTTCAGGTCGCCCTCGATCAGGAAGTGGCCCCGGAACTTGATCATGTGGGCCAGGGCCAGATAGATC

AGCCGCAGGTCGGCCTTGTCGGTGCTGTCCACCAGTTTCTTTCTCAGGTGGTAGATGGTGGGGTACTTCTCGTGGTAGGC

CACCTCGTCCACGATGTTGCCGAAGATGGGGTGCCGCTCGTGCTTCTTATCCTCTTCCACCAGGAAGGACTCTTCCAGTC

TGTGGAAGAAGCTGTCGTCCACCTTGGCCATCTCGTTGCTGAAGATCTCTTGCAGATAGCAGATCCGGTTCTTCCGTCTG

GTGTATCTTCTTCTGGCGGTTCTCTTCAGCCGGGTGGCCTCGGCTGTTTCGCCGCTGTCGAACAGCAGGGCTCCGATCAG

GTTCTTCTTGATGCTGTGCCGGTCGGTGTTGCCCAGCACCTTGAATTTCTTGCTGGGCACCTTGTACTCGTCGGTGATCA

CGGCCCAGCCCACAGAGTTGGTGCCGATGTCCAGGCCGATGCTGTACTTCTTGTCGGCTGCTGGGACTCCGTGGATACCG

ACCTTCCGCTTCTTCTTTGGGGCCATCTTATCGTCATCGTCTTTGTAATCAATATCATGATCCTTGTAGTCTCCGTCGTG

GTCCTTATAGTCCATcctaggTGATATATTTCTATTAGGTATTTATTATTATAAAATATAAATCTTGAATGATAATAAAT

AAAATATTAGTTATTCCTTTTCTAGTTTAAAATATACATATTATAAATATATATATATATATATATATTTTTATTGTGAC

AAGAATATATAATTATAAATTATATTATTTATTTTTGTATTTTTTTTTTTTTTTTTTTTTTTTTCTTTTTTTGTTTTATT

TTTCTTTTTTTTTATAAATATTATTTTTTTCTTTTATCATGCACATTGGAATAATACATTAATATATATATATATATTAT

ATTATACATATATTGAATAATGTTTATAAAAAATGCATAACTTATATGAATATAATTTTTTTTAAATATGACAAAAAGAA

AAAAAAAAAAAACCAAAAAAAATTAAAATTGAAATGAAATATATAAATATATTATTTATATATATTATACATTGTTTAAT

ACTACTACATGTATATATATATATTATATATATATATATATATCAATTTTTTCAAAAATAAATTAATATAAAAAGAGGGG

AAAAAAAAAAAAAAAAAAAAAAAAAGATAATTAAGTAAGCATTTAAAAATATATAAATTGATAATATATAAAATTAATCA

CATATAAACTAATATAATTTATAAAATAAGGAAAATAAAATATTACCATAAAATAAAAATAAAAATAAAAAAAAAAAAAA

AAAACACCTTTTTTTATATATATTAATATATAATTATCTCTTAGAAAAAATATTGTATAATTATATATGTAATGATTTAT

ATAAAAAAATAAAATTATACAAGTATATATTTTGTTTCTATAAATTGATATCTTAATTATTTATTATTAGAAATAGATAT

TTTTATAATAAACCAATAGATAAAATTTGTAGAGAAAAAAAAATAAAAATAAAAATAAAAATAATATAATATATAATAAA

ATAAAATAATATTATATAAATATATTTTAATTTTTTTTACAAAATGGTTGGGGgcgccaccggtTATGTAATAATAATAT

GATCATAATATTATAATAAAACTTATAAAAAAAATATTAAATATTTCATAAATGATTATTATTTATAATAAGATAGATTC

TTTAATTATTTTAAAATTGTATATTTTTTATGTATATTAATTTATTAATATTAATAAGAATATTATAAAAATTCTATTTT

ATTATTTAATGATATTACCCTAATAAAAATAATATAATTATATTACAATATAAATTTATATATATATATATATTTATATA

TTTAAAGAATATTTTATTTTTCAATAAGAACCTTCATTTTAAATTAACATCAAATTATATATATGTATATATACTTCTTA

GTATTATTAATTAAAATACGGAATAATATATAATATATATAAAATGGCAAAACTTTCCTATAGAAAAAAATATTCCATTT

ATTATATTTGTTGTAGGTAATTCTTATTACCGTTTCCTTCTGTTCGTAATGTATATTGGTATGTACTTTATTTTTGCAAT

TTAATTATATGTAAAAAAACGTTAGTACACCATATATATATTATAGTTATAAGAATGCATGCCAAGCCTTTGTCTCAAGA

AGAATCCACCCTCATTGAAAGAGCAACGGCTACAATCAACAGCATCCCCATCTCTGAAGACTACAGCGTCGCCAGCGCAG

CTCTCTCTAGCGACGGCCGCATCTTCACTGGTGTCAATGTATATCATTTTACTGGGGGACCTTGTGCAGAACTCGTGGTG

CTGGGCACTGCTGCTGCTGCGGCAGCTGGCAACCTGACTTGTATCGTCGCGATCGGAAATGAGAACAGGGGCATCTTGAG

CCCCTGCGGACGGTGCCGACAGGTGCTTCTCGATCTGCATCCTGGGATCAAAGCCATAGTGAAGGACAGTGATGGACAGC

CGACGGCAGTTGGGATTCGTGAATTGCTGCCCTCTGGTTATGTGTGGGAGGGCgCTAGcGGCAGTGGAGAGGGCAGAGGA

AGTCTGCTAACATGCGGTGACGTCGAGGAGAATCCTGGCCCAaagcttATGGCACCATAAGGGTAGAAGTACAAATGAAA

TTGAACTTAGCGCAAGAGATGTTTTGGAAAATATTGGAATAGGAATATATAATCAGGAAAAAATAAAAAAGAATCCATAT

GAACAACAATTGAAAGGCACATTATCAAACGCCCGATTTCATGATGGCTTGCACAAGGCAGCTGATTTGGGGGTAATACC

TGGTCCTTCACATTTTTCTCAGCTTTATTACAAAAAGCATACTAATAACACAAAATATTATAAGGATGATAGGCATCCTT

GTCATGGTAGACAAGGAAAACGTTTTGATGAAGGTCAAAAATTTGAATGTGGTAATGATAAAATAATTGGTAATAGCGAT

AAATATGGATCCTGTGCTCCACCTAGAAGAAGACATATATGTGATCAAAATTTAGAATTCTTAGATAACAATCATACTGA

TACTATTCATGATGTATTGGGAAATGTGTTGGTCACAGCAAAATATGAAGGTGAATCTATTGTTAATGATCATCCAGATA

AAAAGAACAATGGTAATAAATCAGGTATATGTACTTCTCTTGCACGAAGTTTTGCCGATATAGGTGATATTGTAAGAGGA

AGAGATATGgtcgacaCTAGTACCGGTACGCGTgacgtCAGGTGGCACTTTTCGGGGAAATGTGCGCGGAACCCCTATTT

GTTTATTTTTCTAAATACATTCAAATATGTATCCGCTCATGAGACAATAACCCTGATAAATGCTTCAATAATATTGAAAA

AGGAAGAGTATGAGTATTCAACATTTCCGTGTCGCCCTTATTCCCTTTTTTGCGGCATTTTGCCTTCCTGTTTTTGCTCA

CCCAGAAACGCTGGTGAAAGTAAAAGATGCTGAAGATCAGTTGGGTGCACGAGTGGGTTACATCGAACTGGATCTCAACA

GCGGTAAGATCCTTGAGAGTTTTCGCCCCGAAGAACGTTTTCCAATGATGAGCACTTTTAAAGTTCTGCTATGTGGCGCG

GTATTATCCCGTATTGACGCCGGGCAAGAGCAACTCGGTCGCCGCATACACTATTCTCAGAATGACTTGGTTGAGTACTC

ACCAGTCACAGAAAAGCATCTTACGGATGGCATGACAGTAAGAGAATTATGCAGTGCTGCCATAACCATGAGTGATAACA

CTGCGGCCAACTTACTTCTGACAACGATCGGAGGACCGAAGGAGCTAACCGCTTTTTTGCACAACATGGGGGATCATGTA

ACTCGCCTTGATCGTTGGGAACCGGAGCTGAATGAAGCCATACCAAACGACGAGCGTGACACCACGATGCCTGTAGCAAT

GCCAACAACGTTGCGCAAACTATTAACTGGCGAACTACTTACTCTAGCTTCCCGGCAACAATTAATAGACTGGATGGAGG

CGGATAAAGTTGCAGGACCACTTCTGCGCTCGGCCCTTCCGGCTGGCTGGTTTATTGCTGATAAATCTGGAGCCGGTGAG

CGTGGGTCTCGCGGTATCATTGCAGCACTGGGGCCAGATGGTAAGCCCTCCCGTATCGTAGTTATCTACACGACGGGGAG

TCAGGCAACTATGGATGAACGAAATAGACAGATCGCTGAGATAGGTGCCTCACTGATTAAGCATTGGTAACTGTCAGACC

AAGTTTACTCATATATACTTTAGATTGATTTAAAACTTCATTTTTAATTTAAAAGGATCTAGGTGAAGATCCTTTTTGAT

AATCTCATGACCAAAATCCCTTAACGTGAGTTTTCGTTCCACTGAGCGTCAGACCCCGTAGAAAAGATCAAAGGATCTTC

TTGAGATCCTTTTTTTCTGCGCGTAATCTGCTGCTTGCAAACAAAAAAACCACCGCTACCAGCGGTGGTTTGTTTGCCGG

ATCAAGAGCTACCAACTCTTTTTCCGAAGGTAACTGGCTTCAGCAGAGCGCAGATACCAAATACTGTCCTTCTAGTGTAG

CCGTAGTTAGGCCACCACTTCAAGAACTCTGTAGCACCGCCTACATACCTCGCTCTGCTAATCCTGTTACCAGTGGCTGC

TGCCAGTGGCGATAAGTCGTGTCTTACCGGGTTGGACTCAAGACGATAGTTACCGGATAAGGCGCAGCGGTCGGGCTGAA

CGGGGGGTTCGTGCACACAGCCCAGCTTGGAGCGAACGACCTACACCGAACTGAGATACCTACAGCGTGAGCTATGAGAA

AGCGCCACGCTTCCCGAAGGGAGAAAGGCGGACAGGTATCCGGTAAGCGGCAGGGTCGGAACAGGAGAGCGCACGAGGGA

GCTTCCAGGGGGAAACGCCTGGTATCTTTATAGTCCTGTCGGGTTTCGCCACCTCTGACTTGAGCGTCGATTTTTGTGAT

GCTCGTCAGGGGGGCGGAGCCTATCGAAAAACGCCAGCAACGCGGCCTTTTTACGGTTCCTGGCCTTTTGCTGGCCTTTT

GCTCACATGTTCTTTCCTGCGTTATCCCCTGATTCTGTGGATAACCGTATTACCGCCTTTGAGTGAGCTGATACCGCTCG

CCGCAGCCGAACGACCGAGCGCAGCGAGTCAGTGAGCGAGGAAGCGGAAGAGCGCCCAATACGCAAACCGCCTCTCCCCG

CGCGTTGGCCGATTCATTAATGCAGCTGGCACGACAGGTTTCCCGACTGGAAAGCGGGCAGTGAGCGCAACGCAATTAAT

GTGAGTTAGCTCACTCATTAGGCACCCCAGGCTTTACACTTTATGCTTCCGGCTCGTATGTTGTGTGGAATTGTGAGCGG

ATAACAATTTCACACAGGAAACAGCTATGACCATGATTACGCCAAGCTATTTAGGTGACACTATAGAATACTC

Nucleotide sequence of plasmid Cas9_dualgRNA_it4var60prom_bsd_2A_exonI_ Mut A_Y73A_K263E

AAGCTTGGGGGGATCCGCCTTAAAAACTTCATTATATTTAAAAATTATTTTATAGGAAATAATAAAAAAAAAAgcaccga

ctcggtgccactttttcaagttgataacggactagccttattttaacttgctaTTTCtagctctaaaacTTTGGTGCCAT

TCTTATAACAATATTATATACTTAATATGAAATATGTGCATATAGGAAAAATTATGCATTTTGGTTACTCTAATATTATA

TATATATATATATATATATATATATTATAATATATTATGTTATATATACATAACATATACATTTTTTAATAATAATTTAC

CCTTTATTTTTACATTATAAAAAATTATATTACAGTAAAAATAAAAGTTTATTATATTAATAGTTTTTTTTTTTTTTTTT

TTAATTTATGAAATATTTAAATATTTAAAATTTTTTAAATGAATAATTATATTTATAATTAGAAAAAAAAAAAAAAAAAA

AAAAAAAAAATATAGCTATTTATATAAATTTCTTTTATTTATCTGAACAAGCAAGAATTTTTTTTTATATTAAATTAGAA

TAAATTATTATTAGTTTATGTATATATTTTTTTTTTTTCATAGTATATAAATATTATATATATTGTACCTTTTTACAATA

TATTTCATATATAGAAGAGAAAAAAAAAAAAAGAAGATATTATTGTAAAACCTCAAGATGTGTAGAAATCCAAATGTCGG

ATCCTCTAGAGTCGACCTGCAGGCATGCTATTTGATGAATTAACTACACTTAAAATAATACAATTATTATTAAATTTTTT

TTTGATTTATTTATTAATTTTTAAACTTAATCATTTGTATTTGGGAGGAATTATATATATCTTTATAATTATTTTATTTT

TTTTTATTTTTTTATTTTTTTATTATTATTATTTTTTTTTATTTTTTTTTTTTACTGTATCAAAGAAAAACCTTTAAAAA

AAAAATTATAATTTCCCCATCTTACTATATTTTTAATACATACGTTTTAAGGAATTAAATTAGACAAAAGCTATATTATG

CTTTACATATAATTAGAATTTATAAACGTTTGGTTATTAGATATTTCATGTCTCAGTAAAGTCTTTCAATACATATGTAA

AAAAATATATATGAATACACATAAGTTGTTAATATATTTTATATGCATAAATGTATAAATATATATATATATATATATAT

ATGTATGTATGTATATGTGTGTATATGAAATTATTTCAATGTTTAATTTTTTAAATTTTAATTTTTTTTTTTTTTTTTTT

TTTTATTATGTATATTGATCTTTATTATTTAAATATTACTTTTTTCGTTTTTTCTTCTTTTTATTATTTTTTTTTTTTTT

TATATTTTATACAAATGGTAATTCAAATAAAAGGTATAAATTTATATTTAATTTTCTTTTATGGATAAATAAAAGAAAAA

TATAAATATATAAAAATATAAAAATATATATATGTATATTGGGGTGATGATAAAATGAAAGATAATATATATATATATAT

ATCTTTATTTTTTTTTTTTTGTAGACCCCATTGTGAGTACATAAATATATTATATAACTCGAGTTACTTTTTCTTTTTTG

CCTGGCCGGCCTTTTTCGTGGCCGCCGGCCTTTTGTCGCCTCCCAGCTGAGACAGGTCGATCCGTGTCTCGTACAGGCCG

GTGATGCTCTGGTGGATCAGGGTGGCGTCCAGCACCTCTTTGGTGCTGGTGTACCTCTTCCGGTCGATGGTGGTGTCAAA

GTACTTGAAGGCGGCAGGGGCTCCCAGATTGGTCAGGGTAAACAGGTGGATGATATTCTCGGCCTGCTCTCTGATGGGCT

TATCCCGGTGCTTGTTGTAGGCGGACAGCACTTTGTCCAGATTAGCGTCGGCCAGGATCACTCTCTTGGAGAACTCGCTG

ATCTGCTCGATGATCTCGTCCAGGTAGTGCTTGTGCTGTTCCACAAACAGCTGTTTCTGCTCATTATCCTCGGGGGAGCC

CTTCAGCTTCTCATAGTGGCTGGCCAGGTACAGGAAGTTCACATATTTGGAGGGCAGGGCCAGTTCGTTTCCCTTCTGCA

GTTCGCCGGCAGAGGCCAGCATTCTCTTCCGGCCGTTTTCCAGCTCGAACAGGGAGTACTTAGGCAGCTTGATGATCAGG

TCCTTTTTCACTTCTTTGTAGCCCTTGGCTTCCAGAAAGTCGATGGGATTCTTCTCGAAGCTGCTTCTTTCCATGATGGT

GATCCCCAGCAGCTCTTTCACACTCTTCAGTTTCTTGGACTTGCCCTTTTCCACTTTGGCCACCACCAGCACAGAATAGG

CCACGGTGGGGCTGTCGAAGCCGCCGTACTTCTTAGGGTCCCAGTCCTTCTTTCTGGCGATCAGCTTATCGCTGTTCCTC

TTGGGCAGGATAGACTCTTTGCTGAAGCCGCCTGTCTGCACCTCGGTCTTTTTCACGATATTCACTTGGGGCATGCTCAG

CACTTTCCGCACGGTGGCAAAATCCCGGCCCTTATCCCACACGATCTCCCCGGTTTCGCCGTTTGTCTCGATCAGAGGCC

GCTTCCGGATCTCGCCGTTGGCCAGGGTAATCTCGGTCTTGAAAAAGTTCATGATGTTGCTGTAGAAGAAGTACTTGGCG

GTAGCCTTGCCGATTTCCTGCTCGCTCTTGGCGATCATCTTCCGCACGTCGTACACCTTGTAGTCGCCGTACACGAACTC

GCTTTCCAGCTTAGGGTACTTTTTGATCAGGGCGGTTCCCACGACGGCGTTCAGGTAGGCGTCGTGGGCGTGGTGGTAGT

TGTTGATCTCGCGCACTTTGTAAAACTGGAAATCCTTCCGGAAATCGGACACCAGCTTGGACTTCAGGGTGATCACTTTC

ACTTCCCGGATCAGCTTGTCATTCTCGTCGTACTTAGTGTTCATCCGGGAGTCCAGGATCTGTGCCACGTGCTTTGTGAT

CTGCCGGGTTTCCACCAGCTGTCTCTTGATGAAGCCGGCCTTATCCAGTTCGCTCAGGCCGCCTCTCTCGGCCTTGGTCA

GATTGTCGAACTTTCTCTGGGTAATCAGCTTGGCGTTCAGCAGCTGCCGCCAGTAGTTCTTCATCTTCTTCACGACCTCT

TCGGAGGGCACGTTGTCGCTCTTGCCCCGGTTCTTGTCGCTTCTGGTCAGCACCTTGTTGTCGATGGAGTCGTCCTTCAG

AAAGCTCTGAGGCACGATATGGTCCACATCGTAGTCGGACAGCCGGTTGATGTCCAGTTCCTGGTCCACGTACATATCCC

GCCCATTCTGCAGGTAGTACAGGTACAGCTTCTCGTTCTGCAGCTGGGTGTTTTCCACGGGGTGTTCTTTCAGGATCTGG

CTGCCCAGCTCTTTGATGCCCTCTTCGATCCGCTTCATTCTCTCGCGGCTGTTCTTCTGTCCCTTCTGGGTGGTCTGGTT

CTCTCTGGCCATTTCGATCACGATGTTCTCGGGCTTGTGCCGGCCCATCACTTTCACGAGCTCGTCCACCACCTTCACTG

TCTGCAGGATGCCCTTCTTAATGGCGGGGCTGCCGGCCAGATTGGCAATGTGCTCGTGCAGGCTATCGCCCTGGCCGGAC

ACCTGGGCTTTCTGGATGTCCTCTTTAAAGGTCAGGCTGTCGTCGTGGATCAGCTGCATGAAGTTTCTGTTGGCGAAGCC

GTCGGACTTCAGGAAATCCAGGATTGTCTTGCCGGACTGCTTGTCCCGGATGCCGTTGATCAGCTTCCGGCTCAGCCTGC

CCCAGCCGGTGTATCTCCGCCGCTTCAGCTGCTTCATCACTTTGTCGTCGAACAGGTGGGCATAGGTTTTCAGCCGTTCC

TCGATCATCTCTCTGTCCTCAAACAGTGTCAGGGTCAGCACGATATCTTCCAGAATGTCCTCGTTTTCCTCATTGTCCAG

GAAGTCCTTGTCCTTGATAATTTTCAGCAGATCGTGGTATGTGCCCAGGGAGGCGTTGAACCGATCTTCCACGCCGGAGA

TTTCCACGGAGTCGAAGCACTCGATTTTCTTGAAGTAGTCCTCTTTCAGCTGCTTCACGGTCACTTTCCGGTTGGTCTTG

AACAGCAGGTCCACGATGGCCTTTTTCTGCTCGCCGCTCAGGAAGGCGGGCTTTCTCATTCCCTCGGTCACGTATTTCAC

TTTGGTCAGCTCGTTATACACGGTGAAGTACTCGTACAGCAGGCTGTGCTTGGGCAGCACCTTCTCGTTGGGCAGGTTCT

TATCGAAGTTGGTCATCCGCTCGATGAAGCTCTGGGCGGAAGCGCCCTTGTCCACCACTTCCTCGAAGTTCCAGGGGGTG

ATGGTTTCCTCGCTCTTTCTGGTCATCCAGGCGAATCTGCTGTTTCCCCTGGCCAGAGGGCCCACGTAGTAGGGGATGCG

GAAGGTCAGGATCTTCTCGATCTTTTCCCGGTTGTCCTTCAGGAATGGGTAAAAATCTTCCTGCCGCCGCAGAATGGCGT

GCAGCTCTCCCAGGTGGATCTGGTGGGGGATGCTGCCGTTGTCGAAGGTCCGCTGCTTCCGCAGCAGGTCCTCTCTGTTC

AGCTTCACGAGCAGTTCCTCGGTGCCGTCCATCTTTTCCAGGATGGGCTTGATGAACTTGTAGAACTCTTCCTGGCTGGC

TCCGCCGTCAATGTAGCCGGCGTAGCCGTTCTTGCTCTGGTCGAAGAAAATCTCTTTGTACTTCTCAGGCAGCTGCTGCC

GCACGAGAGCTTTCAGCAGGGTCAGGTCCTGGTGGTGCTCGTCGTATCTCTTGATCATAGAGGCGCTCAGGGGGGCCTTG

GTGATCTCGGTGTTCACTCTCAGGATGTCGCTCAGCAGGATGGCGTCGGACAGGTTCTTGGCGGCCAGAAACAGGTCGGC

GTACTGGTCGCCGATCTGGGCCAGCAGGTTGTCCAGGTCGTCGTCGTAGGTGTCCTTGCTCAGCTGCAGTTTGGCATCCT

CGGCCAGGTCGAAGTTGCTCTTGAAGTTGGGGGTCAGGCCCAGGCTCAGGGCAATCAGGTTTCCGAACAGGCCATTCTTC

TTCTCGCCGGGCAGCTGGGCGATCAGATTTTCCAGCCGTCTGCTCTTGCTCAGTCTGGCAGACAGGATGGCCTTGGCGTC

CACGCCGCTGGCGTTGATGGGGTTTTCCTCGAACAGCTGGTTGTAGGTCTGCACCAGCTGGATGAACAGCTTGTCCACGT

CGCTGTTGTCGGGGTTCAGGTCGCCCTCGATCAGGAAGTGGCCCCGGAACTTGATCATGTGGGCCAGGGCCAGATAGATC

AGCCGCAGGTCGGCCTTGTCGGTGCTGTCCACCAGTTTCTTTCTCAGGTGGTAGATGGTGGGGTACTTCTCGTGGTAGGC

CACCTCGTCCACGATGTTGCCGAAGATGGGGTGCCGCTCGTGCTTCTTATCCTCTTCCACCAGGAAGGACTCTTCCAGTC

TGTGGAAGAAGCTGTCGTCCACCTTGGCCATCTCGTTGCTGAAGATCTCTTGCAGATAGCAGATCCGGTTCTTCCGTCTG

GTGTATCTTCTTCTGGCGGTTCTCTTCAGCCGGGTGGCCTCGGCTGTTTCGCCGCTGTCGAACAGCAGGGCTCCGATCAG

GTTCTTCTTGATGCTGTGCCGGTCGGTGTTGCCCAGCACCTTGAATTTCTTGCTGGGCACCTTGTACTCGTCGGTGATCA

CGGCCCAGCCCACAGAGTTGGTGCCGATGTCCAGGCCGATGCTGTACTTCTTGTCGGCTGCTGGGACTCCGTGGATACCG

ACCTTCCGCTTCTTCTTTGGGGCCATCTTATCGTCATCGTCTTTGTAATCAATATCATGATCCTTGTAGTCTCCGTCGTG

GTCCTTATAGTCCATcctaggTGATATATTTCTATTAGGTATTTATTATTATAAAATATAAATCTTGAATGATAATAAAT

AAAATATTAGTTATTCCTTTTCTAGTTTAAAATATACATATTATAAATATATATATATATATATATATTTTTATTGTGAC

AAGAATATATAATTATAAATTATATTATTTATTTTTGTATTTTTTTTTTTTTTTTTTTTTTTTTCTTTTTTTGTTTTATT

TTTCTTTTTTTTTATAAATATTATTTTTTTCTTTTATCATGCACATTGGAATAATACATTAATATATATATATATATTAT

ATTATACATATATTGAATAATGTTTATAAAAAATGCATAACTTATATGAATATAATTTTTTTTAAATATGACAAAAAGAA

AAAAAAAAAAAACCAAAAAAAATTAAAATTGAAATGAAATATATAAATATATTATTTATATATATTATACATTGTTTAAT

ACTACTACATGTATATATATATATTATATATATATATATATATCAATTTTTTCAAAAATAAATTAATATAAAAAGAGGGG

AAAAAAAAAAAAAAAAAAAAAAAAAGATAATTAAGTAAGCATTTAAAAATATATAAATTGATAATATATAAAATTAATCA

CATATAAACTAATATAATTTATAAAATAAGGAAAATAAAATATTACCATAAAATAAAAATAAAAATAAAAAAAAAAAAAA

AAAACACCTTTTTTTATATATATTAATATATAATTATCTCTTAGAAAAAATATTGTATAATTATATATGTAATGATTTAT

ATAAAAAAATAAAATTATACAAGTATATATTTTGTTTCTATAAATTGATATCTTAATTATTTATTATTAGAAATAGATAT

TTTTATAATAAACCAATAGATAAAATTTGTAGAGAAAAAAAAATAAAAATAAAAATAAAAATAATATAATATATAATAAA

ATAAAATAATATTATATAAATATATTTTAATTTTTTTTACAAAATGGTTGGGGgcgccaccggtTATGTAATAATAATAT

GATCATAATATTATAATAAAACTTATAAAAAAAATATTAAATATTTCATAAATGATTATTATTTATAATAAGATAGATTC

TTTAATTATTTTAAAATTGTATATTTTTTATGTATATTAATTTATTAATATTAATAAGAATATTATAAAAATTCTATTTT

ATTATTTAATGATATTACCCTAATAAAAATAATATAATTATATTACAATATAAATTTATATATATATATATATTTATATA

TTTAAAGAATATTTTATTTTTCAATAAGAACCTTCATTTTAAATTAACATCAAATTATATATATGTATATATACTTCTTA

GTATTATTAATTAAAATACGGAATAATATATAATATATATAAAATGGCAAAACTTTCCTATAGAAAAAAATATTCCATTT

ATTATATTTGTTGTAGGTAATTCTTATTACCGTTTCCTTCTGTTCGTAATGTATATTGGTATGTACTTTATTTTTGCAAT

TTAATTATATGTAAAAAAACGTTAGTACACCATATATATATTATAGTTATAAGAATGCATGCCAAGCCTTTGTCTCAAGA

AGAATCCACCCTCATTGAAAGAGCAACGGCTACAATCAACAGCATCCCCATCTCTGAAGACTACAGCGTCGCCAGCGCAG

CTCTCTCTAGCGACGGCCGCATCTTCACTGGTGTCAATGTATATCATTTTACTGGGGGACCTTGTGCAGAACTCGTGGTG

CTGGGCACTGCTGCTGCTGCGGCAGCTGGCAACCTGACTTGTATCGTCGCGATCGGAAATGAGAACAGGGGCATCTTGAG

CCCCTGCGGACGGTGCCGACAGGTGCTTCTCGATCTGCATCCTGGGATCAAAGCCATAGTGAAGGACAGTGATGGACAGC

CGACGGCAGTTGGGATTCGTGAATTGCTGCCCTCTGGTTATGTGTGGGAGGGCgCTAGcGGCAGTGGAGAGGGCAGAGGA

AGTCTGCTAACATGCGGTGACGTCGAGGAGAATCCTGGCCCAaagcttATGGCACCAAAGGGTAGAAGTACAAATGAAAT

TGAACTTAGCGCAAGAGATGTTTTGGAAAATATTGGAATAGGAATATATAATCAGGAAAAAATAAAAAAGAATCCATATG

AACAACAATTGAAAGGCACATTATCAAACGCCCGATTTCATGATGGCTTGCACAAGGCAGCTGATTTGGGGGTAATACCT

GGTCCTTCACATTTTTCTCAGCTTGCATACAAAAAGCATACTAATAACACAAAATATTATAAGGATGATAGGCATCCTTG

TCATGGTAGACAAGGAAAACGTTTTGATGAAGGTCAAAAATTTGAATGTGGTAATGATAAAATAATTGGTAATAGCGATA

AATATGGATCCTGTGCTCCACCTAGAAGAAGACATATATGTGATCAAAATTTAGAATTCTTAGATAACAATCATACTGAT

ACTATTCATGATGTATTGGGAAATGTGTTGGTCACAGCAAAATATGAAGGTGAATCTATTGTTAATGATCATCCAGATAA

AAAGAACAATGGTAATAAATCAGGTATATGTACTTCTCTTGCACGAAGTTTTGCCGATATAGGTGATATTGTAAGAGGAA

GAGATATGTTTAAACCTAATGACAAAGATGCAGTGCGGCATGGTTTAAAGGTAGTTTTTAAGAAAATATATGATAAATTG

TCACCTAAAGTACAAGAACATTACAAAGATGTTGATGGATCTGGAAATTACTATAAATTAAGGGAAGATTGGTGGACAGC

GAACAGAGATCAAGTATGGAAAGCCATAACATATGAAGCTCCGCAGGATGCTAATTATTTTAGAAATGTTTCAGGAACAA

CTATGGCGTTTACAAGTGCAGGAAAATGTAGACACAATGACAATAGCGTCCCAACGAATCTAGATTATGTCCCTCAATTT

TTACGTTGGTACGATGAATGGGCAGATGATTTTTGTCGAATAAGAAATCATAAGTTGCAAAAGGTTAAAGACACATGTAC

TAGTACCGGTACGCGTgacgtCAGGTGGCACTTTTCGGGGAAATGTGCGCGGAACCCCTATTTGTTTATTTTTCTAAATA

CATTCAAATATGTATCCGCTCATGAGACAATAACCCTGATAAATGCTTCAATAATATTGAAAAAGGAAGAGTATGAGTAT

TCAACATTTCCGTGTCGCCCTTATTCCCTTTTTTGCGGCATTTTGCCTTCCTGTTTTTGCTCACCCAGAAACGCTGGTGA

AAGTAAAAGATGCTGAAGATCAGTTGGGTGCACGAGTGGGTTACATCGAACTGGATCTCAACAGCGGTAAGATCCTTGAG

AGTTTTCGCCCCGAAGAACGTTTTCCAATGATGAGCACTTTTAAAGTTCTGCTATGTGGCGCGGTATTATCCCGTATTGA

CGCCGGGCAAGAGCAACTCGGTCGCCGCATACACTATTCTCAGAATGACTTGGTTGAGTACTCACCAGTCACAGAAAAGC

ATCTTACGGATGGCATGACAGTAAGAGAATTATGCAGTGCTGCCATAACCATGAGTGATAACACTGCGGCCAACTTACTT

CTGACAACGATCGGAGGACCGAAGGAGCTAACCGCTTTTTTGCACAACATGGGGGATCATGTAACTCGCCTTGATCGTTG

GGAACCGGAGCTGAATGAAGCCATACCAAACGACGAGCGTGACACCACGATGCCTGTAGCAATGCCAACAACGTTGCGCA

AACTATTAACTGGCGAACTACTTACTCTAGCTTCCCGGCAACAATTAATAGACTGGATGGAGGCGGATAAAGTTGCAGGA

CCACTTCTGCGCTCGGCCCTTCCGGCTGGCTGGTTTATTGCTGATAAATCTGGAGCCGGTGAGCGTGGGTCTCGCGGTAT

CATTGCAGCACTGGGGCCAGATGGTAAGCCCTCCCGTATCGTAGTTATCTACACGACGGGGAGTCAGGCAACTATGGATG

AACGAAATAGACAGATCGCTGAGATAGGTGCCTCACTGATTAAGCATTGGTAACTGTCAGACCAAGTTTACTCATATATA

CTTTAGATTGATTTAAAACTTCATTTTTAATTTAAAAGGATCTAGGTGAAGATCCTTTTTGATAATCTCATGACCAAAAT

CCCTTAACGTGAGTTTTCGTTCCACTGAGCGTCAGACCCCGTAGAAAAGATCAAAGGATCTTCTTGAGATCCTTTTTTTC

TGCGCGTAATCTGCTGCTTGCAAACAAAAAAACCACCGCTACCAGCGGTGGTTTGTTTGCCGGATCAAGAGCTACCAACT

CTTTTTCCGAAGGTAACTGGCTTCAGCAGAGCGCAGATACCAAATACTGTCCTTCTAGTGTAGCCGTAGTTAGGCCACCA

CTTCAAGAACTCTGTAGCACCGCCTACATACCTCGCTCTGCTAATCCTGTTACCAGTGGCTGCTGCCAGTGGCGATAAGT

CGTGTCTTACCGGGTTGGACTCAAGACGATAGTTACCGGATAAGGCGCAGCGGTCGGGCTGAACGGGGGGTTCGTGCACA

CAGCCCAGCTTGGAGCGAACGACCTACACCGAACTGAGATACCTACAGCGTGAGCTATGAGAAAGCGCCACGCTTCCCGA

AGGGAGAAAGGCGGACAGGTATCCGGTAAGCGGCAGGGTCGGAACAGGAGAGCGCACGAGGGAGCTTCCAGGGGGAAACG

CCTGGTATCTTTATAGTCCTGTCGGGTTTCGCCACCTCTGACTTGAGCGTCGATTTTTGTGATGCTCGTCAGGGGGGCGG

AGCCTATCGAAAAACGCCAGCAACGCGGCCTTTTTACGGTTCCTGGCCTTTTGCTGGCCTTTTGCTCACATGTTCTTTCC

TGCGTTATCCCCTGATTCTGTGGATAACCGTATTACCGCCTTTGAGTGAGCTGATACCGCTCGCCGCAGCCGAACGACCG

AGCGCAGCGAGTCAGTGAGCGAGGAAGCGGAAGAGCGCCCAATACGCAAACCGCCTCTCCCCGCGCGTTGGCCGATTCAT

TAATGCAGCTGGCACGACAGGTTTCCCGACTGGAAAGCGGGCAGTGAGCGCAACGCAATTAATGTGAGTTAGCTCACTCA

TTAGGCACCCCAGGCTTTACACTTTATGCTTCCGGCTCGTATGTTGTGTGGAATTGTGAGCGGATAACAATTTCACACAG

GAAACAGCTATGACCATGATTACGCCAAGCTATTTAGGTGACACTATAGAATACTC

Nucleotide sequence of plasmid Cas9_itvar60prom_bsd_2A_exon1_Mut F_K97A

AAGCTTGGGGGGATCCGCCTTAAAAACTTCATTATATTTAAAAATTATTTTATAGGAAATAATAAAAAAAAAAgcaccga

ctcggtgccactttttcaagttgataacggactagccttattttaacttgctaTTTCtagctctaaaacTTTGGTGCCAT

TCTTATAACAATATTATATACTTAATATGAAATATGTGCATATAGGAAAAATTATGCATTTTGGTTACTCTAATATTATA

TATATATATATATATATATATATATTATAATATATTATGTTATATATACATAACATATACATTTTTTAATAATAATTTAC

CCTTTATTTTTACATTATAAAAAATTATATTACAGTAAAAATAAAAGTTTATTATATTAATAGTTTTTTTTTTTTTTTTT

TTAATTTATGAAATATTTAAATATTTAAAATTTTTTAAATGAATAATTATATTTATAATTAGAAAAAAAAAAAAAAAAAA

AAAAAAAAAATATAGCTATTTATATAAATTTCTTTTATTTATCTGAACAAGCAAGAATTTTTTTTTATATTAAATTAGAA

TAAATTATTATTAGTTTATGTATATATTTTTTTTTTTTCATAGTATATAAATATTATATATATTGTACCTTTTTACAATA

TATTTCATATATAGAAGAGAAAAAAAAAAAAAGAAGATATTATTGTAAAACCTCAAGATGTGTAGAAATCCAAATGTCGG

ATCCTCTAGAGTCGACCTGCAGGCATGCTATTTGATGAATTAACTACACTTAAAATAATACAATTATTATTAAATTTTTT

TTTGATTTATTTATTAATTTTTAAACTTAATCATTTGTATTTGGGAGGAATTATATATATCTTTATAATTATTTTATTTT

TTTTTATTTTTTTATTTTTTTATTATTATTATTTTTTTTTATTTTTTTTTTTTACTGTATCAAAGAAAAACCTTTAAAAA

AAAAATTATAATTTCCCCATCTTACTATATTTTTAATACATACGTTTTAAGGAATTAAATTAGACAAAAGCTATATTATG

CTTTACATATAATTAGAATTTATAAACGTTTGGTTATTAGATATTTCATGTCTCAGTAAAGTCTTTCAATACATATGTAA

AAAAATATATATGAATACACATAAGTTGTTAATATATTTTATATGCATAAATGTATAAATATATATATATATATATATAT

ATGTATGTATGTATATGTGTGTATATGAAATTATTTCAATGTTTAATTTTTTAAATTTTAATTTTTTTTTTTTTTTTTTT

TTTTATTATGTATATTGATCTTTATTATTTAAATATTACTTTTTTCGTTTTTTCTTCTTTTTATTATTTTTTTTTTTTTT

TATATTTTATACAAATGGTAATTCAAATAAAAGGTATAAATTTATATTTAATTTTCTTTTATGGATAAATAAAAGAAAAA

TATAAATATATAAAAATATAAAAATATATATATGTATATTGGGGTGATGATAAAATGAAAGATAATATATATATATATAT

ATCTTTATTTTTTTTTTTTTGTAGACCCCATTGTGAGTACATAAATATATTATATAACTCGAGTTACTTTTTCTTTTTTG

CCTGGCCGGCCTTTTTCGTGGCCGCCGGCCTTTTGTCGCCTCCCAGCTGAGACAGGTCGATCCGTGTCTCGTACAGGCCG

GTGATGCTCTGGTGGATCAGGGTGGCGTCCAGCACCTCTTTGGTGCTGGTGTACCTCTTCCGGTCGATGGTGGTGTCAAA

GTACTTGAAGGCGGCAGGGGCTCCCAGATTGGTCAGGGTAAACAGGTGGATGATATTCTCGGCCTGCTCTCTGATGGGCT

TATCCCGGTGCTTGTTGTAGGCGGACAGCACTTTGTCCAGATTAGCGTCGGCCAGGATCACTCTCTTGGAGAACTCGCTG

ATCTGCTCGATGATCTCGTCCAGGTAGTGCTTGTGCTGTTCCACAAACAGCTGTTTCTGCTCATTATCCTCGGGGGAGCC

CTTCAGCTTCTCATAGTGGCTGGCCAGGTACAGGAAGTTCACATATTTGGAGGGCAGGGCCAGTTCGTTTCCCTTCTGCA

GTTCGCCGGCAGAGGCCAGCATTCTCTTCCGGCCGTTTTCCAGCTCGAACAGGGAGTACTTAGGCAGCTTGATGATCAGG

TCCTTTTTCACTTCTTTGTAGCCCTTGGCTTCCAGAAAGTCGATGGGATTCTTCTCGAAGCTGCTTCTTTCCATGATGGT

GATCCCCAGCAGCTCTTTCACACTCTTCAGTTTCTTGGACTTGCCCTTTTCCACTTTGGCCACCACCAGCACAGAATAGG

CCACGGTGGGGCTGTCGAAGCCGCCGTACTTCTTAGGGTCCCAGTCCTTCTTTCTGGCGATCAGCTTATCGCTGTTCCTC

TTGGGCAGGATAGACTCTTTGCTGAAGCCGCCTGTCTGCACCTCGGTCTTTTTCACGATATTCACTTGGGGCATGCTCAG

CACTTTCCGCACGGTGGCAAAATCCCGGCCCTTATCCCACACGATCTCCCCGGTTTCGCCGTTTGTCTCGATCAGAGGCC

GCTTCCGGATCTCGCCGTTGGCCAGGGTAATCTCGGTCTTGAAAAAGTTCATGATGTTGCTGTAGAAGAAGTACTTGGCG

GTAGCCTTGCCGATTTCCTGCTCGCTCTTGGCGATCATCTTCCGCACGTCGTACACCTTGTAGTCGCCGTACACGAACTC

GCTTTCCAGCTTAGGGTACTTTTTGATCAGGGCGGTTCCCACGACGGCGTTCAGGTAGGCGTCGTGGGCGTGGTGGTAGT

TGTTGATCTCGCGCACTTTGTAAAACTGGAAATCCTTCCGGAAATCGGACACCAGCTTGGACTTCAGGGTGATCACTTTC

ACTTCCCGGATCAGCTTGTCATTCTCGTCGTACTTAGTGTTCATCCGGGAGTCCAGGATCTGTGCCACGTGCTTTGTGAT

CTGCCGGGTTTCCACCAGCTGTCTCTTGATGAAGCCGGCCTTATCCAGTTCGCTCAGGCCGCCTCTCTCGGCCTTGGTCA

GATTGTCGAACTTTCTCTGGGTAATCAGCTTGGCGTTCAGCAGCTGCCGCCAGTAGTTCTTCATCTTCTTCACGACCTCT

TCGGAGGGCACGTTGTCGCTCTTGCCCCGGTTCTTGTCGCTTCTGGTCAGCACCTTGTTGTCGATGGAGTCGTCCTTCAG

AAAGCTCTGAGGCACGATATGGTCCACATCGTAGTCGGACAGCCGGTTGATGTCCAGTTCCTGGTCCACGTACATATCCC

GCCCATTCTGCAGGTAGTACAGGTACAGCTTCTCGTTCTGCAGCTGGGTGTTTTCCACGGGGTGTTCTTTCAGGATCTGG

CTGCCCAGCTCTTTGATGCCCTCTTCGATCCGCTTCATTCTCTCGCGGCTGTTCTTCTGTCCCTTCTGGGTGGTCTGGTT

CTCTCTGGCCATTTCGATCACGATGTTCTCGGGCTTGTGCCGGCCCATCACTTTCACGAGCTCGTCCACCACCTTCACTG

TCTGCAGGATGCCCTTCTTAATGGCGGGGCTGCCGGCCAGATTGGCAATGTGCTCGTGCAGGCTATCGCCCTGGCCGGAC

ACCTGGGCTTTCTGGATGTCCTCTTTAAAGGTCAGGCTGTCGTCGTGGATCAGCTGCATGAAGTTTCTGTTGGCGAAGCC

GTCGGACTTCAGGAAATCCAGGATTGTCTTGCCGGACTGCTTGTCCCGGATGCCGTTGATCAGCTTCCGGCTCAGCCTGC

CCCAGCCGGTGTATCTCCGCCGCTTCAGCTGCTTCATCACTTTGTCGTCGAACAGGTGGGCATAGGTTTTCAGCCGTTCC

TCGATCATCTCTCTGTCCTCAAACAGTGTCAGGGTCAGCACGATATCTTCCAGAATGTCCTCGTTTTCCTCATTGTCCAG

GAAGTCCTTGTCCTTGATAATTTTCAGCAGATCGTGGTATGTGCCCAGGGAGGCGTTGAACCGATCTTCCACGCCGGAGA

TTTCCACGGAGTCGAAGCACTCGATTTTCTTGAAGTAGTCCTCTTTCAGCTGCTTCACGGTCACTTTCCGGTTGGTCTTG

AACAGCAGGTCCACGATGGCCTTTTTCTGCTCGCCGCTCAGGAAGGCGGGCTTTCTCATTCCCTCGGTCACGTATTTCAC

TTTGGTCAGCTCGTTATACACGGTGAAGTACTCGTACAGCAGGCTGTGCTTGGGCAGCACCTTCTCGTTGGGCAGGTTCT

TATCGAAGTTGGTCATCCGCTCGATGAAGCTCTGGGCGGAAGCGCCCTTGTCCACCACTTCCTCGAAGTTCCAGGGGGTG

ATGGTTTCCTCGCTCTTTCTGGTCATCCAGGCGAATCTGCTGTTTCCCCTGGCCAGAGGGCCCACGTAGTAGGGGATGCG

GAAGGTCAGGATCTTCTCGATCTTTTCCCGGTTGTCCTTCAGGAATGGGTAAAAATCTTCCTGCCGCCGCAGAATGGCGT

GCAGCTCTCCCAGGTGGATCTGGTGGGGGATGCTGCCGTTGTCGAAGGTCCGCTGCTTCCGCAGCAGGTCCTCTCTGTTC

AGCTTCACGAGCAGTTCCTCGGTGCCGTCCATCTTTTCCAGGATGGGCTTGATGAACTTGTAGAACTCTTCCTGGCTGGC

TCCGCCGTCAATGTAGCCGGCGTAGCCGTTCTTGCTCTGGTCGAAGAAAATCTCTTTGTACTTCTCAGGCAGCTGCTGCC

GCACGAGAGCTTTCAGCAGGGTCAGGTCCTGGTGGTGCTCGTCGTATCTCTTGATCATAGAGGCGCTCAGGGGGGCCTTG

GTGATCTCGGTGTTCACTCTCAGGATGTCGCTCAGCAGGATGGCGTCGGACAGGTTCTTGGCGGCCAGAAACAGGTCGGC

GTACTGGTCGCCGATCTGGGCCAGCAGGTTGTCCAGGTCGTCGTCGTAGGTGTCCTTGCTCAGCTGCAGTTTGGCATCCT

CGGCCAGGTCGAAGTTGCTCTTGAAGTTGGGGGTCAGGCCCAGGCTCAGGGCAATCAGGTTTCCGAACAGGCCATTCTTC

TTCTCGCCGGGCAGCTGGGCGATCAGATTTTCCAGCCGTCTGCTCTTGCTCAGTCTGGCAGACAGGATGGCCTTGGCGTC

CACGCCGCTGGCGTTGATGGGGTTTTCCTCGAACAGCTGGTTGTAGGTCTGCACCAGCTGGATGAACAGCTTGTCCACGT

CGCTGTTGTCGGGGTTCAGGTCGCCCTCGATCAGGAAGTGGCCCCGGAACTTGATCATGTGGGCCAGGGCCAGATAGATC

AGCCGCAGGTCGGCCTTGTCGGTGCTGTCCACCAGTTTCTTTCTCAGGTGGTAGATGGTGGGGTACTTCTCGTGGTAGGC

CACCTCGTCCACGATGTTGCCGAAGATGGGGTGCCGCTCGTGCTTCTTATCCTCTTCCACCAGGAAGGACTCTTCCAGTC

TGTGGAAGAAGCTGTCGTCCACCTTGGCCATCTCGTTGCTGAAGATCTCTTGCAGATAGCAGATCCGGTTCTTCCGTCTG

GTGTATCTTCTTCTGGCGGTTCTCTTCAGCCGGGTGGCCTCGGCTGTTTCGCCGCTGTCGAACAGCAGGGCTCCGATCAG

GTTCTTCTTGATGCTGTGCCGGTCGGTGTTGCCCAGCACCTTGAATTTCTTGCTGGGCACCTTGTACTCGTCGGTGATCA

CGGCCCAGCCCACAGAGTTGGTGCCGATGTCCAGGCCGATGCTGTACTTCTTGTCGGCTGCTGGGACTCCGTGGATACCG

ACCTTCCGCTTCTTCTTTGGGGCCATCTTATCGTCATCGTCTTTGTAATCAATATCATGATCCTTGTAGTCTCCGTCGTG

GTCCTTATAGTCCATcctaggTGATATATTTCTATTAGGTATTTATTATTATAAAATATAAATCTTGAATGATAATAAAT

AAAATATTAGTTATTCCTTTTCTAGTTTAAAATATACATATTATAAATATATATATATATATATATATTTTTATTGTGAC

AAGAATATATAATTATAAATTATATTATTTATTTTTGTATTTTTTTTTTTTTTTTTTTTTTTTTCTTTTTTTGTTTTATT

TTTCTTTTTTTTTATAAATATTATTTTTTTCTTTTATCATGCACATTGGAATAATACATTAATATATATATATATATTAT

ATTATACATATATTGAATAATGTTTATAAAAAATGCATAACTTATATGAATATAATTTTTTTTAAATATGACAAAAAGAA

AAAAAAAAAAAACCAAAAAAAATTAAAATTGAAATGAAATATATAAATATATTATTTATATATATTATACATTGTTTAAT

ACTACTACATGTATATATATATATTATATATATATATATATATCAATTTTTTCAAAAATAAATTAATATAAAAAGAGGGG

AAAAAAAAAAAAAAAAAAAAAAAAAGATAATTAAGTAAGCATTTAAAAATATATAAATTGATAATATATAAAATTAATCA

CATATAAACTAATATAATTTATAAAATAAGGAAAATAAAATATTACCATAAAATAAAAATAAAAATAAAAAAAAAAAAAA

AAAACACCTTTTTTTATATATATTAATATATAATTATCTCTTAGAAAAAATATTGTATAATTATATATGTAATGATTTAT

ATAAAAAAATAAAATTATACAAGTATATATTTTGTTTCTATAAATTGATATCTTAATTATTTATTATTAGAAATAGATAT

TTTTATAATAAACCAATAGATAAAATTTGTAGAGAAAAAAAAATAAAAATAAAAATAAAAATAATATAATATATAATAAA

ATAAAATAATATTATATAAATATATTTTAATTTTTTTTACAAAATGGTTGGGGgcgccaccggtTATGTAATAATAATAT

GATCATAATATTATAATAAAACTTATAAAAAAAATATTAAATATTTCATAAATGATTATTATTTATAATAAGATAGATTC

TTTAATTATTTTAAAATTGTATATTTTTTATGTATATTAATTTATTAATATTAATAAGAATATTATAAAAATTCTATTTT

ATTATTTAATGATATTACCCTAATAAAAATAATATAATTATATTACAATATAAATTTATATATATATATATATTTATATA

TTTAAAGAATATTTTATTTTTCAATAAGAACCTTCATTTTAAATTAACATCAAATTATATATATGTATATATACTTCTTA

GTATTATTAATTAAAATACGGAATAATATATAATATATATAAAATGGCAAAACTTTCCTATAGAAAAAAATATTCCATTT

ATTATATTTGTTGTAGGTAATTCTTATTACCGTTTCCTTCTGTTCGTAATGTATATTGGTATGTACTTTATTTTTGCAAT

TTAATTATATGTAAAAAAACGTTAGTACACCATATATATATTATAGTTATAAGAATGCATGCCAAGCCTTTGTCTCAAGA

AGAATCCACCCTCATTGAAAGAGCAACGGCTACAATCAACAGCATCCCCATCTCTGAAGACTACAGCGTCGCCAGCGCAG

CTCTCTCTAGCGACGGCCGCATCTTCACTGGTGTCAATGTATATCATTTTACTGGGGGACCTTGTGCAGAACTCGTGGTG

CTGGGCACTGCTGCTGCTGCGGCAGCTGGCAACCTGACTTGTATCGTCGCGATCGGAAATGAGAACAGGGGCATCTTGAG

CCCCTGCGGACGGTGCCGACAGGTGCTTCTCGATCTGCATCCTGGGATCAAAGCCATAGTGAAGGACAGTGATGGACAGC

CGACGGCAGTTGGGATTCGTGAATTGCTGCCCTCTGGTTATGTGTGGGAGGGCgCTAGcGGCAGTGGAGAGGGCAGAGGA

AGTCTGCTAACATGCGGTGACGTCGAGGAGAATCCTGGCCCAaagcttATGGCACCAAAGGGTAGAAGTACAAATGAAAT

TGAACTTAGCGCAAGAGATGTTTTGGAAAATATTGGAATAGGAATATATAATCAGGAAAAAATAAAAAAGAATCCATATG

AACAACAATTGAAAGGCACATTATCAAACGCCCGATTTCATGATGGCTTGCACAAGGCAGCTGATTTGGGGGTAATACCT

GGTCCTTCACATTTTTCTCAGCTTTATTACAAAAAGCATACTAATAACACAAAATATTATAAGGATGATAGGCATCCTTG

TCATGGTAGACAAGGAGCACGTTTTGATGAAGGTCAAAAATTTGAATGTGGTAATGATAAAATAATTGGTAATAGCGATA

AATATGGATCCTGTGCTCCACCTAGAAGAAGACATATATGTGATCAAAATTTAGAATTCTTAGATAACAATCATACTGAT

ACTATTCATGATGTATTGGGAAATGTGTTGGTCACAGCAAAATATGAAGGTGAATCTATTGTTAATGATCATCCAGATAA

AAAGAACAATGGTAATAAATCAGGTATATGTACTTCTCTTGCACGAAGTTTTGCCGATATAGGTGATATTGTAAGAGGAA

GAGATATGgtcgacaCTAGTACCGGTACGCGTgacgtCAGGTGGCACTTTTCGGGGAAATGTGCGCGGAACCCCTATTTG

TTTATTTTTCTAAATACATTCAAATATGTATCCGCTCATGAGACAATAACCCTGATAAATGCTTCAATAATATTGAAAAA

GGAAGAGTATGAGTATTCAACATTTCCGTGTCGCCCTTATTCCCTTTTTTGCGGCATTTTGCCTTCCTGTTTTTGCTCAC

CCAGAAACGCTGGTGAAAGTAAAAGATGCTGAAGATCAGTTGGGTGCACGAGTGGGTTACATCGAACTGGATCTCAACAG

CGGTAAGATCCTTGAGAGTTTTCGCCCCGAAGAACGTTTTCCAATGATGAGCACTTTTAAAGTTCTGCTATGTGGCGCGG

TATTATCCCGTATTGACGCCGGGCAAGAGCAACTCGGTCGCCGCATACACTATTCTCAGAATGACTTGGTTGAGTACTCA

CCAGTCACAGAAAAGCATCTTACGGATGGCATGACAGTAAGAGAATTATGCAGTGCTGCCATAACCATGAGTGATAACAC

TGCGGCCAACTTACTTCTGACAACGATCGGAGGACCGAAGGAGCTAACCGCTTTTTTGCACAACATGGGGGATCATGTAA

CTCGCCTTGATCGTTGGGAACCGGAGCTGAATGAAGCCATACCAAACGACGAGCGTGACACCACGATGCCTGTAGCAATG

CCAACAACGTTGCGCAAACTATTAACTGGCGAACTACTTACTCTAGCTTCCCGGCAACAATTAATAGACTGGATGGAGGC

GGATAAAGTTGCAGGACCACTTCTGCGCTCGGCCCTTCCGGCTGGCTGGTTTATTGCTGATAAATCTGGAGCCGGTGAGC

GTGGGTCTCGCGGTATCATTGCAGCACTGGGGCCAGATGGTAAGCCCTCCCGTATCGTAGTTATCTACACGACGGGGAGT

CAGGCAACTATGGATGAACGAAATAGACAGATCGCTGAGATAGGTGCCTCACTGATTAAGCATTGGTAACTGTCAGACCA

AGTTTACTCATATATACTTTAGATTGATTTAAAACTTCATTTTTAATTTAAAAGGATCTAGGTGAAGATCCTTTTTGATA

ATCTCATGACCAAAATCCCTTAACGTGAGTTTTCGTTCCACTGAGCGTCAGACCCCGTAGAAAAGATCAAAGGATCTTCT

TGAGATCCTTTTTTTCTGCGCGTAATCTGCTGCTTGCAAACAAAAAAACCACCGCTACCAGCGGTGGTTTGTTTGCCGGA

TCAAGAGCTACCAACTCTTTTTCCGAAGGTAACTGGCTTCAGCAGAGCGCAGATACCAAATACTGTCCTTCTAGTGTAGC

CGTAGTTAGGCCACCACTTCAAGAACTCTGTAGCACCGCCTACATACCTCGCTCTGCTAATCCTGTTACCAGTGGCTGCT

GCCAGTGGCGATAAGTCGTGTCTTACCGGGTTGGACTCAAGACGATAGTTACCGGATAAGGCGCAGCGGTCGGGCTGAAC

GGGGGGTTCGTGCACACAGCCCAGCTTGGAGCGAACGACCTACACCGAACTGAGATACCTACAGCGTGAGCTATGAGAAA

GCGCCACGCTTCCCGAAGGGAGAAAGGCGGACAGGTATCCGGTAAGCGGCAGGGTCGGAACAGGAGAGCGCACGAGGGAG

CTTCCAGGGGGAAACGCCTGGTATCTTTATAGTCCTGTCGGGTTTCGCCACCTCTGACTTGAGCGTCGATTTTTGTGATG

CTCGTCAGGGGGGCGGAGCCTATCGAAAAACGCCAGCAACGCGGCCTTTTTACGGTTCCTGGCCTTTTGCTGGCCTTTTG

CTCACATGTTCTTTCCTGCGTTATCCCCTGATTCTGTGGATAACCGTATTACCGCCTTTGAGTGAGCTGATACCGCTCGC

CGCAGCCGAACGACCGAGCGCAGCGAGTCAGTGAGCGAGGAAGCGGAAGAGCGCCCAATACGCAAACCGCCTCTCCCCGC

GCGTTGGCCGATTCATTAATGCAGCTGGCACGACAGGTTTCCCGACTGGAAAGCGGGCAGTGAGCGCAACGCAATTAATG

TGAGTTAGCTCACTCATTAGGCACCCCAGGCTTTACACTTTATGCTTCCGGCTCGTATGTTGTGTGGAATTGTGAGCGGA

TAACAATTTCACACAGGAAACAGCTATGACCATGATTACGCCAAGCTATTTAGGTGACACTATAGAATACTC

Nucleotide sequence of plasmid Cas9_dualgRNA3_it4var60prom_bsd-2A-exonI_Mut G_K263E

AAGCTTGGGGGGATCCGCCTTAAAAACTTCATTATATTTAAAAATTATTTTATAGGAAATAATAAAAAAAAAAGCACCGA

CTCGGTGCCACTTTTTCAAGTTGATAACGGACTAGCCTTATTTTAACTTGCTATTTCTAGCTCTAAAACTTTGGTGCCAT

TCTTATAACAATATTATATACTTAATATGAAATATGTGCATATAGGAAAAATTATGCATTTTGGTTACTCTAATATTATA

TATATATATATATATATATATATATTATAATATATTATGTTATATATACATAACATATACATTTTTTAATAATAATTTAC

CCTTTATTTTTACATTATAAAAAATTATATTACAGTAAAAATAAAAGTTTATTATATTAATAGTTTTTTTTTTTTTTTTT

TTAATTTATGAAATATTTAAATATTTAAAATTTTTTAAATGAATAATTATATTTATAATTAGAAAAAAAAAAAAAAAAAA

AAAAAAAAAATATAGCTATTTATATAAATTTCTTTTATTTATCTGAACAAGCAAGAATTTTTTTTTATATTAAATTAGAA

TAAATTATTATTAGTTTATGTATATATTTTTTTTTTTTCATAGTATATAAATATTATATATATTGTACCTTTTTACAATA

TATTTCATATATAGAAGAGAAAAAAAAAAAAAGAAGATATTATTGTAAAACCTCAAGATGTGTAGAAATCCAAATGTCGG

ATCCTCTAGACGCCTTAAAAACTTCATTATATTTAAAAATTATTTTATAGGAAATAATAAAAAAAAAAGCACCGACTCGG

TGCCACTTTTTCAAGTTGATAACGGACTAGCCTTATTTTAACTTGCTATTTCTAGCTCTAAAACCAAGACGCTAATTATT

TTAGAATATTATATACTTAATATGAAATATGTGCATATAGGAAAAATTATGCATTTTGGTTACTCTAATATTATATATAT

ATATATATATATATATATATTATAATATATTATGTTATATATACATAACATATACATTTTTTAATAATAATTTACCCTTT

ATTTTTACATTATAAAAAATTATATTACAGTAAAAATAAAAGTTTATTATATTAATAGTTTTTTTTTTTTTTTTTTTAAT

TTATGAAATATTTAAATATTTAAAATTTTTTAAATGAATAATTATATTTATAATTAGAAAAAAAAAAAAAAAAAAAAAAA

AAAAATATAGCTATTTATATAAATTTCTTTTATTTATCTGAACAAGCAAGAATTTTTTTTTATATTAAATTAGAATAAAT

TATTATTAGTTTATGTATATATTTTTTTTTTTTCATAGTATATAAATATTATATATATTGTACCTTTTTACAATATATTT

CATATATAGAAGAGAAAAAAAAAAAAAGAAGATATTATTGTAAAACCTCAAGATGTGTAGAAATCCAAATGTCGCCTGCA

GGCATGCTATTTGATGAATTAACTACACTTAAAATAATACAATTATTATTAAATTTTTTTTTGATTTATTTATTAATTTT

TAAACTTAATCATTTGTATTTGGGAGGAATTATATATATCTTTATAATTATTTTATTTTTTTTTATTTTTTTATTTTTTT

ATTATTATTATTTTTTTTTATTTTTTTTTTTTACTGTATCAAAGAAAAACCTTTAAAAAAAAAATTATAATTTCCCCATC

TTACTATATTTTTAATACATACGTTTTAAGGAATTAAATTAGACAAAAGCTATATTATGCTTTACATATAATTAGAATTT

ATAAACGTTTGGTTATTAGATATTTCATGTCTCAGTAAAGTCTTTCAATACATATGTAAAAAAATATATATGAATACACA

TAAGTTGTTAATATATTTTATATGCATAAATGTATAAATATATATATATATATATATATATGTATGTATGTATATGTGTG

TATATGAAATTATTTCAATGTTTAATTTTTTAAATTTTAATTTTTTTTTTTTTTTTTTTTTTTATTATGTATATTGATCT

TTATTATTTAAATATTACTTTTTTCGTTTTTTCTTCTTTTTATTATTTTTTTTTTTTTTTATATTTTATACAAATGGTAA

TTCAAATAAAAGGTATAAATTTATATTTAATTTTCTTTTATGGATAAATAAAAGAAAAATATAAATATATAAAAATATAA

AAATATATATATGTATATTGGGGTGATGATAAAATGAAAGATAATATATATATATATATATCTTTATTTTTTTTTTTTTG

TAGACCCCATTGTGAGTACATAAATATATTATATAACTCGAGTTACTTTTTCTTTTTTGCCTGGCCGGCCTTTTTCGTGG

CCGCCGGCCTTTTGTCGCCTCCCAGCTGAGACAGGTCGATCCGTGTCTCGTACAGGCCGGTGATGCTCTGGTGGATCAGG

GTGGCGTCCAGCACCTCTTTGGTGCTGGTGTACCTCTTCCGGTCGATGGTGGTGTCAAAGTACTTGAAGGCGGCAGGGGC

TCCCAGATTGGTCAGGGTAAACAGGTGGATGATATTCTCGGCCTGCTCTCTGATGGGCTTATCCCGGTGCTTGTTGTAGG

CGGACAGCACTTTGTCCAGATTAGCGTCGGCCAGGATCACTCTCTTGGAGAACTCGCTGATCTGCTCGATGATCTCGTCC

AGGTAGTGCTTGTGCTGTTCCACAAACAGCTGTTTCTGCTCATTATCCTCGGGGGAGCCCTTCAGCTTCTCATAGTGGCT

GGCCAGGTACAGGAAGTTCACATATTTGGAGGGCAGGGCCAGTTCGTTTCCCTTCTGCAGTTCGCCGGCAGAGGCCAGCA

TTCTCTTCCGGCCGTTTTCCAGCTCGAACAGGGAGTACTTAGGCAGCTTGATGATCAGGTCCTTTTTCACTTCTTTGTAG

CCCTTGGCTTCCAGAAAGTCGATGGGATTCTTCTCGAAGCTGCTTCTTTCCATGATGGTGATCCCCAGCAGCTCTTTCAC

ACTCTTCAGTTTCTTGGACTTGCCCTTTTCCACTTTGGCCACCACCAGCACAGAATAGGCCACGGTGGGGCTGTCGAAGC

CGCCGTACTTCTTAGGGTCCCAGTCCTTCTTTCTGGCGATCAGCTTATCGCTGTTCCTCTTGGGCAGGATAGACTCTTTG

CTGAAGCCGCCTGTCTGCACCTCGGTCTTTTTCACGATATTCACTTGGGGCATGCTCAGCACTTTCCGCACGGTGGCAAA

ATCCCGGCCCTTATCCCACACGATCTCCCCGGTTTCGCCGTTTGTCTCGATCAGAGGCCGCTTCCGGATCTCGCCGTTGG

CCAGGGTAATCTCGGTCTTGAAAAAGTTCATGATGTTGCTGTAGAAGAAGTACTTGGCGGTAGCCTTGCCGATTTCCTGC

TCGCTCTTGGCGATCATCTTCCGCACGTCGTACACCTTGTAGTCGCCGTACACGAACTCGCTTTCCAGCTTAGGGTACTT

TTTGATCAGGGCGGTTCCCACGACGGCGTTCAGGTAGGCGTCGTGGGCGTGGTGGTAGTTGTTGATCTCGCGCACTTTGT

AAAACTGGAAATCCTTCCGGAAATCGGACACCAGCTTGGACTTCAGGGTGATCACTTTCACTTCCCGGATCAGCTTGTCA

TTCTCGTCGTACTTAGTGTTCATCCGGGAGTCCAGGATCTGTGCCACGTGCTTTGTGATCTGCCGGGTTTCCACCAGCTG

TCTCTTGATGAAGCCGGCCTTATCCAGTTCGCTCAGGCCGCCTCTCTCGGCCTTGGTCAGATTGTCGAACTTTCTCTGGG

TAATCAGCTTGGCGTTCAGCAGCTGCCGCCAGTAGTTCTTCATCTTCTTCACGACCTCTTCGGAGGGCACGTTGTCGCTC

TTGCCCCGGTTCTTGTCGCTTCTGGTCAGCACCTTGTTGTCGATGGAGTCGTCCTTCAGAAAGCTCTGAGGCACGATATG

GTCCACATCGTAGTCGGACAGCCGGTTGATGTCCAGTTCCTGGTCCACGTACATATCCCGCCCATTCTGCAGGTAGTACA

GGTACAGCTTCTCGTTCTGCAGCTGGGTGTTTTCCACGGGGTGTTCTTTCAGGATCTGGCTGCCCAGCTCTTTGATGCCC

TCTTCGATCCGCTTCATTCTCTCGCGGCTGTTCTTCTGTCCCTTCTGGGTGGTCTGGTTCTCTCTGGCCATTTCGATCAC

GATGTTCTCGGGCTTGTGCCGGCCCATCACTTTCACGAGCTCGTCCACCACCTTCACTGTCTGCAGGATGCCCTTCTTAA

TGGCGGGGCTGCCGGCCAGATTGGCAATGTGCTCGTGCAGGCTATCGCCCTGGCCGGACACCTGGGCTTTCTGGATGTCC

TCTTTAAAGGTCAGGCTGTCGTCGTGGATCAGCTGCATGAAGTTTCTGTTGGCGAAGCCGTCGGACTTCAGGAAATCCAG

GATTGTCTTGCCGGACTGCTTGTCCCGGATGCCGTTGATCAGCTTCCGGCTCAGCCTGCCCCAGCCGGTGTATCTCCGCC

GCTTCAGCTGCTTCATCACTTTGTCGTCGAACAGGTGGGCATAGGTTTTCAGCCGTTCCTCGATCATCTCTCTGTCCTCA

AACAGTGTCAGGGTCAGCACGATATCTTCCAGAATGTCCTCGTTTTCCTCATTGTCCAGGAAGTCCTTGTCCTTGATAAT

TTTCAGCAGATCGTGGTATGTGCCCAGGGAGGCGTTGAACCGATCTTCCACGCCGGAGATTTCCACGGAGTCGAAGCACT

CGATTTTCTTGAAGTAGTCCTCTTTCAGCTGCTTCACGGTCACTTTCCGGTTGGTCTTGAACAGCAGGTCCACGATGGCC

TTTTTCTGCTCGCCGCTCAGGAAGGCGGGCTTTCTCATTCCCTCGGTCACGTATTTCACTTTGGTCAGCTCGTTATACAC

GGTGAAGTACTCGTACAGCAGGCTGTGCTTGGGCAGCACCTTCTCGTTGGGCAGGTTCTTATCGAAGTTGGTCATCCGCT

CGATGAAGCTCTGGGCGGAAGCGCCCTTGTCCACCACTTCCTCGAAGTTCCAGGGGGTGATGGTTTCCTCGCTCTTTCTG

GTCATCCAGGCGAATCTGCTGTTTCCCCTGGCCAGAGGGCCCACGTAGTAGGGGATGCGGAAGGTCAGGATCTTCTCGAT

CTTTTCCCGGTTGTCCTTCAGGAATGGGTAAAAATCTTCCTGCCGCCGCAGAATGGCGTGCAGCTCTCCCAGGTGGATCT

GGTGGGGGATGCTGCCGTTGTCGAAGGTCCGCTGCTTCCGCAGCAGGTCCTCTCTGTTCAGCTTCACGAGCAGTTCCTCG

GTGCCGTCCATCTTTTCCAGGATGGGCTTGATGAACTTGTAGAACTCTTCCTGGCTGGCTCCGCCGTCAATGTAGCCGGC

GTAGCCGTTCTTGCTCTGGTCGAAGAAAATCTCTTTGTACTTCTCAGGCAGCTGCTGCCGCACGAGAGCTTTCAGCAGGG

TCAGGTCCTGGTGGTGCTCGTCGTATCTCTTGATCATAGAGGCGCTCAGGGGGGCCTTGGTGATCTCGGTGTTCACTCTC

AGGATGTCGCTCAGCAGGATGGCGTCGGACAGGTTCTTGGCGGCCAGAAACAGGTCGGCGTACTGGTCGCCGATCTGGGC

CAGCAGGTTGTCCAGGTCGTCGTCGTAGGTGTCCTTGCTCAGCTGCAGTTTGGCATCCTCGGCCAGGTCGAAGTTGCTCT

TGAAGTTGGGGGTCAGGCCCAGGCTCAGGGCAATCAGGTTTCCGAACAGGCCATTCTTCTTCTCGCCGGGCAGCTGGGCG

ATCAGATTTTCCAGCCGTCTGCTCTTGCTCAGTCTGGCAGACAGGATGGCCTTGGCGTCCACGCCGCTGGCGTTGATGGG

GTTTTCCTCGAACAGCTGGTTGTAGGTCTGCACCAGCTGGATGAACAGCTTGTCCACGTCGCTGTTGTCGGGGTTCAGGT

CGCCCTCGATCAGGAAGTGGCCCCGGAACTTGATCATGTGGGCCAGGGCCAGATAGATCAGCCGCAGGTCGGCCTTGTCG

GTGCTGTCCACCAGTTTCTTTCTCAGGTGGTAGATGGTGGGGTACTTCTCGTGGTAGGCCACCTCGTCCACGATGTTGCC

GAAGATGGGGTGCCGCTCGTGCTTCTTATCCTCTTCCACCAGGAAGGACTCTTCCAGTCTGTGGAAGAAGCTGTCGTCCA

CCTTGGCCATCTCGTTGCTGAAGATCTCTTGCAGATAGCAGATCCGGTTCTTCCGTCTGGTGTATCTTCTTCTGGCGGTT

CTCTTCAGCCGGGTGGCCTCGGCTGTTTCGCCGCTGTCGAACAGCAGGGCTCCGATCAGGTTCTTCTTGATGCTGTGCCG

GTCGGTGTTGCCCAGCACCTTGAATTTCTTGCTGGGCACCTTGTACTCGTCGGTGATCACGGCCCAGCCCACAGAGTTGG

TGCCGATGTCCAGGCCGATGCTGTACTTCTTGTCGGCTGCTGGGACTCCGTGGATACCGACCTTCCGCTTCTTCTTTGGG

GCCATCTTATCGTCATCGTCTTTGTAATCAATATCATGATCCTTGTAGTCTCCGTCGTGGTCCTTATAGTCCATCCTAGG

TGATATATTTCTATTAGGTATTTATTATTATAAAATATAAATCTTGAATGATAATAAATAAAATATTAGTTATTCCTTTT

CTAGTTTAAAATATACATATTATAAATATATATATATATATATATATTTTTATTGTGACAAGAATATATAATTATAAATT

ATATTATTTATTTTTGTATTTTTTTTTTTTTTTTTTTTTTTTTCTTTTTTTGTTTTATTTTTCTTTTTTTTTATAAATAT

TATTTTTTTCTTTTATCATGCACATTGGAATAATACATTAATATATATATATATATTATATTATACATATATTGAATAAT

GTTTATAAAAAATGCATAACTTATATGAATATAATTTTTTTTAAATATGACAAAAAGAAAAAAAAAAAAAACCAAAAAAA

ATTAAAATTGAAATGAAATATATAAATATATTATTTATATATATTATACATTGTTTAATACTACTACATGTATATATATA

TATTATATATATATATATATATCAATTTTTTCAAAAATAAATTAATATAAAAAGAGGGGAAAAAAAAAAAAAAAAAAAAA

AAAAGATAATTAAGTAAGCATTTAAAAATATATAAATTGATAATATATAAAATTAATCACATATAAACTAATATAATTTA

TAAAATAAGGAAAATAAAATATTACCATAAAATAAAAATAAAAATAAAAAAAAAAAAAAAAAACACCTTTTTTTATATAT

ATTAATATATAATTATCTCTTAGAAAAAATATTGTATAATTATATATGTAATGATTTATATAAAAAAATAAAATTATACA

AGTATATATTTTGTTTCTATAAATTGATATCTTAATTATTTATTATTAGAAATAGATATTTTTATAATAAACCAATAGAT

AAAATTTGTAGAGAAAAAAAAATAAAAATAAAAATAAAAATAATATAATATATAATAAAATAAAATAATATTATATAAAT

ATATTTTAATTTTTTTTACAAAATGGTTGGGGGCGCCACCGGTTATGTAATAATAATATGATCATAATATTATAATAAAA

CTTATAAAAAAAATATTAAATATTTCATAAATGATTATTATTTATAATAAGATAGATTCTTTAATTATTTTAAAATTGTA

TATTTTTTATGTATATTAATTTATTAATATTAATAAGAATATTATAAAAATTCTATTTTATTATTTAATGATATTACCCT

AATAAAAATAATATAATTATATTACAATATAAATTTATATATATATATATATTTATATATTTAAAGAATATTTTATTTTT

CAATAAGAACCTTCATTTTAAATTAACATCAAATTATATATATGTATATATACTTCTTAGTATTATTAATTAAAATACGG

AATAATATATAATATATATAAAATGGCAAAACTTTCCTATAGAAAAAAATATTCCATTTATTATATTTGTTGTAGGTAAT

TCTTATTACCGTTTCCTTCTGTTCGTAATGTATATTGGTATGTACTTTATTTTTGCAATTTAATTATATGTAAAAAAACG

TTAGTACACCATATATATATTATAGTTATAAGAATGCATGCCAAGCCTTTGTCTCAAGAAGAATCCACCCTCATTGAAAG

AGCAACGGCTACAATCAACAGCATCCCCATCTCTGAAGACTACAGCGTCGCCAGCGCAGCTCTCTCTAGCGACGGCCGCA

TCTTCACTGGTGTCAATGTATATCATTTTACTGGGGGACCTTGTGCAGAACTCGTGGTGCTGGGCACTGCTGCTGCTGCG

GCAGCTGGCAACCTGACTTGTATCGTCGCGATCGGAAATGAGAACAGGGGCATCTTGAGCCCCTGCGGACGGTGCCGACA

GGTGCTTCTCGATCTGCATCCTGGGATCAAAGCCATAGTGAAGGACAGTGATGGACAGCCGACGGCAGTTGGGATTCGTG

AATTGCTGCCCTCTGGTTATGTGTGGGAGGGCGCTAGCGGCAGTGGAGAGGGCAGAGGAAGTCTGCTAACATGCGGTGAC

GTCGAGGAGAATCCTGGCCCAAAGCTTATGGCACCAAAGGGTAGAAGTACAAATGAAATTGAACTTAGCGCAAGAGATGT

TTTGGAAAATATTGGAATAGGAATATATAATCAGGAAAAAATAAAAAAGAATCCATATGAACAACAATTGAAAGGCACAT

TATCAAACGCCCGATTTCATGATGGCTTGCACAAGGCAGCTGATTTGGGGGTAATACCTGGTCCTTCACATTTTTCTCAG

CTTTATTACAAAAAGCATACTAATAACACAAAATATTATAAGGATGATAGGCATCCTTGTCATGGTAGACAAGGAAAACG

TTTTGATGAAGGTCAAAAATTTGAATGTGGTAATGATAAAATAATTGGTAATAGCGATAAATATGGATCCTGTGCTCCAC

CTAGAAGAAGACATATATGTGATCAAAATTTAGAATTCTTAGATAACAATCATACTGATACTATTCATGATGTATTGGGA

AATGTGTTGGTCACAGCAAAATATGAAGGTGAATCTATTGTTAATGATCATCCAGATAAAAAGAACAATGGTAATAAATC

AGGTATATGTACTTCTCTTGCACGAAGTTTTGCCGATATAGGTGATATTGTAAGAGGAAGAGATATGTTTAAACCTAATG

ACAAAGATGCAGTGCGGCATGGTTTAAAGGTAGTTTTTAAGAAAATATATGATAAATTGTCACCTAAAGTACAAGAACAT

TACAAAGATGTTGATGGATCTGGAAATTACTATAAATTAAGGGAAGATTGGTGGACAGCGAACAGAGATCAAGTATGGAA

AGCCATAACATATGAAGCTCCACAGGATGCCAATTATTTTAGAAATGTTTCAGGAACAACTATGGCGTTTACAAGTGCAG

GAAAATGTAGACACAATGACAATAGCGTCCCAACGAATCTAGATTATGTCCCTCAATTTTTACGTTGGTACGATGAATGG

GCAGATGATTTTTGTCGAATAAGAAATCATAAGTTGCAAAAGGTTAAAGACACATGTACTAGTACCGGTACGCGTGACGT

CAGGTGGCACTTTTCGGGGAAATGTGCGCGGAACCCCTATTTGTTTATTTTTCTAAATACATTCAAATATGTATCCGCTC

ATGAGACAATAACCCTGATAAATGCTTCAATAATATTGAAAAAGGAAGAGTATGAGTATTCAACATTTCCGTGTCGCCCT

TATTCCCTTTTTTGCGGCATTTTGCCTTCCTGTTTTTGCTCACCCAGAAACGCTGGTGAAAGTAAAAGATGCTGAAGATC

AGTTGGGTGCACGAGTGGGTTACATCGAACTGGATCTCAACAGCGGTAAGATCCTTGAGAGTTTTCGCCCCGAAGAACGT

TTTCCAATGATGAGCACTTTTAAAGTTCTGCTATGTGGCGCGGTATTATCCCGTATTGACGCCGGGCAAGAGCAACTCGG

TCGCCGCATACACTATTCTCAGAATGACTTGGTTGAGTACTCACCAGTCACAGAAAAGCATCTTACGGATGGCATGACAG

TAAGAGAATTATGCAGTGCTGCCATAACCATGAGTGATAACACTGCGGCCAACTTACTTCTGACAACGATCGGAGGACCG

AAGGAGCTAACCGCTTTTTTGCACAACATGGGGGATCATGTAACTCGCCTTGATCGTTGGGAACCGGAGCTGAATGAAGC

CATACCAAACGACGAGCGTGACACCACGATGCCTGTAGCAATGCCAACAACGTTGCGCAAACTATTAACTGGCGAACTAC

TTACTCTAGCTTCCCGGCAACAATTAATAGACTGGATGGAGGCGGATAAAGTTGCAGGACCACTTCTGCGCTCGGCCCTT

CCGGCTGGCTGGTTTATTGCTGATAAATCTGGAGCCGGTGAGCGTGGGTCTCGCGGTATCATTGCAGCACTGGGGCCAGA

TGGTAAGCCCTCCCGTATCGTAGTTATCTACACGACGGGGAGTCAGGCAACTATGGATGAACGAAATAGACAGATCGCTG

AGATAGGTGCCTCACTGATTAAGCATTGGTAACTGTCAGACCAAGTTTACTCATATATACTTTAGATTGATTTAAAACTT

CATTTTTAATTTAAAAGGATCTAGGTGAAGATCCTTTTTGATAATCTCATGACCAAAATCCCTTAACGTGAGTTTTCGTT

CCACTGAGCGTCAGACCCCGTAGAAAAGATCAAAGGATCTTCTTGAGATCCTTTTTTTCTGCGCGTAATCTGCTGCTTGC

AAACAAAAAAACCACCGCTACCAGCGGTGGTTTGTTTGCCGGATCAAGAGCTACCAACTCTTTTTCCGAAGGTAACTGGC

TTCAGCAGAGCGCAGATACCAAATACTGTCCTTCTAGTGTAGCCGTAGTTAGGCCACCACTTCAAGAACTCTGTAGCACC

GCCTACATACCTCGCTCTGCTAATCCTGTTACCAGTGGCTGCTGCCAGTGGCGATAAGTCGTGTCTTACCGGGTTGGACT

CAAGACGATAGTTACCGGATAAGGCGCAGCGGTCGGGCTGAACGGGGGGTTCGTGCACACAGCCCAGCTTGGAGCGAACG

ACCTACACCGAACTGAGATACCTACAGCGTGAGCTATGAGAAAGCGCCACGCTTCCCGAAGGGAGAAAGGCGGACAGGTA

TCCGGTAAGCGGCAGGGTCGGAACAGGAGAGCGCACGAGGGAGCTTCCAGGGGGAAACGCCTGGTATCTTTATAGTCCTG

TCGGGTTTCGCCACCTCTGACTTGAGCGTCGATTTTTGTGATGCTCGTCAGGGGGGCGGAGCCTATCGAAAAACGCCAGC

AACGCGGCCTTTTTACGGTTCCTGGCCTTTTGCTGGCCTTTTGCTCACATGTTCTTTCCTGCGTTATCCCCTGATTCTGT

GGATAACCGTATTACCGCCTTTGAGTGAGCTGATACCGCTCGCCGCAGCCGAACGACCGAGCGCAGCGAGTCAGTGAGCG

AGGAAGCGGAAGAGCGCCCAATACGCAAACCGCCTCTCCCCGCGCGTTGGCCGATTCATTAATGCAGCTGGCACGACAGG

TTTCCCGACTGGAAAGCGGGCAGTGAGCGCAACGCAATTAATGTGAGTTAGCTCACTCATTAGGCACCCCAGGCTTTACA

CTTTATGCTTCCGGCTCGTATGTTGTGTGGAATTGTGAGCGGATAACAATTTCACACAGGAAACAGCTATGACCATGATT

ACGCCAAGCTATTTAGGTGACACTATAGAATACTC
